# Biosynthesis of gibberellin-related compounds modulates far-red light responses in the liverwort *Marchantia polymorpha*

**DOI:** 10.1101/2023.05.04.539458

**Authors:** Rui Sun, Maiko Okabe, Sho Miyazaki, Toshiaki Ishida, Kiyoshi Mashiguchi, Keisuke Inoue, Yoshihiro Yoshitake, Shohei Yamaoka, Ryuichi Nishihama, Hiroshi Kawaide, Masatoshi Nakajima, Shinjiro Yamaguchi, Takayuki Kohchi

## Abstract

The phytohormone gibberellins (GAs) are key regulators of growth, development and environmental responses in angiosperms. From an evolutionary perspective, all major steps of GA biosynthesis are conserved among vascular plants, while GA biosynthetic intermediates such as *ent*-kaurenoic acid (KA) are also produced by bryophytes. Here we show that in the liverwort *Marchantia polymorpha*, KA and GA_12_ are synthesized by evolutionarily conserved enzymes, which are required for developmental responses to far-red light (FR). Under FR-enriched conditions, mutants of various biosynthesis enzymes consistently altered thallus growth allometry, delayed the initiation of gametogenesis, and affected the morphology of gamete-bearing structures (gametangiophores). By chemical treatments and LC-MS/MS analyses, we confirmed these phenotypes were caused by deficiency of some GA-related compounds derived from KA, but not bioactive GAs from vascular plants. Transcriptome analysis showed that FR enrichment induced the up-regulation of genes related to stress responses and secondary metabolism in *M. polymorpha*, which was largely dependent on the biosynthesis of GA-related compounds. Due to the lack of the canonical GA receptors in bryophytes, we hypothesize that GA-related compounds are commonly synthesized in land plants but co-opted independently to regulate responses to light quality change in different lineages during the past 450 million years of evolution.

## INTRODUCTION

Throughout evolution, plants developed various chemical tools to optimize growth and development, and to cope with environmental changes. Gibberellins (GAs) are a group of tetracyclic diterpenoid compounds broadly produced by many plants and plant-associated microbes. Among the more than 130 identified GAs, a few of them (GA_1_, GA_3_, GA_4_, and GA_7_) are considered as commonly bioactive in angiosperms (reviewed in Sponsel, 2016), stimulating seed germination (Koornneef and van der Veen, 1980; Toyomasu et al., 1998; Yamaguchi et al., 1998a; Ogawa et al., 2003; Gabriele et al., 2009) and promoting growth of various plant organs (Kurosawa, 1926; Koornneef and van der Veen, 1980; Wenzel et al., 2000; Ubeda-Tomás et al., 2008, 2009; Achard et al., 2009; Nelissen et al., 2012).

In *Arabidopsis thaliana*, the biosynthesis of GAs starts with the production of *ent-*kaurene from geranylgeranyl diphosphate (GGDP) by two terpene synthases (TPSs), *ent*-copalyl diphosphate synthase (CPS) and *ent-*kaurene synthase (KS) (Sun and Kamiya, 1994; Yamaguchi et al., 1998b). Next, *ent-*kaurene is oxidized by two cytochrome P450 monooxygenases (CYPs), first into *ent-*kaurenoic acid (KA) by *ent-*kaurene oxidase (KO), then into GA_12_ by *ent-*kaurenoic acid oxidase (KAO) (Helliwell et al., 1998, 1999, 2001a, 2001b). Finally, GA_12_ is converted into the bioactive form GA_4_ through sequential oxidation by two 2-oxoglutarate-dependent dioxygenases (2-OGDs), GA 20-oxidase (GA20ox) and GA 3-oxidase (GA3ox) (Chiang et al., 1995; Yamaguchi et al., 1998a; Williams et al., 1998). A second bioactive GA, GA_1_, is different from GA_4_ by one additional hydroxyl group on C-13, which is possibly introduced to GA_12_ or KA before the subsequent oxidation steps (Talon et al., 1990; Nomura et al., 2013) (Supplemental Figure 1).

Bioinformatic and chemical analyses revealed that this biosynthetic pathway producing GA_4_ and/or GA_1_ is conserved among vascular plants (Hirano et al., 2007; Tanaka et al., 2014; Cannell et al., 2020). In several fern species, GA-derived compounds modulate sexual differentiation of gametophytes and spore germination in the darkness (Yamane, 1998; Schneller, 2008; Tanaka et al., 2014; Hornych et al., 2021). Bryophytes have no 2-OGD-family members of GA biosynthesis enzymes, thus considered as lacking the production of common bioactive GAs. However, homologs for CPS, KS, KO and KAO still exist in bryophytes, which suggested an ancestral capacity to synthesize the GA precursors KA and GA_12_ in all land plants (Cannell et al., 2020). In the moss *Physcomitrium patens* which lacks KAO, KA is synthesized by a bifunctional CPS/KS and a single KO homolog (Hayashi et al., 2006, 2010; Miyazaki et al., 2011, 2015). The KA derivative, *ent-*3β-OH-KA, is known as a bioactive molecule to regulate protonema differentiation and blue light avoidance in *P. patens* (Hayashi et al., 2010; Miyazaki et al., 2014, 2015, 2018).

In the liverwort *Marchantia polymorpha*, homologs for CPS, KS, KO and KAO were identified in genome-wide analyses (Kumar et al., 2016; Bowman et al., 2017). Also, MpCPS/DTPS3 and MpKS/DTPS4 (hereafter referred to as MpCPS and MpKS) have been reported to catalyze the production of *ent*-copalyl diphosphate and *ent-*kaurene (Kumar et al., 2016; Jia et al., 2022). In transcriptome analysis, upregulation of GA biosynthesis gene homologs was observed under far-red light (FR) enriched conditions, suggesting a role for this pathway in the response to light quality change (Briginshaw et al., 2022). However, no empirical knowledge has been established yet about the exact physiological function of GA-related compounds in liverworts.

FR enrichment mimics the proximity of competitive neighbors in the nature habitat. In many angiosperms, this is perceived by phytochrome (phy) photoreceptors and triggers shade-avoiding responses, including elongation of stem-like structures, hyponastic growth of petioles and acceleration of flowering (Downs et al., 1957; Holmes and Smith, 1975; Morgan and Smith, 1978, 1979; Whitelam and Johnson, 1982; Whitelam and Smith, 1991). GA biosynthesis is often evoked in this process, and is required for the induction of elongative growth (García-Martínez et al., 1987; Beall et al., 1996; Van Tuinen et al., 1999; Hisamatsu et al., 2005; Djakovic-Petrovic et al., 2007; Dubois et al., 2010). For example, in *A. thaliana*, a local FR enrichment at the leaf tip induced the expression of GA biosynthesis genes in both the leaf tip and the petiole, which modulates the hyponastic growth of the petiole (Sessa et al., 2005; Hisamatsu et al., 2005; Djakovic-Petrovic et al., 2007; Bou-Torrent et al., 2014; Kohnen et al., 2016; Küpers et al., 2023). In the gymnosperm *Pinus tabuliformis*, FR-induced shoot elongation was also accompanied with and dependent on the accumulation of bioactive GAs (Li et al., 2020).

As a liverwort, the life cycle of *M. polymorpha* is dominated by the thalloid gametophyte. End-of-day FR irradiation is known to cause hyponastic growth of thallus tips and decrease in chlorophyll (Fredericq, 1964; Ninnemann and Halbsguth, 1965; Fredericq and de Greef, 1966; Fredericq and Greef, 1968). FR enrichment also induced hyponastic thallus growth (Briginshaw et al., 2022), accompanied by the growth activity change of apical meristems and the transition to sexual reproduction (Chiyoda et al., 2008; Inoue et al., 2019; Streubel et al., 2023). Similar to *A. thaliana* and other land plants, these responses in *M. polymorpha* are mediated by the sole phytochrome (Mpphy) and the single-copy transcription factor PHYTOCHROME-INTERACTING FACTOR (MpPIF) (Fredericq, 1964; Inoue et al., 2016, 2019; Streubel et al., 2023). In this study, we characterized evolutionarily conserved GA biosynthesis enzymes in *M. polymorpha* with genetic approach, showing that they were indispensable for developmental and gene expression responses to FR enrichment.

## RESULTS

### Mp*CPS* is required for thallus morphological changes induced by FR enrichment

To explore the function of GA-related hormones in *M. polymorpha*, we used the CRISPR/Cas9^D10A^ nickase system (Hisanaga et al., 2019; Koide et al., 2020) to create large-deletion mutants of Mp*CPS* (Mp2g07200), which encodes the first enzyme of the biosynthesis pathway. Two different mutant alleles (Mp*cps-4^ld^* and Mp*cps-27^ld^*) with complete loss of the coding sequence (CDS) were isolated from Tak-1, a male wild-type accession (Supplemental Figure 2A). Mp*cps-4^ld^* was further complemented by expressing the CDS of Mp*CPS* with C-terminal Citrine fusion under the control of a cauliflower mosaic virus 35S promoter (*_pro_35S:*Mp*CPS-Cit*), generating two independent transgenic lines. First, we observed the thallus morphology of 12-day-old plants grown from gemmae under two different light conditions, either the continuous white light (cW), or cW supplemented with continuous far-red light (cW+cFR). FR enrichment under cW+cFR induced morphological changes in Tak-1 wild-type plants, marked by increased growth angles (hyponasty) and slenderer thallus shapes, the latter shown as the increase of length-width ratio measured from half of the thallus (Figure 1A-C). Such an increase was not observed in Mp*cps^ld^* mutants, suggesting a role for Mp*CPS* in modulating FR-induced growth responses. Furthermore, Mp*cps^ld^*mutants were significantly larger in thallus size than wild-type plants, particularly under cW+cFR conditions (Figure 1A,D). It is likely that MpCPS acts in a pathway producing compounds inhibiting growth, rather than promoting growth like bioactive GAs in angiosperms. The complementation lines carrying *_pro_35S:*Mp*CPS-Cit* recovered the hyponastic growth, the thallus shape and the thallus size of Mp*cps-4^ld^*, confirming that the phenotypes were caused by Mp*CPS* loss-of-function (Figure 1A-D). As a further verification, Mp*cps^ld^* mutant alleles and complementation lines were constructed in the female wild-type accession (Tak-2). Similarly, Mp*cps^ld^* mutations resulted in decrease of thallus hyponasty and the length-width ratio, but increased the thallus area drastically under cW+cFR (Supplemental Figure 3).

**Figure 1.**
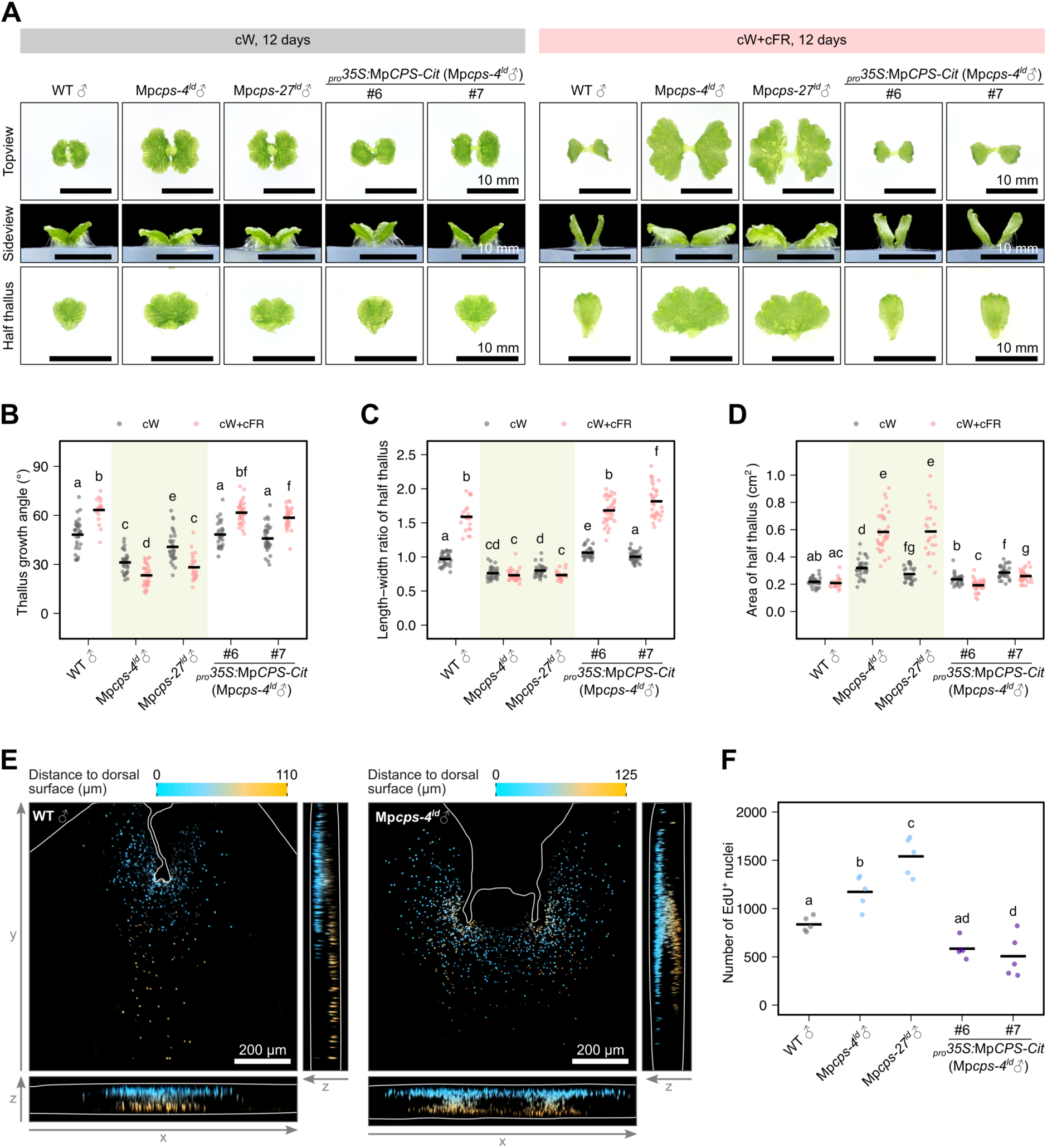
Loss of Mp*CPS* activity alters thallus morphology under far-red light (FR) enriched conditions. A, Morphology of 12-day-old plants grown from gemmae under continuous white light (cW) or continuous white light with far-red light (cW+cFR) conditions. Images were selected randomly from different plants. Bars = 10 mm. B-D, Measurements of the thallus growth angle (B), length-width ratio (C) and thallus area (D) from half plants shown in A. n=21-36. E, Apical regions of plants labelled with 5-ethynyl-2’-deoxyuridine (EdU) after 7-day growth in cW+cFR, showing two-dimensional projections of color-coded z-stacks. White lines mark the boundary of plants or imaging area. F, Number of nuclei with positive EdU signals in the apical regions of 7-day-old plants grown in cW+cFR (n=5). For all figures, WT♂ refer to Tak-1 wild-type plants. For B-D and F, each dot represents data from a “half thallus”, which was developed from a single meristem of the gemma. Horizontal lines represent mean values. Letters represent statistical differences, and groups with no common letters were significantly different (adjusted *p*<0.05). For B-D, non-pooled Welch’s *t*-test with Benjamini-Hochberg (B-H) adjustment was used due to heterogeneity of variance. Tukey’s HSD test was used in F.

To examine the influence of Mp*cps^ld^*on growth activity at the tissue level, we labelled 7-day-old plants grown under cW+cFR with 5-ethynyl-2’-deoxyuridine (EdU), which could be incorporated into actively dividing cells during DNA synthesis. In the wild-type plants, we observed a relatively narrow distribution of EdU-positive nuclei in the slender thallus, dispersing from the apical meristem to the basal region along the midrib. By assigning different pseudo-colors to signals acquired at different depths by confocal microscopy, we found different distributions of dorsal and ventral signals in these plants. Ventral EdU signals along the midrib extended further in the basal direction, suggesting excessive cell divisions specific to the ventral side, which might contribute to the hyponastic growth induced by FR enrichment (Figure 1E). While in Mp*cps^ld^* mutants, EdU signals indicated active cell divisions in a broader range. Two fully separated apical meristems were usually seen from the observed region, and the total numbers of EdU-positive nuclei were significantly higher than those of wild-type plants or complementation lines (Figure 1E-F). In addition, dorsal and ventral EdU signals showed similar distribution ranges in Mp*cps^ld^* mutants, which was consistent with their flat morphology under cW+cFR (Figure 1E).

### Mp*CPS* has a role in modulating gametangiophore development

Continuous FR irradiation is known to induce sexual reproduction in *M. polymorpha*, i.e. the formation of sexual branches called gametangiophores, and the differentiation of sexual organs called gametangia (Yamaoka et al., 2018; Inoue et al., 2019). To observe this process, plants were grown from gemmae under cW for 7 days, then transferred to cW+cFR for gametangiophore induction. As a liverwort, *M. polymorpha* undergoes vegetative growth with dichotomous branching, periodically multiplying the number of apical meristems through bifurcation. Generally, all apical meristems remain indeterminate under white light conditions, while a proportion of them become dormant or differentiate into gametangiophores under FR-enriched light conditions (Streubel et al., 2023). During the induction under cW+cFR, we observed higher numbers of total apical meristems in Mp*cps^ld^* mutants than in wild-type or complementation lines, which is in line with their more active vegetative growth (Figure 2A-B; Supplemental Figure 4A). If the potential for apical meristems to form gametangiophores was similar between the mutants and wild-type plants, higher number of gametangiophore-bearing apexes would be expected in Mp*cps^ld^* mutants. However, Mp*cps^ld^*mutants formed fewer gametangiophores than wild-type plants and complementation lines during the first 16 days after FR irradiation, suggesting that gametangiophore formation is inhibited in these mutants (Figure 2A-B; Supplemental Figure 4A).

**Figure 2.**
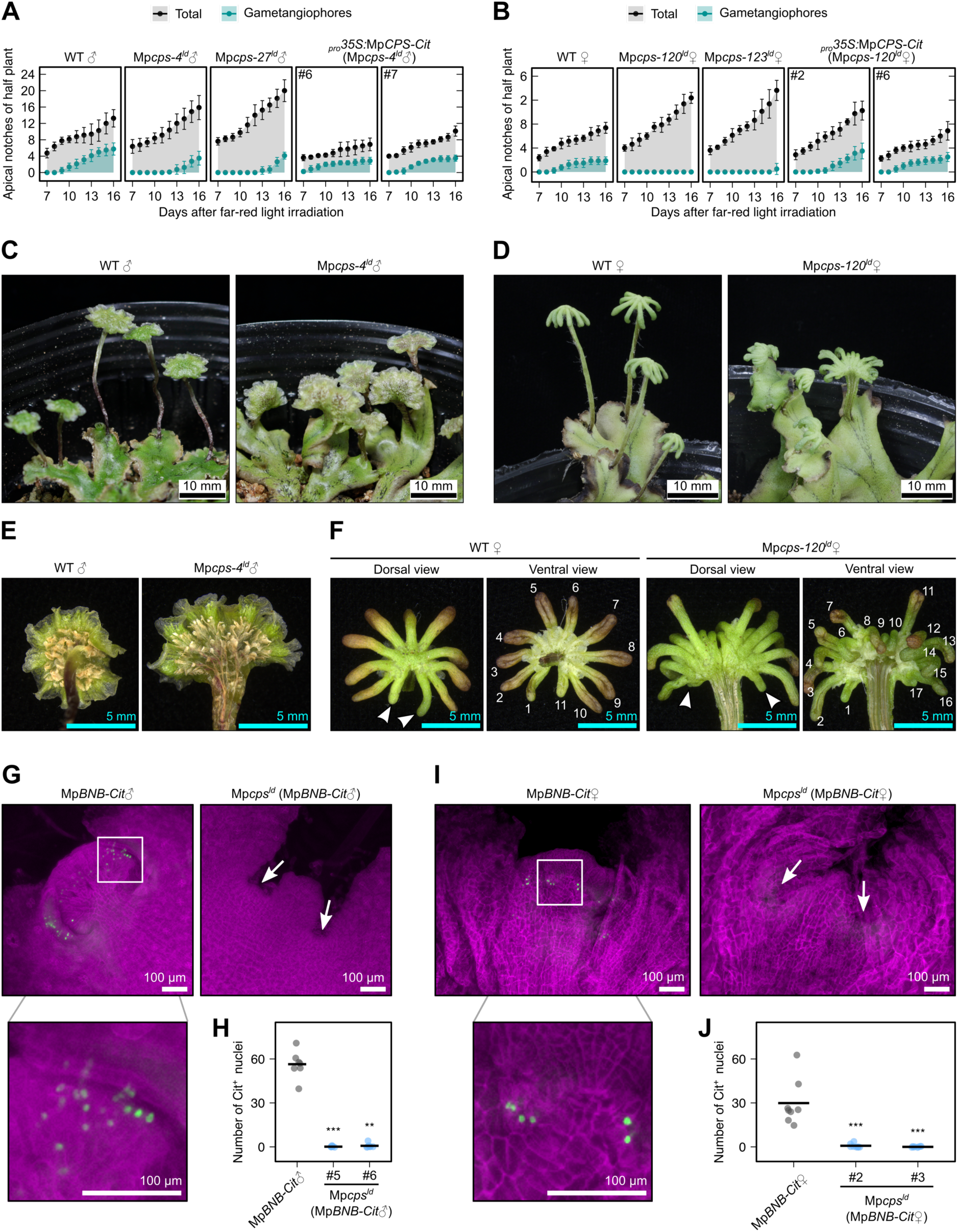
Mp*CPS* is required for delayed sexual reproduction and affected gametangiophore morphology. A-B, Progress of apical bifurcation and gametangiophore formation in male (A) and female (B) plants, which were cultured aseptically under cW for 7 days before transferred to cW+cFR. Dots and error bars represent means and standard deviations, respectively. n=8. WT♂ and WT♀ refer to Tak-1 and Tak-2 wild-type plants, respectively. C-D, Photos of male (C) and female (D) plants bearing gametangiophores, cultured on vermiculite under cW+cFR from thallus fragments for 81 and 63 days, respectively. E, Ventral view of antheridiophores. F, Dorsal and ventral view of archegoniophores. Arrowheads indicate marginal digitate rays, and the number of digitate rays was labeled in the ventral view. G-J, Fluorescence microscopic images and quantification of MpBNB-Cit accumulation in the apical regions of 11-day-old male (G-H, dorsal view), or 14-day-old female (I-J, ventral view) plants cultured under cW+cFR from gemmae. G and I, Z-projections of image stacks, with cell walls stained with calcofluor white (purple) and Citrine signals shown in green. Arrows indicate apical meristems. H and J, Number of Citrine-positive nuclei counted from projection of image stacks. Each dot represents data from 1/2 (H) or 1/4 (J) of the thallus, and horizontal lines represent mean values. Asterisks show statistical difference compared to the control group (Mann-Whitney *U* test; **, *p*<0.01; ***, *p*<0.001). n=7 for H, n=8 for J.

In addition, the morphology of gametangiophores was distorted in Mp*cps^ld^* mutants. As *M. polymorpha* is a dioicous species, gametangiophores are sexually dimorphic among plants with different sex chromosomes. The male gametangiophores (antheridiophores) of wild-type plants had roundish, disc-like receptacles, which were ventrally connected to a relatively long and thin stalk at the center. While for Mp*cps^ld^* mutants, the male receptacles were fan-shaped, attaching to a short and thick stalk(s) at the basal end (Fig 2C,E; Supplemental Figure 4B). Transverse sectioning revealed thallus-like features in the stalks of Mp*cps^ld^* antheridiophores. Usually, no air chambers could be observed in the near-cylindric antheridiophore stalks of wild-type plants. In contrast, Mp*cps^ld^* antheridiophore stalks had a flat dorsal surface beneath which air chambers with photosynthetic filaments were clearly formed (Supplemental Figure 5A), quite similar to the tissue organization in the vegetative thallus (Shimamura, 2016). Besides, Mp*cps^ld^* antheridiophore stalks frequently have more canals with pegged rhizoids in the ventral side, possibly reflecting additional bifurcation events during gametangiophore morphogenesis (Supplemental Figure 5A).

The female gametangiophores (archegoniophores) of wild-type plants were also stalked and had 9-11 finger-like structure (digitate rays) in the receptacle, which radially arranged like umbrella ribs. Two marginal digitate rays could be recognized as no involucres were produced between them (Figure 2D,F; Supplemental Figure 4B) (Cao et al., 2013). The female receptacles of Mp*cps^ld^* were palm-like, positioning the two marginal rays at opposite ends. Excessive number of digitate rays and/or bifurcation in the stalk were often observed in late-stage receptacles (Figure 2D,F; Supplemental Figure 4B). In extreme cases from aseptic culture, the archegoniophore of Mp*cps^ld^* remained the form of bifurcated thalloid branches, with digitate rays formed at the thallus tips (Supplemental Figure 4C). If the relatively fixed number of digitate rays in wild-type receptacles represents a determinate fate for the apical meristem, such excessive bifurcation or additional digitate rays in Mp*cps^ld^* mutants might suggest a loose transition from the indeterminate vegetative growth. Similar to the male case, the stalks of Mp*cps^ld^*archegoniophores were shorter and thicker than the wild-type counterparts, bearing more rhizoid canals in the ventral side (Figure 2D; Supplemental Figure 5C).

Since gametangium differentiation accompanies the morphogenesis of gametangiophores, we investigated this progress by examining Citrine-labelled MpBONOBO proteins (MpBNB-Cit), which specifically accumulate in the initial cells and immature gametangia (Yamaoka et al., 2018). To keep the consistency in genetic background, large-deletions of Mp*CPS* were introduced into Mp*BNB-Cit* knock-in lines through thallus transformation (Supplemental Figure 2), again using the CRISPR/Cas9^D10A^ nickase system (Hisanaga et al., 2019). As expected, the gametangiophore morphogenesis was delayed by Mp*cps^ld^* mutation. After 11 or 14 days of growth under cW+cFR, dome-shaped gametangiophore primordia already formed, respectively, in male and female Mp*BNB-Cit* plants carrying the wild-type Mp*CPS* allele. Under fluorescence microscopes, arrays and/or clusters of Citrine signals appeared at the edge of primordia, indicating on-going gametangium differentiation in these plants. In contrast, no gametangiophore primordium was yet visible in Mp*cps^ld^* Mp*BNB-Cit* lines of the same ages. Few or no Citrine-positive nuclei could be found in the apical regions of these plants, suggesting that Mp*cps^ld^* also caused a delay in gametangium differentiation (Figure 2G-J). Despite such delay, male and female gametangia of normal morphology eventually formed in Mp*cps^ld^* mutants, as shown by the longitudinal sections of the receptacles (Supplemental Figure 5B,D). Crossing experiments further confirmed the fertility of gametes, as mature spores could be produced from all combinations among Mp*cps^ld^* and wild-type plants (Supplemental Figure 5E-F).

### Mp*cps^ld^* phenotypes can be rescued by *ent-*kaurenoic acid (KA)

To explore the GA biosynthesis pathway possibly blocked by Mp*cps^ld^*, we first investigated if common GAs from angiosperms could be detected endogenously in *M. polymorpha*. The plants were cultured for 10 days under cW, then 4 days under cW+cFR before harvested for analysis with liquid chromatography-tandem mass spectrometry (LC-MS/MS). In the Tak-1 wild-type plants, GA_12_ but not any downstream compounds from the angiosperm GA biosynthesis pathway could be detected (Figure 3A-B; Supplemental Figures 1 and 6A). The endogenous level of GA_12_ is 28.9 ± 6.5 pg/g fresh weight on average, which is much lower than the levels in the seedlings of *Arabidopsis thaliana* (Nomura et al., 2013). In Mp*cps-4^ld^* plants cultured under the same conditions, endogenous GA_12_ did not reach the detection limit, which supported the loss of GA biosynthesis in this mutant (Figure 3B).

**Figure 3.**
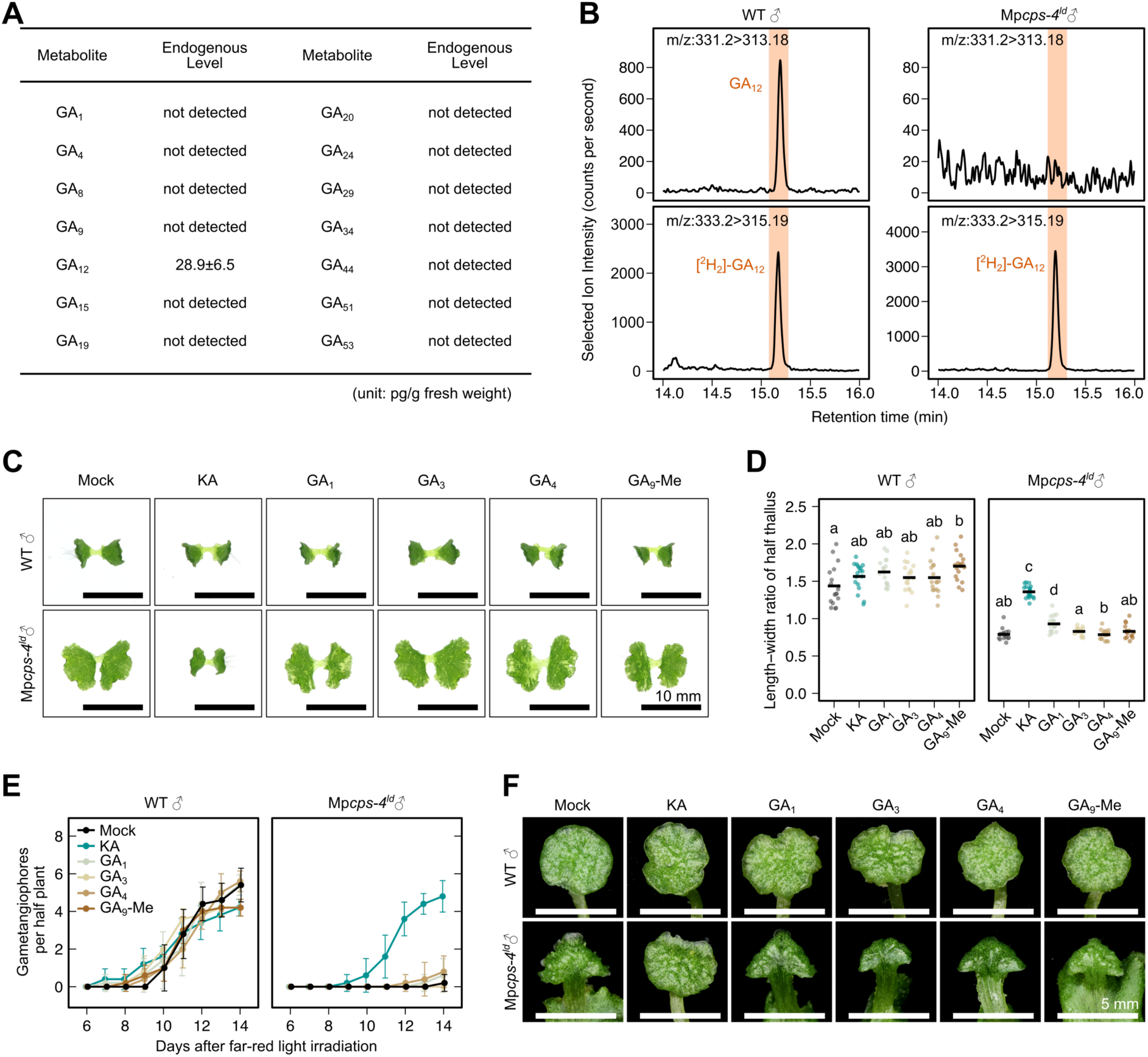
Endogenous levels of and responsiveness to KA or GAs in wild-type and Mp*cps^ld^* plants. A, Endogenous levels of GAs measured in wild-type plants (Tak-1) by LC-MS/MS. B, Selected ion chromatography showing peak of endogenous GA_12_ in comparison with the ^2^H_2_-labeled internal standard. For A-B, plants were cultured under cW for 10 days, then induced under cW+cFR for 3 days. C-D, Effect of 2-μM KA or GAs on the morphology of 12-day-old thalli cultured under cW+cFR from gemmae. Horizontal bars in D represent mean values, and letters represent multiple comparisons with non-pooled Welch’s *t*-test and B-H adjustment (adjusted *p*<0.05 for non-overlapping letters, n=16-18). E-F, Effect of 2-μM KA or GAs on gametangiophore formation (E) and morphology (F). Plants were cultured aseptically under cW for 7 days before transferred to cW+cFR. Dots and error bars represent mean±SD in E (n=5). Bars = 10 mm in C and 5 mm in F.

Next, we tested the effect of GA-related compounds on *M. polymorpha*. For thallus morphology observation, gemmae were planted on agar medium containing different GAs or the solvent control, then cultured under cW+cFR for 12 days. For gametangiophore induction, plants were transferred onto agar medium containing chemicals at the onset of cW+cFR induction. Several bioactive GAs in vascular plants and the GA biosynthesis precursor, KA, were tested on wild-type and Mp*cps-4^ld^* mutants. GA_12_ was not included in the assay due to its limited availability. As a result, 2-μM KA application fully complemented Mp*cps-4^ld^* phenotypes. The thallus shape, the progress of gametangiophore formation and the gametangiophore morphology were all restored to the manner of wild-type plants (Figure 3C-F). Furthermore, KA treatment altered the thallus morphology of Mp*cps-4^ld^*in a dose-dependent manner under cW+cFR. 100-nM KA was sufficient to induce clear changes in the size, shape and hyponasty of the mutant thallus (Supplemental Figure 7A, C-E). However, KA did not evidently change the thallus shape or gametangiophore formation in wild-type plants, which suggested it to be a biosynthetic intermediate rather than being directly bioactive. No active GAs in angiosperms (GA_1_, GA_3_ or GA_4_) rescued Mp*cps-4^ld^* as efficiently as KA, which was consistent with their absence in *M. polymorpha*. GA_9_ methyl ester (GA_9_-Me), which is released by several ferns as a pheromone and could rescue Pp*cps/ks* phenotype in the moss *P. patens* (Yamauchi et al., 1996; Tanaka et al., 2014; Hayashi et al., 2010), did not work on Mp*cps-4^ld^* in *M. polymorpha* (Figure 3C-F).

Taken together, these data indicated that Mp*cps^ld^*phenotypes were likely caused by the deficiency in some GA-related diterpenoid compound(s), which is synthesized from KA but different from bioactive GAs in vascular plants. For convenience, hereafter we refer to this putative bioactive compound as GA_Mp_.

### MpKOL1 and MpKAOL1 catalyze the biosynthesis of KA and GA_12_, respectively

The enzymatic activities for MpCPS to produce *ent*-copalyl diphosphate and MpKS (Mp6g05950) to produce *ent*-kaurene have been biochemically confirmed in the previous research (Kumar et al., 2016). To find the downstream enzymes catalyzing KA and GA_12_ biosynthesis in *M. polymorpha*, we tested the enzymatic activity of KO and KAO homologs (Figure 4A). Three KO homologs were identified by phylogenetic analysis in *M. polymorpha* (Supplemental Figure 8), and each of them was expressed in the methylotrophic yeast *Pichia pastoris* together with the *Arabidopsis* CYTOCHROME P450 REDUCTASE 1 (AtCPR1, AT4G24520). The yeasts were co-cultured with the substrate (*ent*-kaurene) for two days, then the culture infiltrates were extracted and analyzed with gas chromatography-mass spectrometry (GC-MS). KA was generated as a major product in the yeast culture expressing MpKOL1 (Mp3g18320), but not detected in cultures expressing MpKOL2 (Mp2g01950) or MpKOL3 (Mp2g01940) (Figure 4B; Supplemental Figure 6B).

**Figure 4.**
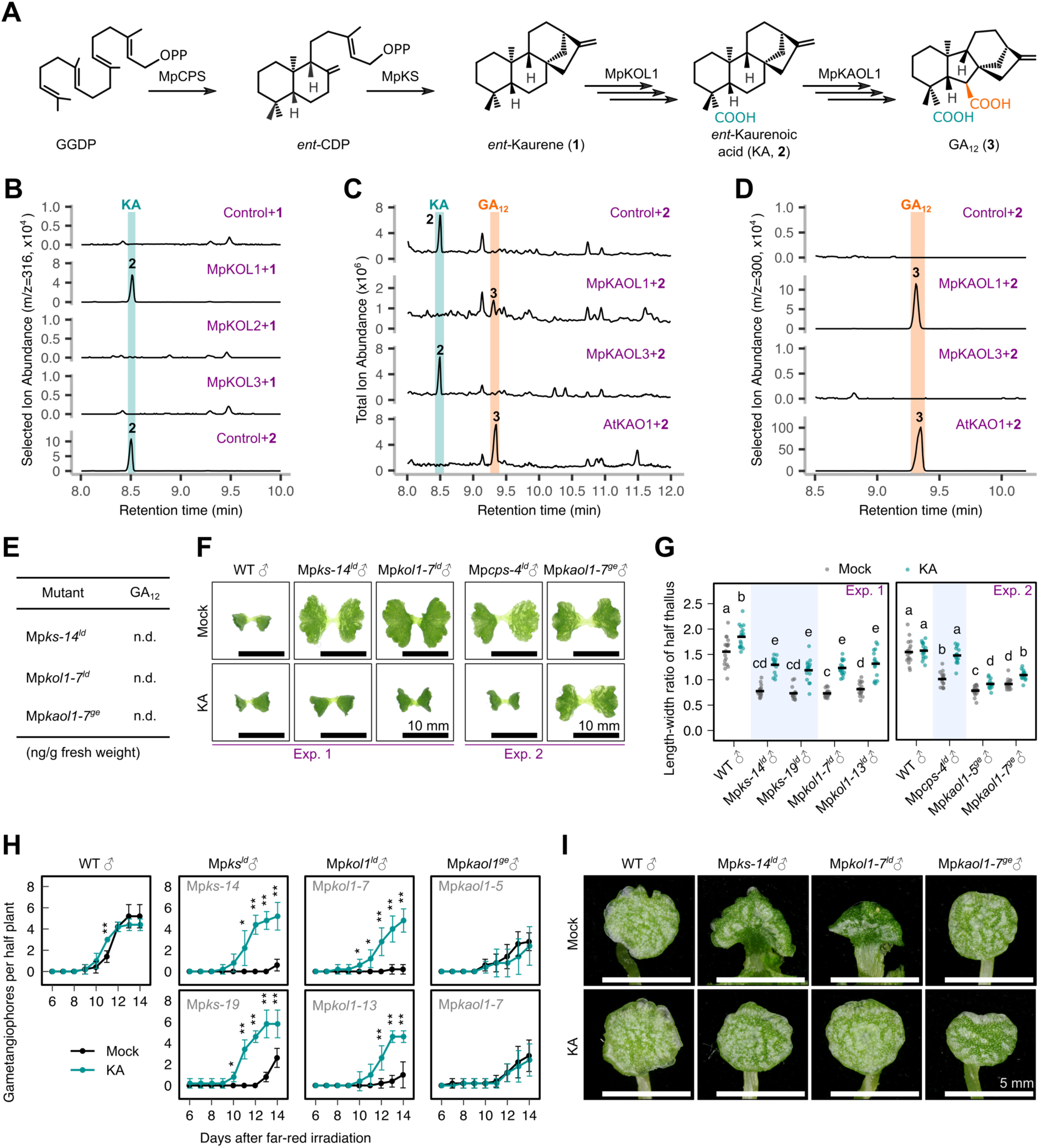
Biosynthetic route for GA_12_ production in *M. polymorpha*. A, Proposed enzymatic steps for GA_12_ biosynthesis in *M. poymorpha*. B-D, GC-MS analysis showing conversion of *ent*-kaurene to KA (B), and KA to GA_12_ (C-D) by yeast cultures expressing KO and KAO homologs. E, biosynthesis mutants with GA_12_ below the detection limit. n.d., not detected. F-G, morphlogy of 12-day-old mutants grown under cW+cFR with or without 2 μM KA. Horizontal bars in G represent mean values, and letters represent multiple comparisons with non-pooled Welch’s *t*-test and B-H adjustment (adjusted *p*<0.05 for non-overlapping letters, n=14-18). H-I, Gametangiophore formation progress (H) and morphology (I) of mutants, cultured aseptically under cW for 7 days before transferred to cW+cFR and treated with 2 μM KA. Dots and error bars represent mean±SD in H (n=5). Bars = 10 mm in F and 5 mm in I.

All currently known plant-type KAOs belong to the CYP88 family, in which two *M. polymorpha* members (MpKAOL1, Mp4g23680; and MpKAOL3, Mp2g10420) were confirmed by our phylogenetic analysis. Previous analysis considered the protein encoded by Mp1g25410 as a KAO homolog and named it MpKAOL2 (Bowman et al., 2017). In our current analysis, this protein and its liverwort homologs were closely related to CYP729 family members, which are distinctively different from CYP88 proteins (Supplemental Figures 9-10). After two day’s culturing, *Pichia* cells expressing MpKAOL1 showed a clear consumption of KA and production of GA_12_, which displayed identical retention time and mass spectra with the major product from the AtKAO1-expressing culture, i.e. the positive control. While in the MpKAOL3-expressing culture, consumption of KA was limited and no GA_12_ production was detected (Figure 4C-D, Supplemental Figure 6B). It seems that MpKOL1 and MpKAOL1, but not their *M. polymorpha* homologs, harbor catalytic activities similar to angiosperm GA biosynthesis enzymes.

By expressing Citrine-fused proteins under the control of the 35S promoter, we observed the subcellular localization of the four *M. polymorpha* enzymes which showed catalytic activities related to GA biosynthesis (Supplemental Figure 11). MpCPS-Cit and MpKS-Cit proteins were localized in the chloroplasts, most likely in the stroma as the Citrine signal intensities displayed complementary patterns to the thylakoid-enriched chlorophyll (Supplemental Figure 11A-B). The signals of MpKOL1-Cit proteins were also associated with chloroplasts, being strongest in chloroplast envelopes (Supplemental Figure 11C). On the other hand, MpKAOL1-Cit proteins seemed to aggregate in the endomembrane system and could be observed near the nuclear envelope and the plasma membrane (Supplemental Figure 11D). Overall, the subcellular distribution of these proteins were similar to their homologs in *A. thaliana* (Sun and Kamiya, 1994; Helliwell et al., 2001b), supporting them as being evolutionary conserved in land plants.

### KA is a pivotal intermediate in the biosynthesis of GA-related hormone

To validate the physiological role of GA biosynthesis enzymes other than MpCPS *in vivo*, we created loss-of-function mutants for all KS, KO and KAO homologs in *M. polymorpha*, using the CRISPR/Cas9^D10A^ nickase or the CRISPR/Cas9 system (Supplemental Figure 2B-D; Supplemental Figure 12). Consistent with the enzymatic activities in yeast or *in vitro* (Kumar et al., 2016), Mp*ks-14^ld^*, Mp*kol1-7^ld^*, and Mp*kaol1-7^ge^* mutants completely lost the ability to produce GA_12_ endogenously (Figure 4E; Supplemental Figure 6C-D). Under cW+cFR conditions, the phenotypes of Mp*ks^ld^* and Mp*kol1^ld^* mutants were similar to that of Mp*cps^ld^*. The thalli of these mutants were significantly larger, wider and flatter than those of wild-type plants (Figure 4F-G). Mp*ks^ld^* and Mp*kol1^ld^* mutants were delayed in gametangiophore formation, and their antheridiophores had fan-shaped receptacles and short thick stalks (Figure 4H-I). As MpKS and MpKOL1 catalyze reactions prior to KA biosynthesis, we tested if KA could rescue the phenotypes of Mp*ks^ld^* and Mp*kol1^ld^.* Indeed, 2-μM KA application fully restored the thallus morphology and gametangiophore formation in these mutants, which further supported that KA biosynthesis has a pivotal role in the FR response of *M. polymorpha* (Figure 4F-I).

In addition to MpKS, the other TPS-e/f clade member, MpTPS1 (Mp6g05430), is known to produce *ent-*kaurene as a minor product (Kumar et al., 2016). However, Mp*tps1^ld^* mutants showed similar morphology to wild-type plants under cW or cW+cFR conditions, suggesting that Mp*TPS1* is not a major gene required for GA_Mp_ biosynthesis (Supplemental Figure 12A; Supplemental Figure 13). Mp*KOL2* and Mp*KOL3*, the homologs of Mp*KOL1*, are tandemly arranged in the genome, so we removed the whole genomic fragment containing both genes with CRISPR/Cas9^D10A^ nickase (Supplemental Figure 12B). Again, no phenotype related to FR response was observed, which is consistent with the lack of KA-producing activity for both enzymes (Supplemental Figure 13).

As MpKAOL1 could catalyze the KA to GA_12_ conversion, we hypothesized that this is a reaction leading to the biosynthesis of GA_Mp_ in *M. polymorpha*. As a fact, Mp*kaol1^ge^* mutants were defective in FR response, in a manner similar to the KA biosynthesis mutants. After 12 days of growth under cW+cFR, large wide and flat thallus was observed in Mp*kaol1^ge^*mutants (Figure 4F-G). Since MpKAOL1 is likely to act downstream of KA synthesis, these mutants were not supposed to be very sensitive to KA treatment. In the experiments, although 2 μM of KA slightly altered the thallus morphology under cW+cFR, while no significant changes could be observed at concentrations equal to or lower than 1 μM (Figure 4F-G; Supplemental Figure 7B-E). Mp*kaol1^ge^* mutants also showed a moderate delay in gametangiophore formation, which was insensitive to KA treatment. However, the antheridiophore morphology of Mp*kaol1^ge^*mutants was similar to that of wild-type plants, suggesting that the biosynthesis of GA_Mp_ might not be completely abolished by Mp*kaol1^ge^* (Figure 4H-I). Complete deletion of the other KAO homolog, Mp*kaol3^ld^*, failed to show any severe defects in FR response, yet double mutants would be needed for future research to carefully examine the redundancy with MpKAOL1 (Supplemental Figures 12C and 13).

### FR enrichment induced GA biosynthesis in a Mp*PIF*-dependent manner

Recent research reported that several GA biosynthesis genes were up-regulated under FR-enriched light conditions (Briginshaw et al., 2022). In our previous transcriptome data (Hernández-García et al., 2021), irradiation solely with FR light significantly increased the expression of Mp*KOL1* within 1 h, and the expression of Mp*CPS* and Mp*KAOL1* after 4 h of treatment. Such responses were only seen in the wild-type plants but not the Mp*pif^ko^* mutant, indicating that it is a process controlled by the Mpphy-MpPIF signaling module (Supplemental Figure 14A). Similarly, when we transferred 7-day-old plants grown under cW conditions to cW+cFR, Mp*PIF*-dependent up-regulation of Mp*CPS*, Mp*KS*, Mp*KOL1* and Mp*KAOL1* expression could be detected by quantitative polymerase chain reaction (qPCR) within 24 h (Figure 5A, Supplemental Figure 14B). Interestingly, Mp*KOL1* showed a slightly different expression pattern to the other three genes. MpKOL1 expression peaked at 8 h after induction but slightly decreased at the 24-h time point, while the expression of Mp*CPS*, Mp*KS* and Mp*KAOL1* gradually increased in the 24 h after induction, which might reflect different modes of transcriptional regulation. Consistent with the gene expression, we detected a higher level of endogenous GA_12_ in plants induced with cW+cFR than those kept under cW conditions, which suggested the accumulation of GA-related compounds under FR-enriched conditions (Figure 5B).

**Figure 5.**
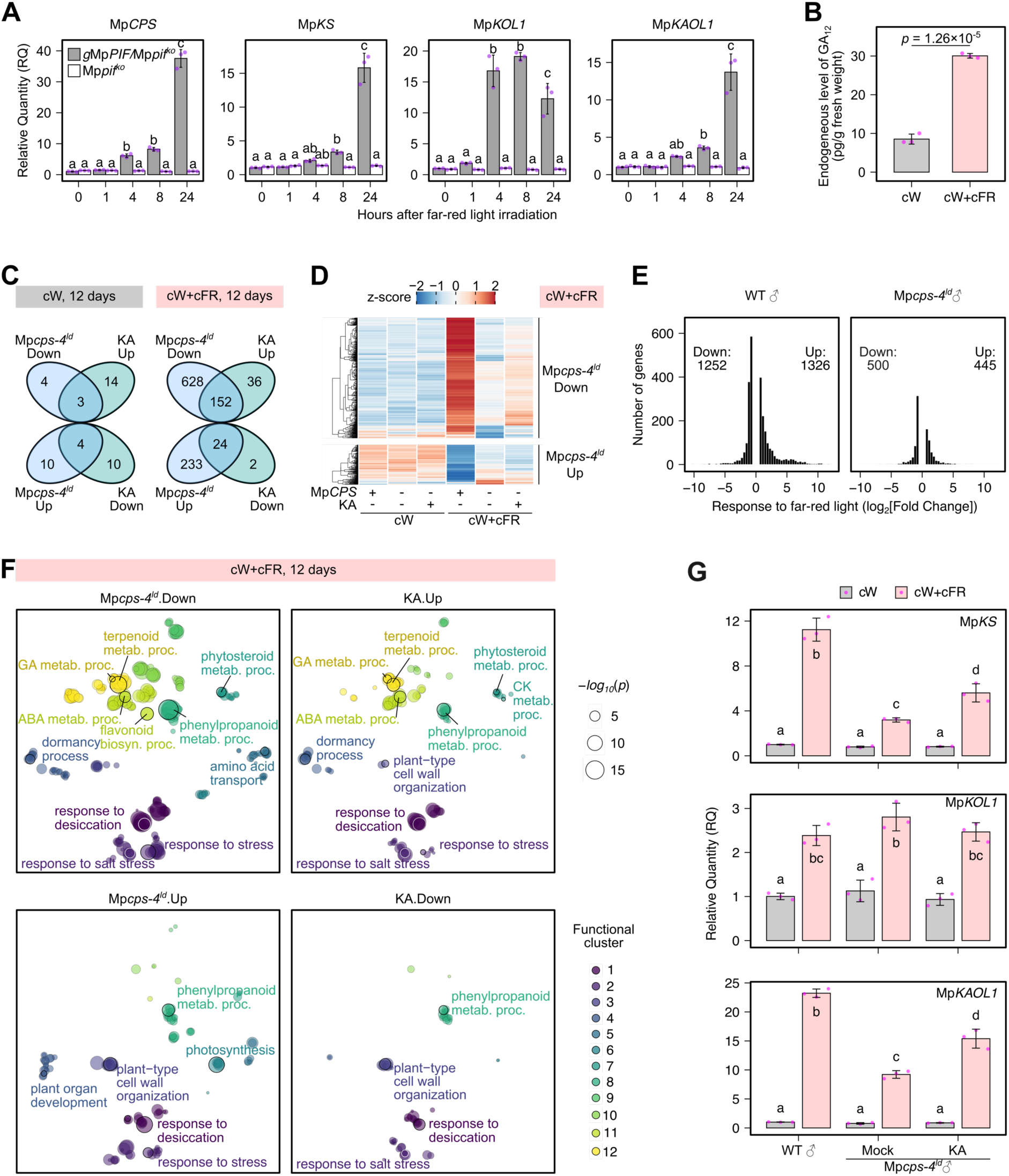
Transcriptional regulation related to GA biosynthesis in *M. polymorpha*. A, Relative expression level of GA biosynthesis genes by qPCR in Mp*pif^ko^* and its complementation line, cultured under cW for 7 days and then transferred to cW+cFR. B, endogenous level of GA_12_ in Tak-1 plants cultured under cW for 10 days, then under cW or cW+cFR for 4 days. The data for cW+cFR group is the same as in Figure 3A. The *p*-value was calculated by Student’s *t*-test. C, Number of differentially expressed genes (DEGs) between 12-day-old Mp*cps^ld^*and Tak-1 plants, or Mp*cps-4^ld^* plants grown with or without 2-μM KA under indicated light conditions (Wald test with B-H adjustment by DESeq2, DEG defined as adjusted *p*<0.01 and |log2(Fold Change)|>0.585). D, expression patterns of up- and down-regulated genes in Mp*cps-4^ld^*. E, Distribution of DEGs induced by FR enrichment in Tak-1 and Mp*cps-4^ld^* plants, comparing transcriptomes of cW+cFR to cW conditions. F, GO term enrichment analysis of DEG sets shown in (C). Each dot shows a significant enriched biological process term (*p*<0.01 by Fisher’s exact test), and semantically similar terms were plotted in color-coded clusters. Selected terms were highlighted with annotations. metab. proc., metabolic process. See Supplemental Data Set 2 for full lists. G, Relative expression level of GA biosynthesis genes by qPCR in plants cultured under cW for 11 days, then transferred to cW+cFR with or without 2-μM KA treatment. All bar plots with error bars (A-B, G) represent mean±SD from 3 biological replicates. Letters in A and G represent multiple comparisons with Tukey’s HSD test (adjusted *p*<0.05 for non-overlapping letters).

### GA-related hormone regulated transcriptional responses to FR enrichment

To explore gene expression changes related to GA biosynthesis in *M. polymorpha*, we analyzed transcriptomes from thalli grown under cW or cW+cFR for 12 days, either of wild-type plants, or of Mp*cps-4^ld^* mutants with or without 2-μM KA treatment. In agreement with the mild change in thallus morphology, only a few genes were differentially expressed between Mp*cps-4^ld^* and wild-type plants under cW conditions (Figure 5C; Supplemental Data Set 1). In particular, the expression of Mp*TPS6*, which encodes an enzyme producing *cis*-kolavenol (Jia et al., 2022), was significantly decreased in Mp*cps-4^ld^* and rescued by KA under cW. In contrast, 780 and 257 genes were down- and up-regulated, respectively, in Mp*cps-4^ld^* mutants under cW+cFR, suggesting a more active function of GA_Mp_ under FR-enriched conditions. A proportion of the differentially expressed genes could be rescued by KA application (Figure 5C; Supplemental Data Set 1). For most of the Mp*cps^ld^*-affected genes, the expression was strongly altered by FR enrichment in the wild-type plant, and the differential expression in Mp*cps-4^ld^* was essentially the reduction in FR-induced gene expression change (Figure 5D). If we compare the transcriptomes between cW+cFR and cW, the numbers of up- and down-regulated genes both declined by more than 60% in the Mp*cps-4^ld^* mutant (Figure 5E).

Using fussy gene ontology (GO) annotations generated with the Blast2GO algorithm (Conesa and Götz, 2008; Hernández-García et al., 2021), we performed GO enrichment analyses for biological processes on differentially expressed genes in Mp*cps-4^ld^*under cW+cFR (Figure 5F; Supplemental Data Set 2). Although KA application rescued only a limited proportion of gene expression changes in Mp*cps^ld^*, the patterns of enriched GO terms were quite similar in both comparisons. Mp*cps-4^ld^*down-regulated or KA-upregulated genes were enriched in GO terms related to stress response and secondary metabolism, among which top-ranked the phenylpropanoid metabolic process. When we carefully checked gene homologs, 23 genes putatively catalyzing biosynthesis of auronidins, bibenzyls or lignin monomers were up-regulated by FR enrichment in a Mp*CPS*-dependent manner (Supplemental Figure 15A). In the Mp*cps-4^ld^*down-regulated gene set, we also found 30 CYPs, 12 2-OGDs and 7 uridine 5’-diphospho-glucuronosyltransferases (UGTs), all of which were mostly homologs specific to *M. polymorpha* or liverworts, possibly associated with lineage-specific metabolites (Supplemental Figure 15B-D). It seemed that *M. polymorpha* was allocating more resources to stress and defense responses under FR-enriched conditions, yet less doing so when GA_Mp_ biosynthesis was defective. On the other side, GO enrichment captured photosynthetic genes in the Mp*cps-4^ld^* up-regulated gene set (Figure 5F), which suggested that this mutant might be more resistant to FR-induced chlorophyll reduction (Fredericq and de Greef, 1966). Cell-wall related enzymes and peroxidases were found in both down- and up-regulated gene sets, which was in agreement with the change in thallus growth allometry caused by FR enrichment or Mp*cps^ld^*, but might also be related to stress and defense responses (Supplemental Figure 15E-F).

Gene expressions of several phytohormone pathways were altered by Mp*cps^ld^* under cW+cFR. First of all, FR-induction of Mp*KS* and Mp*KAOL1* was reduced in Mp*cps-4^ld^*, which suggested that GA biosynthesis might be regulated by positive feedback in *M. polymorpha* (Supplemental Figure 15G). Such an effect was confirmed with qPCR, in which 3 days of KA treatment partially restored Mp*KS* and Mp*KAOL1* expression (Figure 5G). Mp*KOL1* expression was not affected by Mp*cps^ld^*or KA treatment, suggesting that it is not a target for such feedback regulation. For cytokinin, the biosynthesis enzymes Mp*IPT2* and Mp*LOG*, and the deactivation enzyme Mp*CKX2* were down-regulated in Mp*cps-4^ld^*(Supplemental Figure 15G). Previous research showed that Mp*CKX2* overexpression reduced the level of cytokinins, restricted thallus growth and increased thallus hyponasty under cW (Aki et al., 2019). It is possible that FR enrichment changed the thallus growth via coordinated deactivation of cytokinins and accumulation of GA_Mp_. Furthermore, expression of three abscisic acid (ABA) metabolic genes (MpABA4, MpNCED, MpCYP707A) and more than 40 ABA-responsive, LATE EMBRYOGENESIS ABUNDANT-like (LEA-like) proteins were induced by FR enrichment, similar to the response in *A. thaliana* (Michaud et al., 2023) (Supplemental Figure 15H). Such induction was reduced in Mp*cps-4^ld^* and restored by KA application, which showed similar tendency as the overall stress response.

## DISCUSSION

Light quality, i.e. the ratio of red and far-red lights, is an important environmental clue for nearly all land plants to optimize their growth and photosynthesis efficiency. Using a reverse-genetic approach, we showed that biosynthesis of gibberellin-related diterpenoids is required for many aspects of FR response in the liverwort *M. polymorpha*. Under FR-enriched conditions, wild-type plants often develop narrow and hyponastic thallus and begin to form gametangiophores, which is accompanied with increased expression of GA biosynthesis genes and accumulation of GA_12_ (Figures 1, 2, 5) (Fredericq and de Greef, 1966; Briginshaw et al., 2022). In contrast, Mp*cps^ld^* and other GA biosynthesis mutants developed wide and flat thallus, with a delay in the gametangiophore formation (Figures 1, 2, 4). Application of the biosynthesis intermediate KA rescued all mutants deficient of its biosynthesis (Figures 3, 4), indicating that these phenotypes were likely caused by the deficiency of some KA-derived diterpenoid compound(s), which we named GA_Mp_. In angiosperms, GA constantly modulates growth and development throughout the life cycle, mostly by promoting growth via cell elongation and/or cell division (Hedden, 2020). But in *M. polymorpha*, GA_Mp_ deficiency had little influence on the vegetative growth under white light conditions. The bioactivity of GA_Mp_ was observed only after its induced accumulation under FR-enriched conditions, suggesting that it served as a hormone coping with environmental changes, rather than constitutively regulating growth. Moreover, the thallus size was increased in GA_Mp_ biosynthesis mutants under cW+cFR (Figures 1, 4), implying that GA_Mp_ actively inhibits rather than promotes growth in the gametophyte of *M. polymorpha*.

FR enrichment induced hyponastic growth in the thallus of *M. polymorpha*, which is comparable to the increase of leaf hyponasty in shade-avoiding angiosperms. In *A. thaliana*, FR enrichment triggers the hyponastic growth by preferentially enhancing cell elongation in the abaxial side of the petiole (Küpers et al., 2023). While in *M. polymorpha*, such response relies on the regional growth driven by apical meristems. In previously reported end-of-day FR irradiation experiments, if the treatment was discontinued after several cycles, the newly-grown apical region resumed pleiotropic orientation, while the basal part remained hyponastic (Fredericq and Greef, 1968). With EdU analysis, we observed cell division patterns under FR-enriched conditions, which was most active near the apical meristem. The hyponastic growth in wild-type plants were marked with excessive cell division in the ventral side of the thallus, implying a role for differential cell proliferation in this process. The large and flat thallus of Mp*cps^ld^*mutants have more actively dividing cells in total, which showed no dorsiventrally biased distribution (Figure 1). Since Mp*cps^ld^* phenotypes could be complemented by expression of Mp*CPS* using the constitutive 35S promoter, which was equally active in dorsal and ventral tissues (Althoff et al., 2014), it is unlikely that FR-induced hyponasty is established directly by differential biosynthesis of GA_Mp_. This is similar to the situation in *A. thaliana* petioles, where GA served as a modulator for FR-induced hyponasty but showed no biased adaxial-abaxial activity (Küpers et al., 2023).

The erected, stalked gametangiophores of *M. polymorpha* could be considered as an extreme form of hyponastic growth, and dorsal tissues were evidently reduced in the cylindrical stalks of wild-type plants (Shimamura, 2016). The stalks of Mp*cps^ld^* gametangiophores were relatively short and thick, bearing dorsal air chambers similar to the vegetative thallus, which showed less biased dorsoventral growth (Figure 2; Supplemental Figure 5). This phenotype resembled that of Mp*kanadi* (Mp*kan*), which was depleted of the sole *M. polymorpha* homolog for KANADI, a family of transcription factors regulating tissue polarity in angiosperms (Briginshaw et al., 2022). Under cW+cFR, both Mp*kan* and Mp*cps^ld^* mutants were reduced in hyponasty, formed thallus-like gametangiophores with delay but generated fertile gametes, which could be explained by the dysfunction of Mp*kan* to up-regulate the expression of GA_Mp_ biosynthesis genes in response to FR irradiation (Briginshaw et al., 2022).

By tracing germ cell progenitors with MpBNB accumulation, we found that Mp*cps^ld^* mutants were delayed in germline differentiation (Figure 2). Many fern species use GA-related pheromones, i.e. antheridiogens, to control germline differentiation in the population. Particularly, antheridiogens promote the formation of male antheridia, and inhibits female archegonia in undifferentiated gametophytes (Näf et al., 1975; Tanaka et al., 2014; Hornych et al., 2021). In *M. polymorpha*, sexual differentiation is known to be determined by a sex chromosome-located gene (Iwasaki et al., 2021), and MpBNB accumulation was similarly delayed in both male and female Mp*cps^ld^* gametophytes, suggesting no sexual bias in the function of GA_Mp_.

We partially elucidated the biosynthetic pathway of GA_Mp_ with biochemical and genetic approaches. Previous bioinformatic research suggested that early steps of GA biosynthesis are conserved in land plants, and the *M. polymorpha* genome contains multiple homologs for KO and KAO (Bowman et al., 2017; Cannell et al., 2020; Yoshida et al., 2020). Using the yeast expression system, we detected the production of KA by MpKOL1 and GA_12_ by MpKAOL1, showing the two enzymes possessing enzymatic activities similar to their angiosperm homologs. Genetic analyses also supported that KA is synthesized via the sequential action of three single enzymes (Mp*CPS*, Mp*KS*, and Mp*KOL1*), as their mutants were consistently defective in FR light responses (Figure 4). Intriguingly, although GA_12_ production was almost completely blocked in Mp*kaol1^ge^* mutants, we only saw morphological phenotypes in the vegetative thallus, but not in the gametangiophores. As monooxygenases, it takes three sequential steps for CYP enzymes to catalyze the KA-to-GA_12_ conversion, first from KA to *ent*-7-hydroxy-kaurenoic acid (7OH-KA), then from 7OH-KA to GA_12_-aldehyde, and finally from GA_12_-aldehyde to GA_12_ (Helliwell et al., 2001a). In the GA-producing fungus *Fusarium fujikuroi*, even though GA_12_ production still occurs, it is largely bypassed through 3β-hydroxylation of GA_12_-aldehyde to form GA_14_-aldehyde, which is then converted into GA_14_ and other gibberellin compounds (Hedden et al., 1974; Urrutia et al., 2001; Rojas et al., 2001). Therefore, we could not conclude that Mp*kaol1^ge^*mutant phenotypes were caused by deficiency in GA_12_ production, and MpKAOL1 might work redundantly with other enzymes to produce other intermediates, synthesizing GA_Mp_ in a GA_12_-independent pathway.

Despite the conserved activities of MpKOL1 and MpKAOL1, we detected no KA synthesis activity for MpKOL2 or MpKOL3, and no GA_12_ production by MpKAOL3 (Figure 4). Also, no morphological abnormality related to far-red light response was observed in their loss-of-function mutants, suggesting that they are not major enzymes required for GA_Mp_ biosynthesis (Supplemental Figure 13). In the phylogenetic tree of KOs (CYP701s) (Supplemental Figure 8), all liverwort homologs formed a monophyletic clade with two major subclades, both containing sequences from Jungermanniopsida and Marchantiopsida species. MpKOL2 and MpKOL3 belong to a subclade different from MpKOL1, suggesting these KO homologs were diverged soon after the emergence of the most recent liverwort common ancestor. The phylogenetic relationship of KAOs (CYP88s) from liverworts and hornworts is less clear, but similarly, MpKAOL1 and MpKAOL3 belong to different clades deeply diverged in liverworts (Supplemental Figure 9). It remains to be explored if such divergence contributes to the diversification of diterpenoid metabolism in liverworts.

Comparing the biosynthesis of GA-related diterpenoids between *M. polymorpha* and *P. patens*, we see both conservation and divergence of this pathway in evolution. On the one hand, both bryophyte species share the biosynthesis route for KA with vascular plants. KO is present in almost all genomes and transcriptomes included in our phylogenetic analysis. As a recent study proposed that the ancestral *TPS* gene in land plants encoded a bifunctional CPS/KS enzyme (Jia et al., 2022), it is possible that production of KA is conserved in all major lineages of land plants. On the other hand, *M. polymorpha* and *P. patens* seem to take different routes to synthesize bioactive compounds from KA. Consistent with the lack of KAO (CYP88) homologs in all mosses, *P. patens* produced *ent*-3β-hydroxy-kaurenoic acid (3OH-KA) and *ent*-2α-hydroxy-kaurenoic acid (2OH-KA) from KA, with the former having physiological activities on protonemal cell differentiation (Miyazaki et al., 2018). Meanwhile, the production of GA_12_ in *M. polymorpha* opened the possibility of using C20-GAs in growth regulation, although no common GA from angiosperms was detected in either bryophyte species (Figure 3) (Hayashi et al., 2010).

Due to the lack of GIBBERELLIN-INSENSITIVE DWARF1 (GID1) receptors in bryophytes, it is unlikely that GA-related hormones are perceived by the canonical GID1-DELLA module in *M. polymorpha* or *P. patens*. Previously we reported that overexpression of Mp*DELLA* inhibited MpPIF-mediated FR responses, reducing gemma dormancy and delaying the formation of gametangiophores (Hernández-García et al., 2021). However, Mp*DELLA* overexpression strongly inhibited cell division and thallus growth, which was opposite to the effect of GA_Mp_ deficiency. It is possible that the functions of GA_Mp_ and MpDELLA remain uncoupled in *M. polymorpha*.

Gene expression analysis confirmed that Mp*CPS*, Mp*KS*, Mp*KOL1* and Mp*KAOL1* were induced by FR enrichment in an Mp*PIF*-dependent manner (Figure 5). The regulatory role of Mp*PIF* is further supported by phenotypic similarities between GA_Mp_ biosynthesis mutants and Mp*pif*. Under FR enriched conditions, Mp*pif* mutants developed wider, flatter and larger thallus, and failed to form any gametangiophores (Inoue et al., 2019; Streubel et al., 2023). In angiosperms, FR enrichment also activates the expression GA biosynthesis enzymes, yet more frequently regulating 2-OGD family genes such as GA20ox or GA3ox (Hisamatsu et al., 2005; Kohnen et al., 2016; Küpers et al., 2023), which are absent from bryophytes. Considering the putatively different mechanism for GA perception, GA biosynthesis might be independently incorporated into the gene regulatory network by liverworts and angiosperms in evolution, in response to the common threat of a FR-enriched environment.

## METHODS

### Plant materials and maintenance

The *M. polymorpha* subsp. *ruderalis* accessions Takaragaike-1 (Tak-1) and Takaragaike-2 (Tak-2) were used as male and female wild-type materials, respectively (Ishizaki et al., 2008). For maintenance, *M. polymorpha* plants were cultured aseptically on half-strength Gamborg’s B5 medium (Gamborg et al., 1968) with 1% agar at 22°C under continuous white light (40-50 μmol photons m^-2^ s^-1^), which was supplemented by cold cathode fluorescent lamps (CCFLs, OPT-40C-N-L from Optrom, Japan). If not specified, this medium and temperature was used for all aseptic cultures for *M. polymorpha*.

### Light sources

In all the experiments, plants were cultured under continuous white light conditions (cW, approximately 40 μmol photons m^-2^ s^-1^) with or without continuous far-red light (cFR, approximately 25 μmol photons m^-2^ s^-1^). For thallus morphology observation, cW and cFR were supplemented by CCFLs and light-emitting diodes (LEDs), respectively, in the multi-chambered incubator (LH-80CCFL-6CT from NK systems, Japan). For gametangiophore induction of aseptic cultures, cW was supplemented by CCFLs (ST-40C-BN, Shinshu Trading, Japan) and cFR was supplemented by LEDs (IR LED STICK, NAMOTO, Japan). For plants grown on the vermiculite, cW was supplemented by fluorescent tubes (FLR40SN/M/36, Toshiba, Japan) and cFR was supplemented by LEDs (VBL-TFL600-IR730, Valore, Japan).

### Construction of mutants and transgenic plants

To create large-deletion mutants of a target gene, four guide RNAs (gRNAs) with target sequences flanking the CDS were first cloned separately into pMpGE_En04, pBCGE12, pBCGE23 and pBCGE34 (Hisanaga et al., 2019; Koide et al., 2020) by ligation of BsaI-digested vectors with annealed complementary DNA oligos. Then the fragments containing the MpU6-1 promoter and gRNAs were digested from the pBCGE12, pBCGE23 and pBCGE34 constructs and inserted between the BglI sites of the pMpGE_En04 construct to create a multiplex entry vector. Finally, this entry vector was recombined with pMpGE017 or pMpGE018 (Hisanaga et al., 2019) using Gateway LR Clonase II (Thermo Fisher Scientific) to generate the binary vector for plant transformation. To generate the Mp*kaol1^ge^* mutants, the gRNA was designed to target sequences near the start codon of Mp*KAOL1*. The corresponding DNA oligos were annealed and ligated into the BsaI sites of pMpGE_En03, then subcloned to the binary vector pMpGE011 (Sugano et al., 2018).

For the complementation of mutants and/or subcellular localization observations, CDSs of Mp*CPS*, Mp*KS*, Mp*KOL1* and Mp*KAOL1* were amplified from the complementary DNA (cDNA) of Tak-1 plants and directionally cloned into pENTR/D-TOPO (Thermo Fisher Scientific) to create entry vectors without stop codon. To mutate the gRNA target sequence in Mp*KAOL1*, the whole plasmid of pENTR-MpKAOL1-CDS was amplified with the site-directed mutagenesis primers MpKAOL1-mut-F and MpKAOL1-mut-R, then re-circularized to generate pENTR-MpKAOL1-CDSmut. All the CDSs were then subcloned from entry vectors to pMpGWB106 or pMpGWB306 using Gateway LR reactions, which were used for expressing the target proteins under the 35S promoter with an in-frame Citrine fusion at the C terminus (Ishizaki et al., 2015).

All the binary vectors were introduced into *M. polymorpha* by transformation of regenerating thalli using the *Agrobacterium tumefaciens* strain GV2260 as previously described (Kubota et al., 2013). Successful mutations were identified by genotyping PCR and Sanger-sequenced to confirm non-identical alleles. See Supplemental Tables 1-3 for full lists of plant materials generated by this research, and the plasmids and DNA oligos used for construction.

### Measurement of thallus morphology

For thallus morphological observation, plants were grown from gemmae on the medium containing 1% sucrose under cW or cW+cFR in a multi-chambered incubator (LH-80CCFL-6CT from NK systems, Japan). After 12 days of growth, camera photos of the plants were taken vertically from the top. Then medium blocks with individual plants were carefully cut out, aligned at a certain distance to a fixed camera for taking side-view photos. Typically, the two apical meristems from one gemma both develop into mature thallus branches, and we define the part developed from a single gemma meristem as a “half thallus”. These half thalli were cut apart, flattened on a filter paper and photographed for the measurements of length, width and area. All the measurements were done in Fiji (Schindelin et al., 2012) as describe in Supplemental Figure 16 with in-house macro scripts, which are available at https://github.com/dorrenasun/Mp_GA_biosynthesis. Data was excluded for a whole plant if more than two apical meristems were active in the gemma, or for a half thallus if it showed severe defects in dorsoventral differentiation.

### EdU assay

To analyze the cell division activity, plants were grown on the medium containing 1% sucrose under cW+cFR for 7 days, then soaked for 2 h in liquid half-strength Gamborg’s B5 medium containing 20 mM 5-ethynyl-2’-deoxyuridine (EdU). Then the samples were fixed in 3.7% formaldehyde for 1 h, washed twice with phosphate buffer saline (PBS, 5 min each), and treated with 0.5% Triton X-100 in PBS for 20 min. After two washes with 3% bovine serum albumin (BSA) in PBS, the samples were kept in the reaction mixture from the Click-iT EdU Imaging Kit with Alexa Fluor 555 (Thermo Fisher Scientific, #C10338) for 1 h in darkness, then washed twice with PBS before soaked overnight in 20% caprylyl sulfobetaine (#D4246, TCI, Japan) to remove the chlorophyll. After another two washes with PBS, the thalli were treated with 75.5% (w/v) iohexol (GE Healthcare Pharma) for 1 h and mounted on glass slides following the iTOMEI protocol (Sakamoto et al., 2022). Image stacks of the apical regions were obtained with a confocal laser scanning microscope (Olympus FLUOVIEW FV1000) at 5-μm steps in the z direction. Excitation and emission wavelengths were 543 nm and 505-605 nm, respectively. Numbers of EdU-positive nuclei were quantified in the Z-projections of the image stacks using the StarDist plugin of Fiji and in-house scripts (Schmidt et al., 2018; Schindelin et al., 2012).

### Observation of gametangiophores

To observe the progress of gametangiophore formation in aseptic culture, plants were first grown from gemmae on the medium containing 1% sucrose under cW for 7 days, then half thalli developed from single gemma meristems were cut apart and cultured on fresh medium plates under cW+cFR. Chemical treatments were included in the medium of second-stage culturing, starting together with the FR irradiation. Apical regions of each plant were observed under a stereoscope (Olympus SZX16) every day, and the emergence of gametangiophores was recorded if a stalk or dark-green primordia was formed. Images of gametangiophores were taken with stereoscopes (Olympus SZX16 or Leica M205C). For morphological observation of gametangiophores, the plants were also grown on the vermiculite in the open air. After an initial culturing of 7-14 days from gemmae under aseptic maintenance conditions, thallus fragments were planted in pots containing vermiculite and grown under cW+cFR conditions with regular watering. For sectioning of gametangiophores, fresh stalks or receptacles were embedded in 5-6% agar and sectioned with a vibratome (DOSAKA LinearSlicer Pro7) at the desired thickness. Images of the sections were obtained with the microscope under bright field (Keyence BZ-X710).

### Microscopy of MpBNB-Cit

To capture the early stage of gametangium differentiation, male and female plants carrying the Mp*BNB-Cit* knock-in locus were cultured from gemmae on the medium containing 1% sucrose under cW+cFR for 11 and 14 days, respectively. For male plants, half thalli developed from single gemma meristems were collected for observation. For female plants, the second bifurcation has already occurred, so one of the four apical regions was selected randomly on each plant for observation. Thallus fragments were fixed with 4% paraformaldehyde (PFA) for 1 h at room temperature, washed twice with PBS and soaked in the ClearSeeAlpha solution (Kurihara et al., 2021) for 2 days to remove chlorophyll. After that, the samples were washed twice with PBS, stained with 1 mg/mL calcofluor white for 10 min and washed again twice with PBS. Finally, the samples were infiltrated with 75.5% (w/v) iohexol (GE Healthcare Pharma) for 1 h and mounted onto slides. Image stacks were obtained with the fluorescent microscope (Keyence BZ-X710) at 5-μm steps and processed into full-focus projections with BZ-X Analyzer, and the number of Citrine-positive nuclei was counted in Fiji (Schindelin et al., 2012). The BZ-X DAPI (Excitation: 360±20 nm; Emission: 460±25 nm; Dichroic mirror: 400 nm) and customized (Excitation: 500±10 nm; Emission: 535±15 nm; Dichroic mirror: 515 nm) filters were used for acquiring signals from calcofluor white and Citrine, respectively.

### LC-MS/MS analysis of endogenous GAs

To analyze the endogenous GAs, most plants were cultured from gemmae on the medium containing 1% sucrose under cW for 10 days, then under cW+cFR for another 4 days. Tak-1 wild-type plants were also cultured under cW for 14 days to examine the effect of FR enrichment. For each sample, 3 g of thallus tissue was harvested, frozen with liquid nitrogen, and stored at - 80 °C before extraction. To prepare samples for LC-MS/MS analysis, each frozen sample was homogenized in 15 mL acetone with ^2^H_2_-labelled authentic compounds, then left at 4 °C for a 2-h extraction. After filtering with defatted cotton, the acetone extract was concentrated to approximately 1 mL with nitrogen blow and mixed with 1 mL acetonitrile. The mixture was extracted for 3 times with 2 mL hexane, and the remaining aqueous phase was alkalified with 1 mL saturated solution of NaHCO_3_. After two more extractions with 2 mL chloroform, the aqueous phase was passed through a cartridge filled with polyvinylpyrrolidone (PVP), acidified to pH2-3 with HCl (6 M), and sequentially purified with a reverse-phase cartridge (Oasis HLB 3 cc/60 mg, Waters), an anion-exchange cartridge (Bond Elut DEA 100 mg/1 mL, Agilent), and a silica cartridge (Sep-Pak Silica 1 cc Vac Cartridge, Waters). The final elute was dried up with a vacuum centrifuge concentrator and dissolved in 1% acetic acid before loading to the LC-MS/MS.

The LC-MS/MS system consisted of an ultra-performance LC (ExionLC, Sciex) equipped with a reverse-phase column (CORTECS UPLC C18+, φ1.6 μm, 2.1×100 mm, Waters) and a quadrupole time-of-flight mass spectrometer (X500R QTOF, Sciex). LC separations were performed at 40 °C with a flow rate of 0.3 mL/min using solvent A (0.05% acetic acid in water) and solvent B (0.05% acetic acid in acetonitrile) and the following program: a linear gradient of B from 3% to 65% over 17 min, followed by an isocratic elution with 98% of B for 2 min. Quantification of GAs was performed in the multiple reaction monitoring (MRM) mode, and the mass spectrum of GA_12_ was confirmed in the TOF-MS/MS mode.

### Phylogenetic analysis

For the phylogenetic analysis of KO and KAO homologs, the known protein sequences from *A. thaliana* and *Oryza sativa* were used as the input for BLAST (Altschul et al., 1990) search in the annotated proteins from published genomes and transcriptomes (See Supplemental Data Set 3 for a full list of species and references). Considering the relative low redundancy of these enzymes, the top 20 BLAST hits from each species were retrieved and aligned with MAFFT using the progressive FFT-NS-2 algorithm (Katoh and Standley, 2013). After removing the positions with >80% gaps, an initial maximum likelihood phylogenetic tree was built with all candidate sequences using IQ-TREE 2 with automatic substitution model selection (ModelFinder) and 1000 bootstraps from the ultrafast bootstrap approximation (UFBoot) (Minh et al., 2020; Kalyaanamoorthy et al., 2017; Hoang et al., 2018). Highly-relevant candidate sequences and outgroup sequences were selected from the initial tree and re-aligned with MAFFT using the L-INS-I algorithm (Katoh and Standley, 2013) and trimmed off positions with >98% gaps. The final maximum likelihood tree was inferred using IQ-TREE 2 with automatic substitution model selection (LG+I+G4 for both KOs and KAOs) and 1000 standard non-parametric bootstraps (Minh et al., 2020; Kalyaanamoorthy et al., 2017; Hoang et al., 2018; Guindon et al., 2010). The complete pipeline including scripts, sequence alignments, and the data file for the phylogenetic trees are available at https://github.com/dorrenasun/Mp_GA_biosynthesis. The ggtree R package suite was used for visualization (Yu, 2022; Xu et al., 2022; Yu, 2020b; Yu et al., 2018, 2017).

### Enzymatic assay in *P. pastoris* and GC-MS analysis

To express MpKOLs and MpKAOLs in the yeast *P. pastoris*, the CDSs were amplified from pENTR-CDS entry vectors and inserted into the pPICZA vector (Thermo Fisher Scientific) with homologous recombination using the seamless ligation cloning extract (SLiCE) from *Escherichia coli* (Zhang et al., 2012; Motohashi, 2015). The CDSs of At*KO* (AT5G25900) and At*KAO1* (AT1G05160) were amplified from the cDNA of 14-day-old Col-0 seedlings and cloned into pPICZA with the same method. All these plasmids were transformed into a previously described *P. pastoris* X-33 strain carrying the *A. thaliana* cytochrome reductase gene, At*ATR1* (AT4G24520) (Katsumata et al., 2008), following the manufacturer’s protocol. The selected transformants were incubated in 2 mL of BMG medium (100 mM potassium phosphate (pH 6.0), 1.34% yeast nitrogen base (YNB), 4×10^-5^% biotin, 1% glycerol) with shaking at 30 °C until they reached an OD_600_ value of 2. Cells were collected by centrifugation and resuspended in 50 mL of MM medium (1.34% YNB, 4×10^-5^% biotin, 0.5% methanol), and cultured at 30 °C with shaking and addition of methanol every 24 h (final concentration: 0.5% v/v) to maintain protein induction. After 24 h of initial culturing, substrates (7 µg of *ent*-kaurene, or 15 µg of KA) were added into the medium. After another 48 h of incubation, the supernatants were isolated with centrifugation and extracted two times with equal volume of ethyl acetate. The organic phase from the two extractions were combined together, concentrated *in vacuo*, and derivatized with diazomethane in ether solution. After that, the samples were concentrated to approximately 100 μL with nitrogen blow and subjected to GC-MS analysis with a GC (Agilent 6890) equipped with a DB-1 capillary column (15 m/0.25 mm/0.25 µm, Agilent J&W) and a mass selective detector (Agilent 5975C MSD, ionization energy at 70 eV). The oven temperature was programmed as described previously (Hayashi et al., 2006).

### Subcellular localization of proteins

To observe the subcellular localizations of MpCPS, MpKOL1 and MpKAOL1, plants were grown from gemmae on the medium containing 1% sucrose under cW+cFR conditions for 7 days. The thallus tissues were fixed with 4% PFA at room temperature for 20 min, washed twice with PBS, and stained with 1 mg/mL calcofluor white for 10 min. After another two washes with PBS, the samples were soaked in 75.5% (w/v) iohexol (GE Healthcare Pharma) for 1 h and mounted onto slides. For MpKS, fresh gemmae from gemma cups were mounted in water. For all experiments, complementation lines which used the 35S promoter to express target proteins fused with Citrine in the C-teminus were used for observation, and the parent lines of genetic mutants were used as the negative control.

Fluorescent images were acquired using a confocal microscope (Leica TCS SP8X Falcon) equipped with a hybrid detector (HyD). For the observation of MpCPS-Cit and MpKS-Cit, calcofluor white signals were excited with the 405 nm UV laser, and obtained in the xyz mode within the 425-435 nm wavelength region; the Citrine signals were excited with the the pulsed white light laser (WLL) at 488 nm, and obtained at 500-541 nm with time gating (1.8-12 ns for MpCPS-Cit; 1.2-12 ns for MpKS-Cit); the autofluorescence from chlorophyll was excited with the WLL at 592 nm, and obtained at 680-700 nm. For the observation of MpKOL1-Cit and MpKAOL1-Cit, calcofluor white signals were excited with the 405 nm UV laser, and obtained in at 425-475 nm; the Citrine signals were excited with the WLL at 495 nm, and obtained at 500-530 nm with time gating (1.2-6.0 ns); the autofluorescence from chlorophyll was excited with the WLL at 649 nm, and obtained at 655-755 nm.

### RNA extraction and qPCR

For qPCR experiments, 50-100 mg of whole thallus was collected for each sample, which is frozen with liquid nitrogen and crashed into fine powders with metal beads in a shaking device (Shake Master Auto, BMS-A20TP, Bio Medical Science, Japan). The total RNA was extracted with the TRIzol reagent (Thermo Fisher Scientific) following the manufacturer’s protocol. After treatment with RQ1 RNase-Free DNase (Promega), the total RNA was reverse-transcribed using ReverTra Ace (Toyobo Life Science) and the (dT)_20_ oligo. Quantitative real-time PCR reactions with Taq polymerase prepared following (Pluthero, 1993) and SYBR Green I Nucleic Acid Gel Stain (Lonza) was performed in triplicates using the CFX96 real-time PCR detection system (Bio-Rad) with the following program: an initial denaturation for 30 s at 95 °C, then 40 cycles of 5 s at 95 °C followed by 30 s at 60 °C. A standard melting curve analysis was performed at the end of the program to validate amplified products. Mp*EF1* (Mp3g23400) was used as the reference gene for quantification, and the relative gene expression levels were calculated following the method in (Hellemans et al., 2008). All primers for target and reference genes were listed in Supplemental Table 3, and the amplification efficiency for each primer pair was measured with serial dilution of mixed cDNA samples.

### Transcriptome analysis

For transcriptome analysis, plants were cultured from gemmae on the medium with 1% sucrose, either containing 2 μM KA or the solvent control. After 12 days of growth under cW or cW+cFR, ∼50 mg whole thallus was collected for each sample, frozen with liquid nitrogen and homogenized with metal beads in a shaking device (Shake Master Auto, BMS-A20TP, Bio Medical Science, Japan). The total RNA was isolated with the RNeasy Plant Mini Kit (QIAGEN), and the RNA quality was confirmed using the Bioanalyzer RNA 6000 pico assay (Agilent). The mRNA was enriched with the NEBNext Poly(A) mRNA Magnetic Isolation Module (New England Biolabs, #E7490). The library was prepared with the NEBNext Ultra II Directional RNA Library Prep Kit for Illumina (New England Biolabs, #E7760) and amplified using NEBNext Multiplex Oligos for Illumina (96 Unique Dual Index Primer Pairs Set 2, New England Biolabs, #E6442). After quality check with the Bioanalyzer High Sensitivity DNA assay (Agilent), the samples were sequenced with the NextSeq 500 system using the NextSeq 500/550 High Output Kit v2.5 for 75 cycles (Illumina). Approximately 7.8 million of single-end reads were obtained for each sample, and automatically de-multiplexed by the BaseSpace Sequence Hub (Illumina).

The sequence reads were quasi-mapped to the *M. polymorpha* MpTak_v6.1 genome (Iwasaki et al., 2021) and quantified using Salmon (v1.9.0) (Patro et al., 2017), with U-chromosome transcripts excluded from the index. After that, differential gene expression analysis was performed in R (v4.2.2) with the package DESeq2 (v1.38.3) in default settings (Love et al., 2014). Gene ontology enrichment analyses was performed with the R package topGO (v2.50.0) (Alexa and Rahnenfuhrer, 2022) using annotations previously described (Hernández-García et al., 2021) and the classic Fisher’s exact test, and *p*-values were left unadjusted following the package’s instructions. To visualize the enriched terms, semantic similarities were calculated using the R package GOSemSim (v2.24.0) (Yu et al., 2010; Yu, 2020a), and used for building a two-dimensional map with the package Rtsne (Krijthe, 2015). The transcriptome data is deposited to the Sequence Read Archive at the DNA Data Bank of Japan (DDBJ) under the Bioproject PRJDB15786. The scripts for data analysis are available at https://github.com/dorrenasun/Mp_GA_biosynthesis.

## Supplemental Data

**Supplemental Figure 1.** Gibberellin (GA) biosynthesis pathway in vascular plants, showing compounds analyzed or used for treatment in this research.

**Supplemental Figure 2.** Genotype information for Mp*cps^ld^*, Mp*ks^ld^*, Mp*kol1^ld^*, and Mp*kaol1^ge^* mutants.

**Supplemental Figure 3.** Thallus morphology of Mp*cps^ld^* mutants in Tak-2 background under cW+cFR.

**Supplemental Figure 4.** Morphology of Mp*cps^ld^* plants during gametangiophore formation in the aseptic culture.

**Supplemental Figure 5.** Sections of gametangiophores in wild-type and Mp*cps^ld^* plants, as well as the fertility test.

**Supplemental Figure 6.** LC-MS/MS and GC-MS profiles for KA and GA_12_ detection.

**Supplemental Figure 7**. Response of Mp*cps-4^ld^* and Mp*kaol1-5^ge^* to different concentrations of KA.

**Supplemental Figure 8.** Phylogenetic tree of KO homologs in land plants.

**Supplemental Figure 9**. Phylogenetic tree of KAO and closely-related CYP enzymes in land plants.

**Supplemental Figure 10.** Heatmap of percentage identity for KAO and closely-related CYP enzymes in land plants.

**Supplemental Figure 11.** Subcellular localizations of GA biosynthesis enzymes in *M. polymorpha*.

**Supplemental Figure 12.** Genotype information for Mp*tps1^ld^*, [Mp*kol2* Mp*kol3*]*^ld^* and Mp*kaol3^ld^* mutants.

**Supplemental Figure 13.** Phenotypes of Mp*tps1^ld^*, [Mp*kol2* Mp*kol3*]*^ld^* and Mp*kaol3^ld^*mutants.

**Supplemental Figure 14.** Up-regulation of GA biosynthesis genes by FR irradiation.

**Supplemental Figure 15**. Heatmaps of genes from selected pathways or gene families, showing differential expression in Mp*cps-4^ld^*under cW+cFR.

**Supplemental Figure 16.** Quantification method for thallus morphology.

**Supplemental Table 1**. List of plant materials

**Supplemental Table 2**. List of plasmids

**Supplemental Table 3**. List of DNA oligos

**Supplemental Data Set 1**. Differentially expressed genes in the transcriptome analysis

**Supplemental Data Set 2**. Enriched gene ontology (GO) terms in the transcriptome analysis

**Supplemental Data Set 3**. List of genome and transcriptome sources for phylogenetic analysis

**Supplemental References**.

## AKNOWLEDGEMENTS

The authors thank Ryunosuke Kusunoki for plasmid construction and collecting initial experimental data; Wakako Fukuda for data collection during GC-MS analysis; Takefumi Kondo and Yukari Sando for the support on RNA sequencing; Megumi Iwano for the help on confocal microscopy and Chikako Inoue for plant maintenance. This work has been supported by MEXT KAKENHI Grant Number JP19H05675 and JSPS KAKENHI Grant Numbers JP17H07424, JP22H00417 to T.K.; JSPS KAKENHI Grant Numbers JP 24380060, JP15H04492 to M.N. and H.K.; JSPS KAKENHI Grant Number JP16K18693 to S.M.; as well as the International Collaborative Research Program of Institute for Chemical Research, Kyoto University (Grant Numbers 2020-53, 2021-60, 2022-63, 2023-74) to T.K. and S.Yamaguchi.

## AUTHOR CONTRIBUTIONS

R.S. and M.O. collected and analyzed most of the final data with the help and supervision from K.M., Y.Y., S.Yamaoka., R.N. and T.K.. M.O., S.M., T.I., K.M., M.N. and S.Yamaguchi analyzed endogenous GAs. S.M. and H.K. assayed enzymatic activity. R.K. performed initial experiments that conceptualized the research. H.K., M.N., S.Yamaguchi and T.K. organized the project. R.S., S.Yamaoka., R.N and T.K. wrote the paper with the review and editing from all authors.

## REFERENCES

Achard, P., Gusti, A., Cheminant, S., Alioua, M., Dhondt, S., Coppens, F., Beemster, G.T.S., and Genschik, P. (2009). Gibberellin Signaling Controls Cell Proliferation Rate in Arabidopsis. Current Biology 19: 1188–1193.

Aki, S.S., Mikami, T., Naramoto, S., Nishihama, R., Ishizaki, K., Kojima, M., Takebayashi, Y., Sakakibara, H., Kyozuka, J., Kohchi, T., and Umeda, M. (2019). Cytokinin Signaling Is Essential for Organ Formation in Marchantia polymorpha. Plant and Cell Physiology 60: 1842–1854.

Alexa, A. and Rahnenfuhrer, J. (2022). topGO: Enrichment Analysis for Gene Ontology. R package version 2.50.0.

Althoff, F., Kopischke, S., Zobell, O., Ide, K., Ishizaki, K., Kohchi, T., and Zachgo, S. (2014). Comparison of the MpEF1α and CaMV35 promoters for application in Marchantia polymorpha overexpression studies. Transgenic Res 23: 235–244.

Altschul, S.F., Gish, W., Miller, W., Myers, E.W., and Lipman, D.J. (1990). Basic local alignment search tool. Journal of Molecular Biology 215: 403–410.

Beall, F.D., Yeung, E.C., and Pharis, R.P. (1996). Far-red light stimulates internode elongation, cell division, cell elongation, and gibberellin levels in bean. Can. J. Bot. 74: 743–752.

Bou-Torrent, J., Galstyan, A., Gallemí, M., Cifuentes-Esquivel, N., Molina-Contreras, M.J., Salla-Martret, M., Jikumaru, Y., Yamaguchi, S., Kamiya, Y., and Martínez-García, J.F. (2014). Plant proximity perception dynamically modulates hormone levels and sensitivity in Arabidopsis. Journal of Experimental Botany 65: 2937–2947.

Bowman, J.L. et al. (2017). Insights into land plant evolution garnered from the *Marchantia polymorpha* genome. Cell 171: 287–304.e15.

Briginshaw, L.N., Flores-Sandoval, E., Dierschke, T., Alvarez, J.P., and Bowman, J.L. (2022). KANADI promotes thallus differentiation and FR-induced gametangiophore formation in the liverwort Marchantia. New Phytologist 234: 1377–1393.

Cannell, N., Emms, D.M., Hetherington, A.J., MacKay, J., Kelly, S., Dolan, L., and Sweetlove, L.J. (2020). Multiple Metabolic Innovations and Losses Are Associated with Major Transitions in Land Plant Evolution. Current Biology 30: 1783–1800.e11.

Cao, J.-G., Wang, Q.-X., Zou, H.-M., Dai, X.-L., and Cao, T. (2013). New Observations on the Morphology and Structure of *Marchantia polymorpha* Gametophores in Sexual Reproduction Adaption. Plant Science Journal 31: 555.

Chiang, H.-H., Hwang, I., and Goodman, H.M. (1995). Isolation of the Arabidopsis *GA4* locus. the Plant Cell 7: 195–201.

Chiyoda, S., Ishizaki, K., Kataoka, H., Yamato, K.T., and Kohchi, T. (2008). Direct transformation of the liverwort Marchantia polymorpha L. by particle bombardment using immature thalli developing from spores. Plant Cell Reports 27: 1467–1473.

Conesa, A. and Götz, S. (2008). Blast2GO: A comprehensive suite for functional analysis in plant genomics. International Journal of Plant Genomics 2008.

Djakovic-Petrovic, T., Wit, M.D., Voesenek, L.A.C.J., and Pierik, R. (2007). DELLA protein function in growth responses to canopy signals. Plant Journal 51: 117–126.

Downs, R.J., Hendricks, S.B., and Borthwick, H.A. (1957). Photoreversible Control of Elongation of Pinto Beans and Other Plants under Normal Conditions of Growth. Botanical Gazette 118: 199–208.

Dubois, P.G., Olsefski, G.T., Flint-Garcia, S., Setter, T.L., Hoekenga, O.A., and Brutnell, T.P. (2010). Physiological and Genetic Characterization of End-of-Day Far-Red Light Response in Maize Seedlings. Plant Physiology 154: 173–186.

Fredericq, H. (1964). Influence formatrice de la lumière rouge-foncé sur le développement des thalles de Marchantia polymorpha L. Bull. Soc. Roy. Bot. Belgique 98: 67–76.

Fredericq, H. and de Greef, J. (1966). Red (R), Far-red (FR) photoreversible control of growth and chlorophyll content in light-grown thalli of Marchantia polymorpha L. Die Naturwissenschaften 53: 337.

Fredericq, H. and Greef, J.D. (1968). Photomorphogenic and Chlorophyll Studies in the Bryophyte Marchantia polymorpha. I. Effect of Red, Far-red Irradiations in Short and Long-term Experiments. Physiol Plant 21: 346–359.

Gabriele, S., Rizza, A., Martone, J., Circelli, P., Costantino, P., and Vittorioso, P. (2009). The Dof protein DAG1 mediates PIL5 activity on seed germination by negatively regulating GA biosynthetic gene AtGA3ox1: DAG1 represses seed germination via PIL5 signalling. The Plant Journal 61: 312–323.

Gamborg, O.L., Miller, R.A., and Ojima, K. (1968). Nutrient requirements of suspension cultures of soybean root cells. Experimental Cell Research 50: 151–158.

García-Martínez, J.L., Keith, B., Bonner, B.A., Stafford, A.E., and Rappaport, L. (1987). Phytochrome Regulation of the Response to Exogenous Gibberellins by Epicotyls of *Vigna sinensis*. Plant Physiol. 85: 212–216.

Guindon, S., Dufayard, J.-F., Lefort, V., Anisimova, M., Hordijk, W., and Gascuel, O. (2010). New algorithms and methods to estimate maximum-likelihood phylogenies: Assessing the performance of PhyML 3.0. Systematic Biology 59: 307–321.

Hayashi, K., Horie, K., Hiwatashi, Y., Kawaide, H., Yamaguchi, S., Hanada, A., Nakashima, T., Nakajima, M., Mander, L.N., Yamane, H., Hasebe, M., and Nozaki, H. (2010). Endogenous diterpenes derived from *ent-*kaurene, a common gibberellin precursor, regulate protonema differentiation of the moss *Physcomitrella patens*. Plant Physiology 153: 1085–1097.

Hayashi, K. ichiro, Kawaide, H., Notomi, M., Sakigi, Y., Matsuo, A., and Nozaki, H. (2006). Identification and functional analysis of bifunctional *ent-*kaurene synthase from the moss *Physcomitrella patens*. FEBS Letters 580: 6175–6181.

Hedden, P. (2020). The Current Status of Research on Gibberellin Biosynthesis. Plant and Cell Physiology 61: 1832–1849.

Hedden, P., MacMillan, J., and Phinney, B.O. (1974). Fungal products. Part XII. Gibberellin A14-aldehyde, an intermediate in gibberellin biosynthesis in Gibberella fujikuroi. Journal of the Chemical Society, Perkin Transactions 1: 587–592.

Hellemans, J., Mortier, G., De Paepe, A., Speleman, F., and Vandesompele, J. (2008). qBase relative quantification framework and software for management and automated analysis of real-time quantitative PCR data. Genome Biology 8.

Helliwell, C.A., Chandler, P.M., Poole, A., Dennis, E.S., and Peacock, W.J. (2001a). The CYP88A cytochrome P450, *ent*-kaurenoic acid oxidase, catalyzes three steps of the gibberellin biosynthesis pathway. Proceedings of the National Academy of Sciences of the United States of America 98: 2065–2070.

Helliwell, C.A., Poole, A., James Peacock, W., and Dennis, E.S. (1999). Arabidopsis *ent* - kaurene oxidase catalyzes three steps of gibberellin biosynthesis. Plant Physiology 119: 507–510.

Helliwell, C.A., Sheldon, C.C., Olive, M.R., R.Walker, A., Zeevaart, J.A.D., Peacock, W.J., and Dennis, E.S. (1998). Cloning of the *Arabidopsis ent-*kaurene oxidase gene *GA3*. Proceedings of the National Academy of Sciences of the United States of America 95: 9019–9024.

Helliwell, C.A., Sullivan, J.A., Mould, R.M., Gray, J.C., James Peacock, W., and Dennis, E.S. (2001b). A plastid envelope location of *Arabidopsis ent*-kaurene oxidase links the plastid and endoplasmic reticulum steps of the gibberellin biosynthesis pathway. Plant Journal 28: 201–208.

Hernández-García, J., Sun, R., Serrano-Mislata, A., Inoue, K., Vargas-Chávez, C., Esteve-Bruna, D., Arbona, V., Yamaoka, S., Nishihama, R., Kohchi, T., and Blázquez, M.A. (2021). Coordination between growth and stress responses by DELLA in the liverwort Marchantia polymorpha. Current Biology.

Hirano, K. et al. (2007). The GID1-mediated gibberellin perception mechanism is conserved in the lycophyte Selaginella moellendorffii but not in the bryophyte *Physcomitrella patens*. The Plant Cell 19: 3058–3079.

Hisamatsu, T., King, R.W., Helliwell, C. a, and Koshioka, M. (2005). The involvement of gibberellin 20-oxidase genes in phytochrome-regulated petiole elongation of Arabidopsis. Plant Physiology 138: 1106–1116.

Hisanaga, T., Okahashi, K., Yamaoka, S., Kajiwara, T., Nishihama, R., Shimamura, M., Yamato, K.T., Bowman, J.L., Kohchi, T., and Nakajima, K. (2019). A cis-acting bidirectional transcription switch controls sexual dimorphism in the liverwort. The EMBO Journal 38: 1–12.

Hoang, D.T., Chernomor, O., von Haeseler, A., Minh, B.Q., and Vinh, L.S. (2018). UFBoot2: Improving the Ultrafast Bootstrap Approximation. Molecular Biology and Evolution 35: 518–522.

Holmes, M.G. and Smith, H. (1975). The function of phytochrome in plants growing in the natural environment. Nature 254: 512–514.

Hornych, O., Testo, W.L., Sessa, E.B., Watkins, J.E., Campany, C.E., Pittermann, J., and Ekrt, L. (2021). Insights into the evolutionary history and widespread occurrence of antheridiogen systems in ferns. New Phytologist 229: 607–619.

Inoue, K., Nishihama, R., Araki, T., and Kohchi, T. (2019). Reproductive induction is a far-red high irradiance response that is mediated by phytochrome and PHYTOCHROME INTERACTING FACTOR in Marchantia polymorpha. Plant and Cell Physiology 60: 1136– 1145.

Inoue, K., Nishihama, R., Kataoka, H., Hosaka, M., Manabe, R., Nomoto, M., Tada, Y., Ishizaki, K., and Kohchi, T. (2016). Phytochrome signaling is mediated by PHYTOCHROME INTERACTING FACTOR in the liverwort *Marchantia polymorpha*. The Plant Cell 28: 1406–1421.

Ishizaki, K., Chiyoda, S., Yamato, K.T., and Kohchi, T. (2008). *Agrobacterium*-mediated transformation of the haploid liverwort *Marchantia polymorpha* L., an emerging model for plant biology. Plant and Cell Physiology 49: 1084–1091.

Ishizaki, K., Nishihama, R., Ueda, M., Inoue, K., Ishida, S., Nishimura, Y., Shikanai, T., and Kohchi, T. (2015). Development of Gateway binary vector series with four different selection markers for the liverwort *Marchantia polymorpha*. PLoS ONE 10: e0138876.

Iwasaki, M. et al. (2021). Identification of the sex-determining factor in the liverwort Marchantia polymorpha reveals unique evolution of sex chromosomes in a haploid system. Current Biology 31: 5522–5532.e7.

Jia, Q., Brown, R., Köllner, T.G., Fu, J., Chen, X., Wong, G.K.S., Gershenzon, J., Peters, R.J., and Chen, F. (2022). Origin and early evolution of the plant terpene synthase family. Proceedings of the National Academy of Sciences of the United States of America 119: 1–9.

Kalyaanamoorthy, S., Minh, B.Q., Wong, T.K.F., von Haeseler, A., and Jermiin, L.S. (2017). ModelFinder: fast model selection for accurate phylogenetic estimates. Nat Methods 14: 587–589.

Katoh, K. and Standley, D.M. (2013). MAFFT multiple sequence alignment software version 7: Improvements in performance and usability. Molecular Biology and Evolution 30: 772– 780.

Katsumata, T., Hasegawa, A., Fujiwara, T., Komatsu, T., Notomi, M., Abe, H., Natsume, M., and Kawaide, H. (2008). Arabidopsis CYP85A2 Catalyzes Lactonization Reactions in the Biosynthesis of 2-Deoxy-7-oxalactone Brassinosteroids. Bioscience, Biotechnology, and Biochemistry 72: 2110–2117.

Kohnen, M.V., Schmid-Siegert, E., Trevisan, M., Petrolati, L.A., Sénéchal, F., Müller-Moulé, P., Maloof, J., Xenarios, I., and Fankhauser, C. (2016). Neighbor Detection Induces Organ-Specific Transcriptomes, Revealing Patterns Underlying Hypocotyl-Specific Growth. Plant Cell 28: 2889–2904.

Koide, E., Suetsugu, N., Iwano, M., Gotoh, E., Nomura, Y., Stolze, S.C., Nakagami, H., Kohchi, T., and Nishihama, R. (2020). Regulation of Photosynthetic Carbohydrate Metabolism by a Raf-Like Kinase in the Liverwort Marchantia polymorpha. Plant and Cell Physiology 61: 631–643.

Koornneef, M. and van der Veen, J.H. (1980). Induction and analysis of gibberellin sensitive mutants in *Arabidopsis thaliana* (L.) heynh. Theoretical and Applied Genetics 58: 257– 263.

Krijthe, J.H. (2015). Rtsne: T-Distributed Stochastic Neighbor Embedding using a Barnes-Hut Implementation.

Kubota, A., Ishizaki, K., Hosaka, M., and Kohchi, T. (2013). Efficient *Agrobacterium*-mediated transformation of the liverwort *Marchantia polymorpha* using regenerating thalli. Bioscience, Biotechnology and Biochemistry 77: 167–172.

Kumar, S. et al. (2016). Molecular diversity of terpene synthases in the liverwort *Marchantia polymorpha*. The Plant Cell 28: 2632–2650.

Küpers, J.J. et al. (2023). Local light signaling at the leaf tip drives remote differential petiole growth through auxin-gibberellin dynamics. Current Biology 33: 75–85.e5.

Kurihara, D., Mizuta, Y., Nagahara, S., and Higashiyama, T. (2021). ClearSeeAlpha: Advanced Optical Clearing for Whole-Plant Imaging. Plant and Cell Physiology 62: 1302–1310.

Kurosawa, E. (1926). Experimental studies on the nature of the substance excreted by the “bakanae” fungus. Trans. Nat. Hist. Soc. Formosa 16: 213–227.

Li, W., Liu, S.-W., Ma, J.-J., Liu, H.-M., Han, F.-X., Li, Y., and Niu, S.-H. (2020). Gibberellin Signaling Is Required for Far-Red Light-Induced Shoot Elongation in *Pinus tabuliformis* Seedlings. Plant Physiol. 182: 658–668.

Love, M.I., Huber, W., and Anders, S. (2014). Moderated estimation of fold change and dispersion for RNA-seq data with DESeq2. Genome Biology 15: 550.

Michaud, O., Krahmer, J., Galbier, F., Lagier, M., Galvão, V.C., Ince, Y.Ç., Trevisan, M., Knerova, J., Dickinson, P., Hibberd, J.M., Zeeman, S.C., and Fankhauser, C. (2023). Abscisic acid modulates neighbor proximity-induced leaf hyponasty in Arabidopsis. Plant Physiology 191: 542–557.

Minh, B.Q., Schmidt, H.A., Chernomor, O., Schrempf, D., Woodhams, M.D., von Haeseler, A., and Lanfear, R. (2020). IQ-TREE 2: New Models and Efficient Methods for Phylogenetic Inference in the Genomic Era. Molecular Biology and Evolution 37: 1530–1534.

Miyazaki, S., Hara, M., Ito, S., Tanaka, K., Asami, T., Hayashi, K., Kawaide, H., and Nakajima, M. (2018). An Ancestral Gibberellin in a Moss Physcomitrella patens. Molecular Plant 11: 1097–1100.

Miyazaki, S., Katsumata, T., Natsume, M., and Kawaide, H. (2011). The CYP701B1 of *Physcomitrella patens* is an ent-kaurene oxidase that resists inhibition by uniconazole-P. FEBS Letters 585: 1879–1883.

Miyazaki, S., Nakajima, M., and Kawaide, H. (2015). Hormonal diterpenoids derived from ent-kaurenoic acid are involved in the blue-light avoidance response of Physcomitrella patens. Plant Signaling and Behavior 10: 1–4.

Miyazaki, S., Toyoshima, H., Natsume, M., Nakajima, M., and Kawaide, H. (2014). Blue-light irradiation up-regulates the *ent*-kaurene synthase gene and affects the avoidance response of protonemal growth in *Physcomitrella patens*. Planta 240: 117–124.

Morgan, D.C. and Smith, H. (1979). A systematic relationship between phytochrome-controlled development and species habitat, for plants grown in simulated natural radiation. Planta 145: 253–258.

Morgan, D.C. and Smith, H. (1978). The relationship between phytochrome-photoequilibrium and Development in light grown Chenopodium album L. Planta 142: 187–193.

Motohashi, K. (2015). A simple and efficient seamless DNA cloning method using SLiCE from Escherichia coli laboratory strains and its application to SLiP site-directed mutagenesis. BMC Biotechnol 15: 47.

Näf, U., Nakanishi, K., and Endo, M. (1975). On the Physiology and Chemistry of Fern Antheridiogens. Botanical Review 41: 315–359.

Nelissen, H., Rymen, B., Jikumaru, Y., Demuynck, K., Van Lijsebettens, M., Kamiya, Y., Inzé, D., and Beemster, G.T.S. (2012). A Local Maximum in Gibberellin Levels Regulates Maize Leaf Growth by Spatial Control of Cell Division. Current Biology 22: 1183–1187.

Ninnemann, H. and Halbsguth, W. (1965). Rolle des Phytochroms beim Etiolement von Marchantia polymorpha. Naturwissenschaften 52: 110–111.

Nomura, T., Magome, H., Hanada, A., Takeda-Kamiya, N., Mander, L.N., Kamiya, Y., and Yamaguchi, S. (2013). Functional analysis of *Arabidopsis* CYP714A1 and CYP714A2 reveals that they are distinct gibberellin modification enzymes. Plant and Cell Physiology 54: 1837–1851.

Ogawa, M., Hanada, A., Yamauchi, Y., Kuwahara, A., Kamiya, Y., and Yamaguchi, S. (2003). Gibberellin Biosynthesis and Response during Arabidopsis Seed Germination. The Plant Cell 15: 1591–1604.

Patro, R., Duggal, G., Love, M.I., Irizarry, R.A., and Kingsford, C. (2017). Salmon provides fast and bias-aware quantification of transcript expression. Nature Methods 14: 417–419.

Pluthero, F.G. (1993). Rapid purification of high-activity Taq DNA polymerase. Nucleic Acids Res 21: 4850–4851.

Rojas, M.C., Hedden, P., Gaskin, P., and Tudzynski, B. (2001). The *P450–1* gene of *Gibberella fujikuroi* encodes a multifunctional enzyme in gibberellin biosynthesis. Proc. Natl. Acad. Sci. U.S.A. 98: 5838–5843.

Sakamoto, Y., Ishimoto, A., Sakai, Y., Sato, M., Nishihama, R., Abe, K., Sano, Y., Furuichi, T., Tsuji, H., Kohchi, T., and Matsunaga, S. (2022). Improved clearing method contributes to deep imaging of plant organs. Commun Biol 5: 12.

Schindelin, J. et al. (2012). Fiji: an open-source platform for biological-image analysis. Nat Methods 9: 676–682.

Schmidt, U., Weigert, M., Broaddus, C., and Myers, G. (2018). Cell Detection with Star-Convex Polygons. In Medical Image Computing and Computer Assisted Intervention – MICCAI 2018, A.F. Frangi, J.A. Schnabel, C. Davatzikos, C. Alberola-López, and G. Fichtinger, eds, Lecture Notes in Computer Science. (Springer International Publishing: Cham), pp. 265–273.

Schneller, J.J. (2008). Antheridiogens. In Biology and Evolution of Ferns and Lycophytes, T.A. Ranker and C.H. Haufler, eds (Cambridge University Press: New York), pp. 134–158.

Sessa, G., Carabelli, M., Sassi, M., Ciolfi, A., Possenti, M., Mittempergher, F., Becker, J., Morelli, G., and Ruberti, I. (2005). A dynamic balance between gene activation and repression regulates the shade avoidance response in *Arabidopsis*. Genes Dev. 19: 2811–2815.

Shimamura, M. (2016). *Marchantia polymorpha*: Taxonomy, phylogeny and morphology of a model system. Plant and Cell Physiology 57: 230–256.

Sponsel, V.M. (2016). Signal achievements in gibberellin research: the second half-century. Annual Plant Reviews: The Gibberellins 49: 1–36.

Streubel, S., Deiber, S., Rötzer, J., Mosiolek, M., Jandrasits, K., and Dolan, L. (2023). Meristem dormancy in Marchantia polymorpha is regulated by a liverwort-specific miRNA and a clade III SPL gene. Current Biology: S0960982222019996.

Sugano, S.S., Nishihama, R., Shirakawa, M., Takagi, J., Matsuda, Y., Ishida, S., Shimada, T., Hara-Nishimura, I., Osakabe, K., and Kohchi, T. (2018). Efficient CRISPR/Cas9-based genome editing and its application to conditional genetic analysis in Marchantia polymorpha. PLoS ONE 13: e0205117.

Sun, T. and Kamiya, Y. (1994). The Arabidopsis *GA1* locus encodes the cyclase ent-kaurene synthetase A of gibberellin biosynthesis. the Plant Cell 6: 1509–1518.

Talon, M., Koornneef, M., and Zeevaart, J.A.D. (1990). Endogenous gibberellins in Arabidopsis thaliana and possible steps blocked in the biosynthetic pathways of the semidwarf ga4 and ga5 mutants. Proceedings of the National Academy of Sciences of the United States of America 87: 7983–7987.

Tanaka, J., Yano, K., Aya, K., Hirano, K., Takehara, S., Koketsu, E., Ordonio, R.L., Park, S.-H., Nakajima, M., Ueguchi-Tanaka, M., and Matsuoka, M. (2014). Antheridiogen determines sex in ferns via a spatiotemporally split gibberellin synthesis pathway. Science 346: 469– 473.

Toyomasu, T., Kawaide, H., Mitsuhashi, W., Inoue, Y., and Kamiya, Y. (1998). Phytochrome regulates gibberellin biosynthesis during germination of photoblastic lettuce seeds. Plant Physiology 118: 1517–1523.

Ubeda-Tomás, S., Federici, F., Casimiro, I., Beemster, G.T.S., Bhalerao, R., Swarup, R., Doerner, P., Haseloff, J., and Bennett, M.J. (2009). Gibberellin Signaling in the Endodermis Controls Arabidopsis Root Meristem Size. Current Biology 19: 1194–1199.

Ubeda-Tomás, S., Swarup, R., Coates, J., Swarup, K., Laplaze, L., Beemster, G.T.S., Hedden, P., Bhalerao, R., and Bennett, M.J. (2008). Root growth in Arabidopsis requires gibberellin/DELLA signalling in the endodermis. Nat Cell Biol 10: 625–628.

Urrutia, O., Hedden, P., and Rojas, M.C. (2001). Monooxygenases involved in GA12 and GA14 synthesis in Gibberella fujikuroi. Phytochemistry 56: 505–511.

Van Tuinen, A., Peters, A.H.L.J., Kendrick, R.E., Zeevaart, J. a. D., and Koornneef, M. (1999). Characterisation of the procera mutant of tomato and the interaction of gibberellins with end-of-day far-red light treatments. Physiologia Plantarum 106: 121–128.

Wenzel, C.L., Williamson, R.E., and Wasteneys, G.O. (2000). Gibberellin-Induced Changes in Growth Anisotropy Precede Gibberellin-Dependent Changes in Cortical Microtubule Orientation in Developing Epidermal Cells of Barley Leaves. Kinematic and Cytological Studies on a Gibberellin-Responsive Dwarf Mutant, M489. Plant Physiology 124: 813– 822.

Whitelam, G.C. and Johnson, C.B. (1982). PHOTOMORPHOGENESIS IN IMPATIENS PARVIFLORA AND OTHER PLANT SPECIES UNDER SIMULATED NATURAL CANOPY RADIATIONS. New Phytol 90: 611–618.

Whitelam, G.C. and Smith, H. (1991). Retention of Phytochrome-Mediated Shade Avoidance Responses in Phytochrome-Deficient Mutants of Arabidopsis, Cucumber and Tomato. Journal of Plant Physiology 139: 119–125.

Williams, J., Phillips, A.L., Gaskin, P., and Hedden, P. (1998). Function and Substrate Specificity of the Gibberellin 3β-Hydroxylase Encoded by the Arabidopsis GA4Gene. Plant Physiology 117: 559–563.

Xu, S., Li, L., Luo, X., Chen, M., Tang, W., Zhan, L., Dai, Z., Lam, T.T., Guan, Y., and Yu, G. (2022). Ggtree: A serialized data object for visualization of a phylogenetic tree and annotation data. iMeta 1: e56.

Yamaguchi, S., Smith, M.W., Brown, R.G.S., Kamiya, Y., and Sun Tai-ping (1998a). Phytochrome Regulation and Differential Expression of Gibberellin 3β-Hydroxylase Genes in Germinating Arabidopsis Seeds. The Plant Cell 10: 2115–2126.

Yamaguchi, S., Sun, T., Kawaide, H., and Kamiya, Y. (1998b). The *GA2* locus of *Arabidopsis thaliana* encodes *ent*-kaurene synthase of gibberellin biosynthesis. Plant Physiology 116: 1271–1278.

Yamane, H. (1998). Fern Antheridiogens. International Review of Cytology 184: 1–32.

Yamaoka, S. et al. (2018). Generative Cell Specification Requires Transcription Factors Evolutionarily Conserved in Land Plants. Current Biology 28: 479–486.e5.

Yamauchi, T., Oyama, N., Yamane, H., Murofushi, N., Schraudolf, H., Pour, M., Furber, M., and Mander, L.N. (1996). Identification of Antheridiogens in Lygodium circinnatum and Lygodium flexuosum. Plant Physiology 111: 741–745.

Yoshida, H., Takehara, S., Mori, M., Ordonio, R.L., and Matsuoka, M. (2020). Evolution of GA metabolic enzymes in land plants. Plant and Cell Physiology 61: 1919–1934.

Yu, G. (2022). Data Integration, Manipulation and Visualization of Phylogenetic Trees (Chapman and Hall/CRC: New York).

Yu, G. (2020a). Gene ontology semantic similarity analysis using GOSemSim. In Methods in Molecular Biology (Humana Press Inc.), pp. 207–215.

Yu, G. (2020b). Using ggtree to Visualize Data on Tree-Like Structures. Current Protocols in Bioinformatics 69: e96.

Yu, G., Lam, T.T.-Y., Zhu, H., and Guan, Y. (2018). Two Methods for Mapping and Visualizing Associated Data on Phylogeny Using Ggtree. Molecular Biology and Evolution 35: 3041– 3043.

Yu, G., Li, F., Qin, Y., Bo, X., Wu, Y., and Wang, S. (2010). GOSemSim: an R package for measuring semantic similarity among GO terms and gene products. Bioinformatics 26: 976–978.

Yu, G., Smith, D.K., Zhu, H., Guan, Y., and Lam, T.T.-Y. (2017). ggtree: an r package for visualization and annotation of phylogenetic trees with their covariates and other associated data. Methods in Ecology and Evolution 8: 28–36.

Zhang, Y., Werling, U., and Edelmann, W. (2012). SLiCE: A novel bacterial cell extract-based DNA cloning method. Nucleic Acids Research 40: 1–10.

