## Supplemental Figures for "Biosynthesis of gibberellin-related compounds modulates far-red light responses in the liverwort *Marchantia polymorpha*"

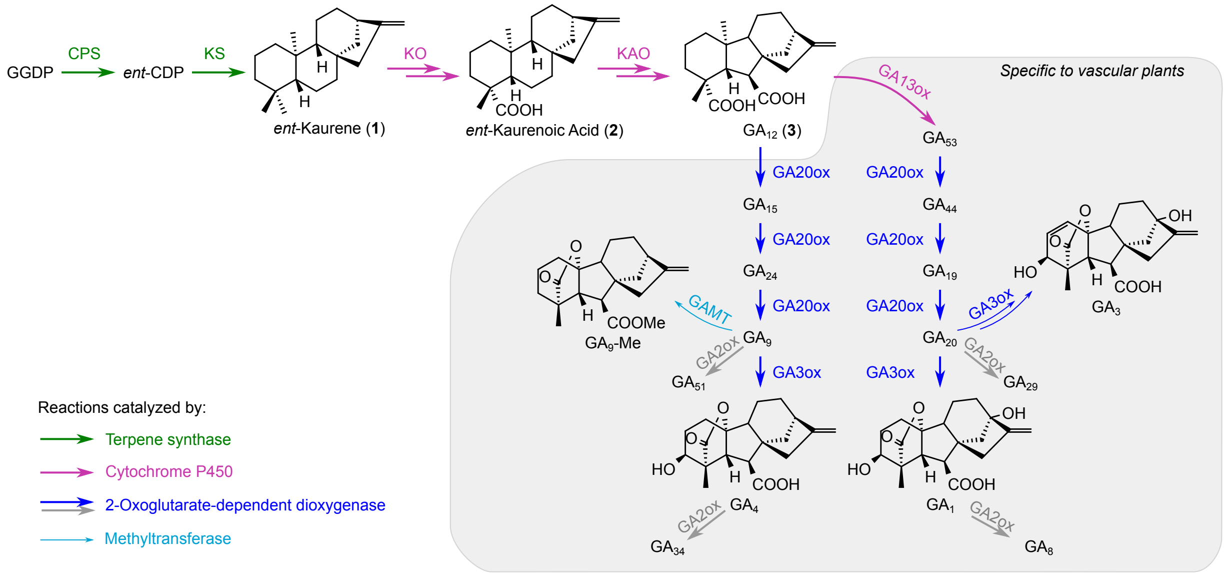


**Supplemental Figure 1** Gibberellin (GA) biosynthesis pathway in vascular plants, showing compounds analyzed or used for treatment in this research. GGDP, geranylgeranyl diphosphate; *ent*-CDP, *ent*-copalyl diphosphate; CPS, *ent*-copalyl diphosphate synthase; KS, *ent*-kaurene synthase; KO, *ent*-kaurene oxidase; KAO, *ent*-kaurenoic acid oxidase. GA13ox, GA 13-oxidase; GA20ox, GA 20-oxidase; GA3ox, GA 3-oxidase; GA2ox, GA 2-oxidase; GAMT, GA methyltransferase. Deactivation steps catalyzed by GA2ox was shown with grey arrows. $GA_3$ is a major product from the fungus *Fusarium fujikuroi*, but could also be produced from $GA_{20}$ by angiosperms like *Zea mays* or *Marah macrocarpa* (Fujioka et al., 1990; Albone et al., 1990). GAMT and GA2ox could act on broader ranges of substrates than shown in the figure, but reactions irrelevant to compounds used in this research were omitted. Related to Figure 3.

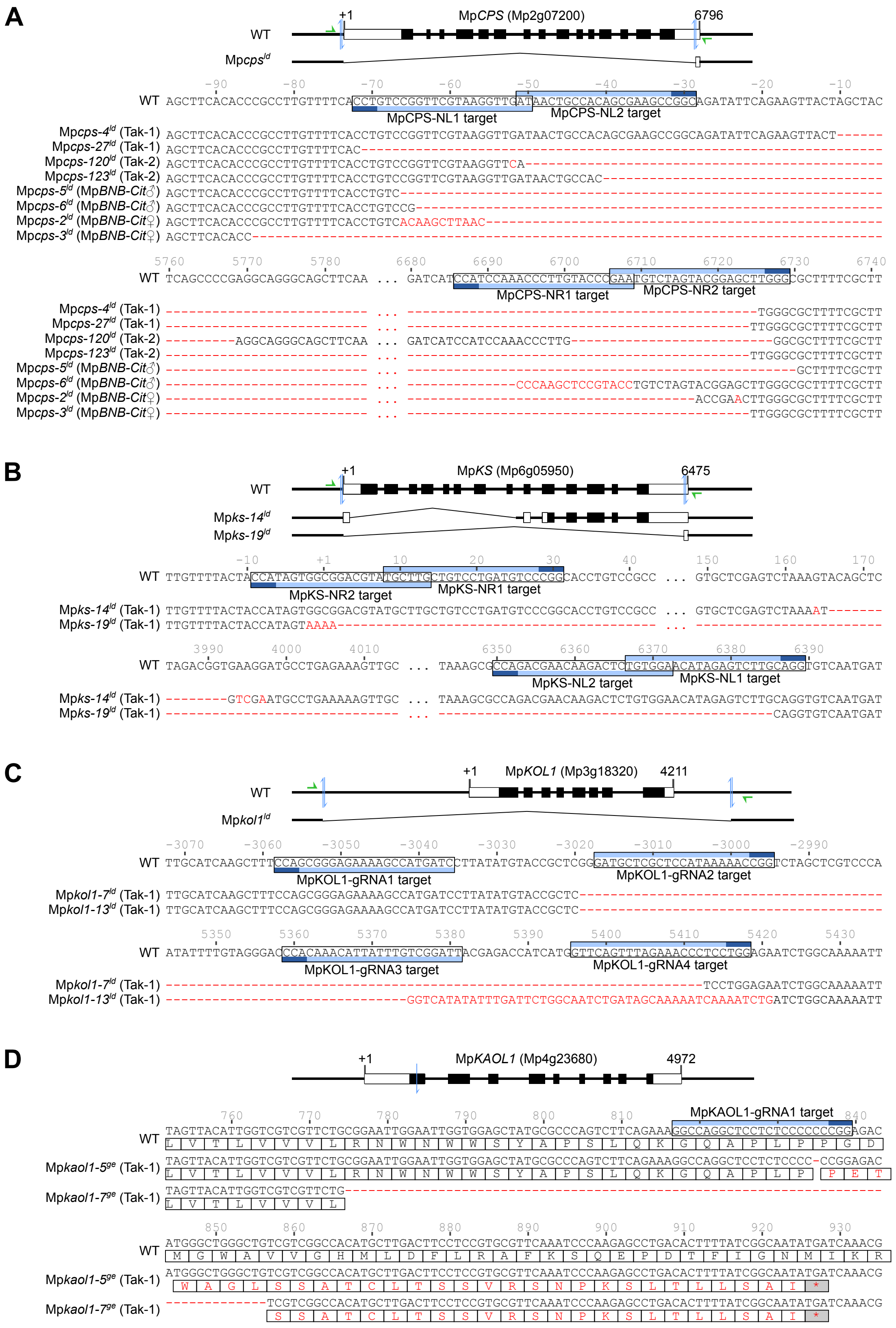


**Supplemental Figure 2** Genotype information for Mp*cps*$^{ld}$ (A), Mp*ks*$^{ld}$ (B), Mp*kol1*$^{ld}$ (C), and Mp*kaol1*$^{ge}$ (D) mutants. WT refers to reference sequences from MpTak_v6.1 genome assembly. In the schematic presentations of genomic structures, white and black rectangles represent untranslated and coding regions of exons, respectively. Targets of guide RNAs are indicated by blue arrows in the scheme and frames in the sequence (dark blue: the protospacer adjacent motif). Green arrows indicate the binding sites of genotyping primers. Indels and substitutions are shown in red letters. Numbers above the sequences indicate positions relative to the transcription start sites (+1). Putative protein translations are shown with framed letters in D, and the premature stop codons are marked with asterisks (*). Related to Figures 1 and 4.

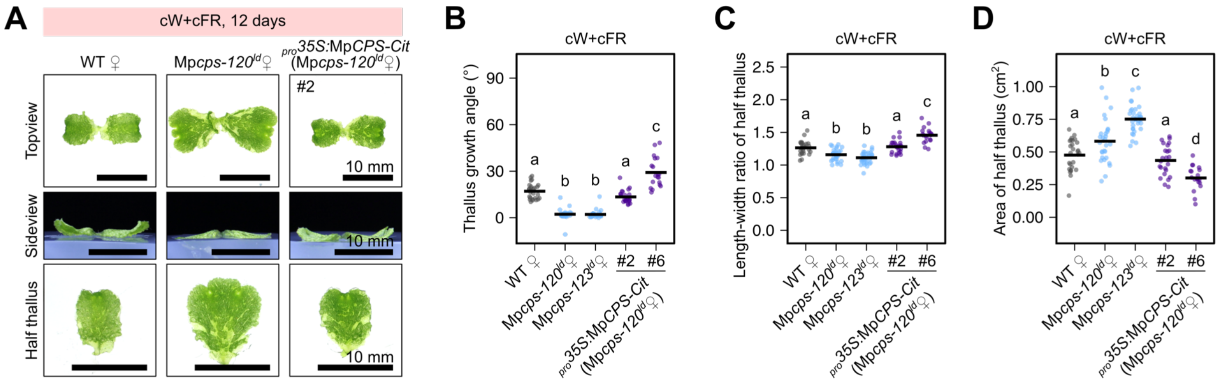


**Supplemental Figure 3** Thallus morphology of female Mp*cps*$^{ld}$ mutants in Tak-2 background under cW+cFR. A, Photos of 12-day-old plants grown from gemmae under cW+cFR. Bars = 10 mm. WT♀ refers to Tak-2 wild-type plants. B-D, Measurements of the thallus growth angle (B), length-width ratio (C) and thallus area (D) from half plants shown in A. Horizontal lines represent mean values, and letters represent multiple comparisons with non-pooled Welch's *t*-test and B-H adjustment (adjusted $p<0.05$ for non-overlapping letters, n=19-34). Related to Figure 1.

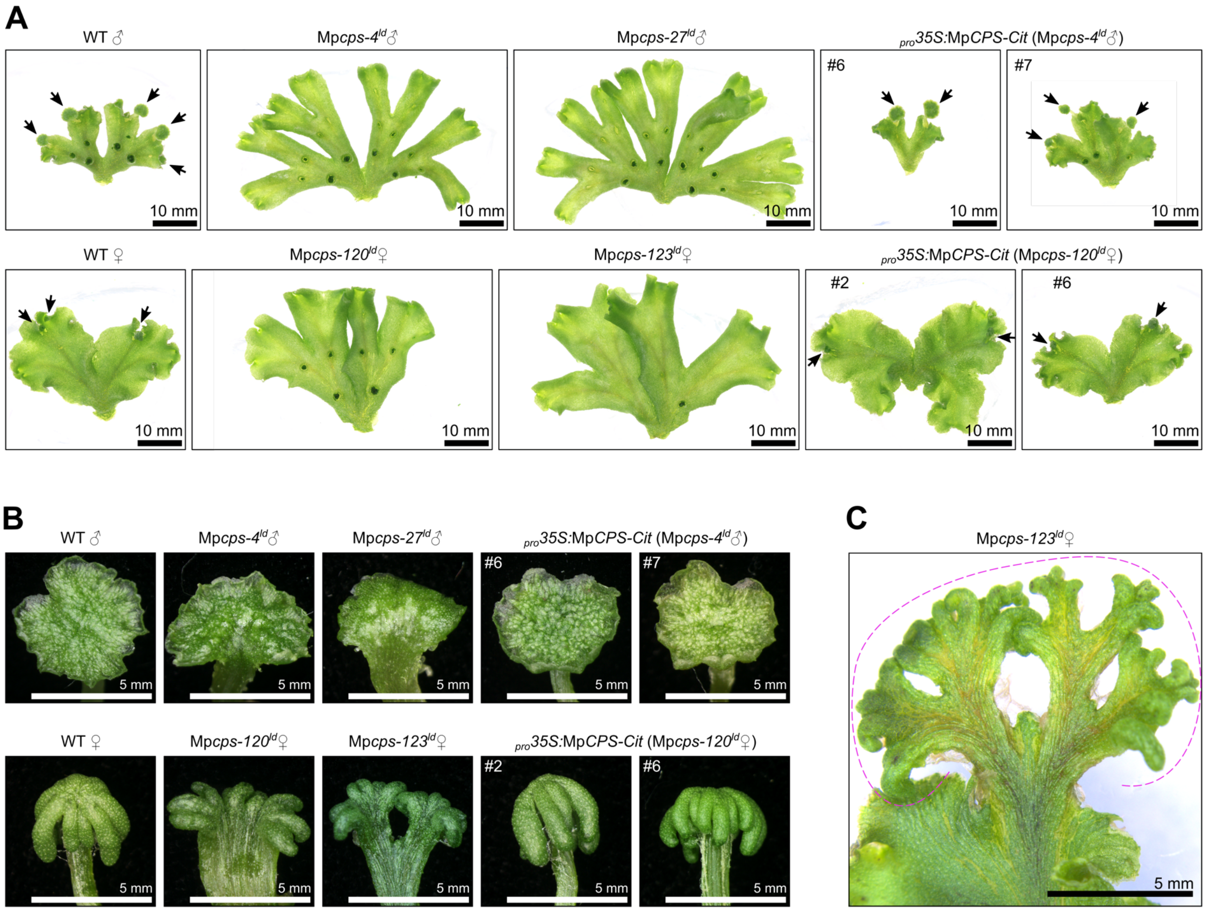


**Supplemental Figure 4** Morphology of Mp*cps$^{ld}$* plants during gametangiophore formation in aseptic culture. A, Morphology of half thalli cultured under cW for 7 days, then under cW+cFR for 16 days. Arrows indicate gametangiophores visible by the naked eye. WT♂ and WT♀ refer to Tak-1 and Tak-2 wild-type plants, respectively. B, Morphology of young gametangiophores from aseptic culture. C, An extreme example of Mp*cps$^{ld}$* female gametangiophore with indeterminate bifurcation. The dashed line indicates the whole structure equivalent to a single gametangiophore. Related to Figure 2.

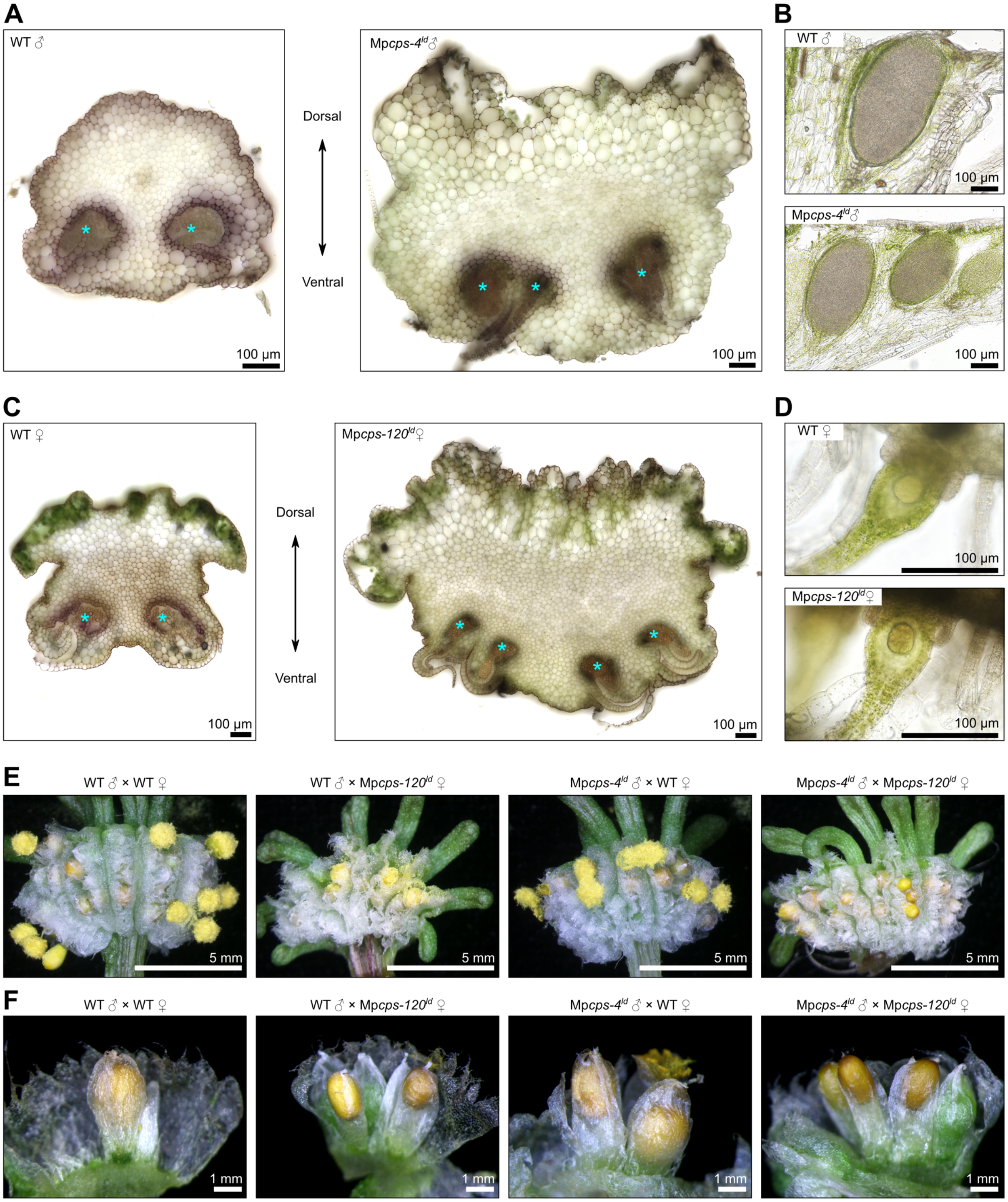


**Supplemental Figure 5** Sections of gametangiophores in wild-type and Mp*cps*$^{ld}$ plants, as well as the fertility test. A and C, Transverse sections of antheridiophore (A) or archegoniophore (C) stalks. Asterisks indicate bundles of pegged rhizoids. Thickness: 200 µm. B and D, longitudinal sections of antheridiophore (B) or archegoniophore (D) receptacles, showing the antheridium (B) or the egg cell in archegonium (D). Thickness: 70 µm. E-F, Mature sporangia from crossing experiments with various combinations among wild-type and Mp*cps*$^{ld}$ plants. Related to Figure 2.

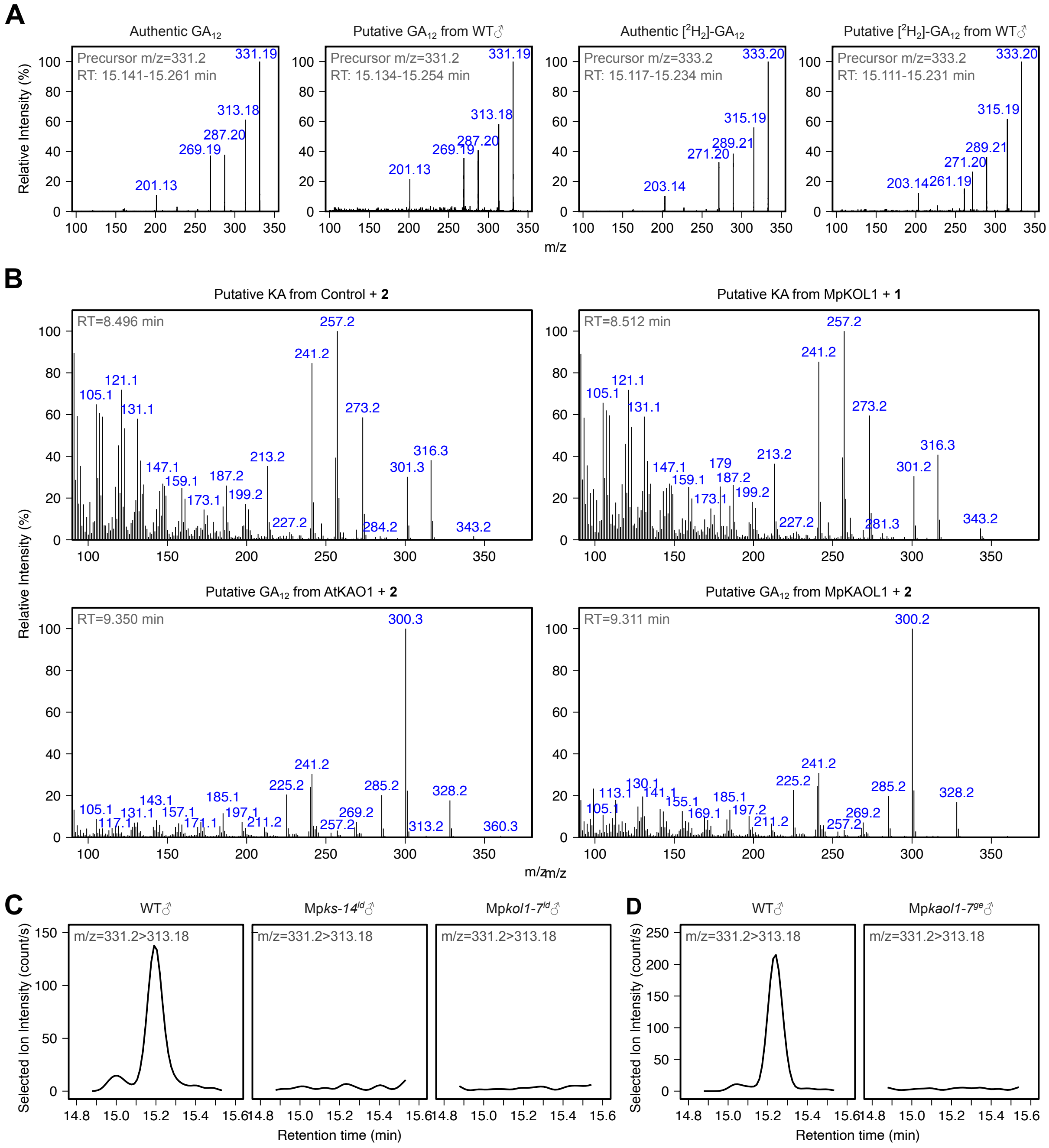


**Supplemental Figure 6** LC-MS/MS and GC-MS profiles for KA and $GA_{12}$ detection. A, Mass spectra of $GA_{12}$ and $[^2H_2]$-$GA_{12}$ in wild-type *M. polymorpha* samples were identical to those of authentic compounds. Detected by LC-MS/MS in the product-ion scanning mode. B, Mass spectra of KA and $GA_{12}$ detected by GC-MS in *P. pastoris* cultures expressing MpKOL1 or MpKAOL1, compared to the positive controls. C-D, Selected ion chromatograph showing $GA_{12}$ deficiency in Mp*ks-14*$^{ld}$, Mp*kol1-7*$^{ld}$ or Mp*kaol1-7*$^{ge}$, detected by LC-MS/MS with the multiple reaction monitoring (MRM) mode. Related to Figures 3 and 4.

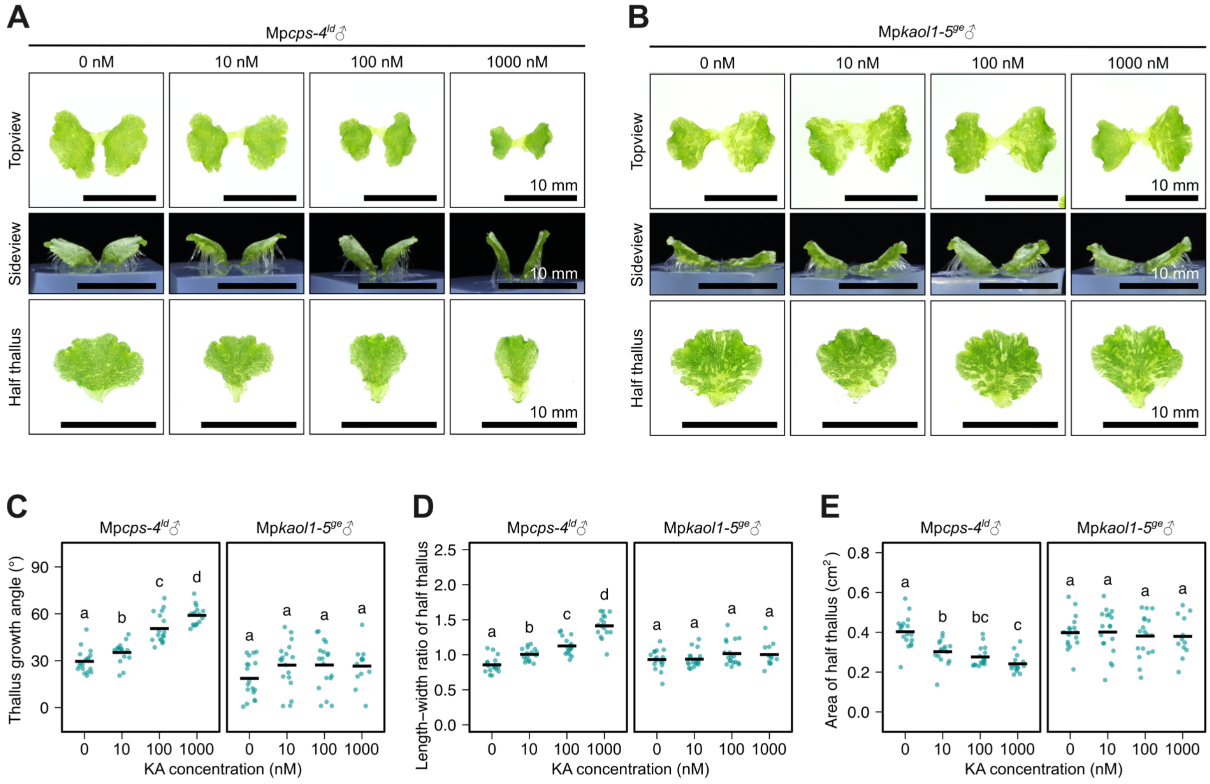


**Supplemental Figure 7** Response of Mp*cps-4*$^{ld}$ and Mp*kaol1-5*$^{ge}$ to different concentrations of KA. A-B, Morphology of 12-day-old plants cultured under cW+cFR from gemmae with different concentrations of KA. Bars = 10 mm. C-E, Measurements of the thallus growth angle (C), length-width ratio (D) and thallus area (E) from half plants shown in A-B. Horizontal lines represent mean values, and letters represent multiple comparisons with non-pooled Welch's *t*-test and B-H adjustment (adjusted $p<0.05$ for non-overlapping letters, n=12-18). Related to Figures 3 and 4.

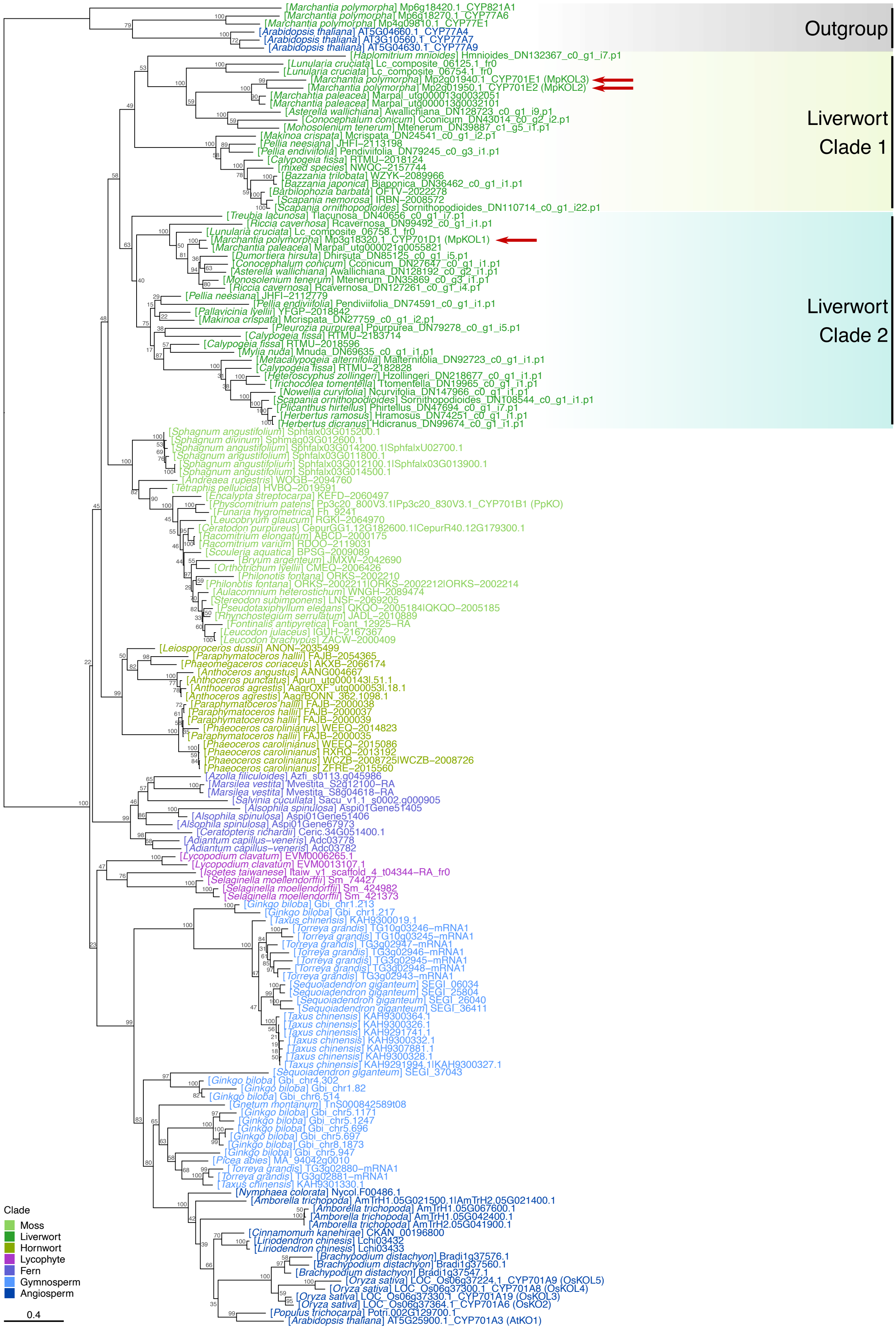

Outgroup
Liverwort Clade 1
Liverwort Clade 2
[Marchantia polymorpha] Mp6g18420.1_CYP821A1
[Marchantia polymorpha] Mp6g18270.1_CYP77A6
[Marchantia polymorpha] Mp4g09810.1_CYP77E1
[Arabidopsis thaliana] AT5G04660.1_CYP77A4
[Arabidopsis thaliana] AT3G10560.1_CYP77A7
[Arabidopsis thaliana] AT5G04630.1_CYP77A9
[Haplomitrium mnioides] Hmnioides_DN132367_c0_g1_i7.p1
[Lunularia cruciata] Lc_composite_06125.1_fr0
[Lunularia cruciata] Lc_composite_06754.1_fr0
[Marchantia polymorpha] Mp2g01940.1_CYP701E1 (MpKOL3)
[Marchantia polymorpha] Mp2g01950.1_CYP701E2 (MpKOL2)
[Marchantia paleacea] Marpal_utg000013g0032051
[Marchantia paleacea] Marpal_utg000013g0032101
[Asterella wallichiana] Awallichiana_DN128723_c0_g1_i9.p1
[Conocephalum conicum] Cconicum_DN43014_c0_g2_i2.p1
[Monosolenium tenerum] Mtenerum_DN39887_c1_g5_i1.p1
[Makinoa crispata] Mcrispata_DN24541_c0_g1_i2.p1
[Pellia neesiana] JHFI–2113198
[Pellia endiviifolia] Pendiviifolia_DN79245_c0_g3_i1.p1
[Calypogeia fissa] RTMU–2018124
[mixed species] NWQC–2157744
[Bazzania trilobata] WZYK–2089966
[Bazzania japonica] Bjaponica_DN36462_c0_g1_i1.p1
[Barbilophozia barbata] OFTV–2022278
[Scapania nemorosa] IRBN–2008572
[Scapania ornithopodioides] Sornithopodioides_DN110714_c0_g1_i22.p1
[Treubia lacunosa] Tlacunosa_DN40656_c0_g1_i7.p1
[Riccia cavernosa] Rcavernosa_DN99492_c0_g1_i1.p1
[Lunularia cruciata] Lc_composite_06758.1_fr0
[Marchantia polymorpha] Mp3g18320.1_CYP701D1 (MpKOL1)
[Marchantia paleacea] Marpal_utg000021g0055821
[Dumortiera hirsuta] Dhirsuta_DN85125_c0_g1_i5.p1
[Conocephalum conicum] Cconicum_DN27647_c0_g1_i1.p1
[Asterella wallichiana] Awallichiana_DN128192_c0_g2_i1.p1
[Monosolenium tenerum] Mtenerum_DN35869_c0_g3_i1.p1
[Riccia cavernosa] Rcavernosa_DN127261_c0_g1_i4.p1
[Pellia neesiana] JHFI–2112779
[Pellia endiviifolia] Pendiviifolia_DN74591_c0_g1_i1.p1
[Pallavicinia lyellii] YFGP–2018842
[Makinoa crispata] Mcrispata_DN27759_c0_g1_i2.p1
[Pleurozia purpurea] Ppurpurea_DN79278_c0_g1_i5.p1
[Calypogeia fissa] RTMU–2183714
[Calypogeia fissa] RTMU–2018596
[Mylia nuda] Mnuda_DN69635_c0_g1_i1.p1
[Metacalypogeia alternifolia] Malternifolia_DN92723_c0_g1_i1.p1
[Calypogeia fissa] RTMU–2182828
[Heteroscyphus zollingeri] Hzollingeri_DN218677_c0_g1_i1.p1
[Trichocolea tomentella] Ttomentella_DN19965_c0_g1_i1.p1
[Nowellia curvifolia] Ncurvifolia_DN147966_c0_g1_i1.p1
[Scapania ornithopodioides] Sornithopodioides_DN108544_c0_g1_i1.p1
[Plicanthus hirtellus] Phirtellus_DN47694_c0_g1_i7.p1
[Herbertus ramosus] Hramosus_DN74251_c0_g1_i1.p1
[Herbertus dicranus] Hdicranus_DN99674_c0_g1_i1.p1
[Sphagnum angustifolium] Sphfalx03G015200.1
[Sphagnum divinum] Sphmag03G012600.1
[Sphagnum angustifolium] Sphfalx03G014200.1|SphfalxU02700.1
[Sphagnum angustifolium] Sphfalx03G011800.1
[Sphagnum angustifolium] Sphfalx03G012100.1|Sphfalx03G013900.1
[Sphagnum angustifolium] Sphfalx03G014500.1
[Andreaea rupestris] WOGB–2094760
[Tetraphis pellucida] HVBQ–2019591
[Encalypta streptocarpa] KEFD–2060497
[Physcomitrium patens] Pp3c20_800V3.1|Pp3c20_830V3.1_CYP701B1 (PpKO)
[Funaria hygrometrica] Fh_9241
[Leucobryum glaucum] RGKI–2064970
[Ceratodon purpureus] CepurGG1.12G182600.1|CepurR40.12G179300.1
[Racomitrium elongatum] ABCD–2000175
[Racomitrium varium] RDOO–2119031
[Scouleria aquatica] BPSG–2009089
[Bryum argenteum] JMXW–2042690
[Orthotrichum lyellii] CMEQ–2006426
[Philonotis fontana] ORKS–2002210
[Philonotis fontana] ORKS–2002211|ORKS–2002212|ORKS–2002214
[Aulacomnium heterostichum] WNGH–2089474
[Stereodon subimponens] LNSF–2069205
[Pseudotaxiphyllum elegans] QKQO–2005184|QKQO–2005185
[Rhynchostegium serrulatum] JADL–2010889
[Fontinalis antipyretica] Foant_12925–RA
[Leucodon julaceus] IGUH–2167367
[Leucodon brachypus] ZACW–2000409
[Leiosporoceros dussii] ANON–2035499
[Paraphymatoceros hallii] FAJB–2054365
[Phaeomegaceros coriaceus] AKXB–2066174
[Anthoceros angustus] AANG004667
[Anthoceros punctatus] Apun_utg000143l.51.1
[Anthoceros agrestis] AagrOXF_utg000053l.18.1
[Anthoceros agrestis] AagrBONN_362.1098.1
[Paraphymatoceros hallii] FAJB–2000038
[Paraphymatoceros hallii] FAJB–2000037
[Paraphymatoceros hallii] FAJB–2000039
[Phaeoceros carolinianus] WEEQ–2014823
[Paraphymatoceros hallii] FAJB–2000035
[Phaeoceros carolinianus] WEEQ–2015086
[Phaeoceros carolinianus] RXRQ–2013192
[Phaeoceros carolinianus] WCZB–2008725|WCZB–2008726
[Phaeoceros carolinianus] ZFRE–2015560
[Azolla filiculoides] Azfi_s0113.g045986
[Marsilea vestita] Mvestita_S2g12100–RA
[Marsilea vestita] Mvestita_S8g04618–RA
[Salvinia cucullata] Sacu_v1.1_s0002.g000905
[Alsophila spinulosa] Aspi01Gene51405
[Alsophila spinulosa] Aspi01Gene51406
[Alsophila spinulosa] Aspi01Gene67973
[Ceratopteris richardii] Ceric.34G051400.1
[Adiantum capillus–veneris] Adc03778
[Adiantum capillus–veneris] Adc03782
[Lycopodium clavatum] EVM0006265.1
[Lycopodium clavatum] EVM0013107.1
[Isoetes taiwanese] Itaiw_v1_scaffold_4_t04344–RA_fr0
[Selaginella moellendorffii] Sm_74427
[Selaginella moellendorffii] Sm_424982
[Selaginella moellendorffii] Sm_421373
[Ginkgo biloba] Gbi_chr1.213
[Ginkgo biloba] Gbi_chr1.217
[Taxus chinensis] KAH9300019.1
[Torreya grandis] TG10g03246–mRNA1
[Torreya grandis] TG10g03245–mRNA1
[Torreya grandis] TG3g02947–mRNA1
[Torreya grandis] TG3g02946–mRNA1
[Torreya grandis] TG3g02945–mRNA1
[Torreya grandis] TG3g02948–mRNA1
[Torreya grandis] TG3g02943–mRNA1
[Sequoiadendron giganteum] SEGI_06034
[Sequoiadendron giganteum] SEGI_25804
[Sequoiadendron giganteum] SEGI_26040
[Sequoiadendron giganteum] SEGI_36411
[Taxus chinensis] KAH9300364.1
[Taxus chinensis] KAH9300326.1
[Taxus chinensis] KAH9291741.1
[Taxus chinensis] KAH9300332.1
[Taxus chinensis] KAH9307881.1
[Taxus chinensis] KAH9300328.1
[Taxus chinensis] KAH9291994.1|KAH9300327.1
[Sequoiadendron giganteum] SEGI_37043
[Ginkgo biloba] Gbi_chr4.302
[Ginkgo biloba] Gbi_chr1.82
[Ginkgo biloba] Gbi_chr6.514
[Gnetum montanum] TnS000842589t08
[Ginkgo biloba] Gbi_chr5.1171
[Ginkgo biloba] Gbi_chr5.1247
[Ginkgo biloba] Gbi_chr5.696
[Ginkgo biloba] Gbi_chr5.697
[Ginkgo biloba] Gbi_chr8.1873
[Ginkgo biloba] Gbi_chr5.947
[Picea abies] MA_94042g0010
[Torreya grandis] TG3g02880–mRNA1
[Torreya grandis] TG3g02881–mRNA1
[Taxus chinensis] KAH9301330.1
[Nymphaea colorata] Nycol.F00486.1
[Amborella trichopoda] AmTrH1.05G021500.1|AmTrH2.05G021400.1
[Amborella trichopoda] AmTrH1.05G067600.1
[Amborella trichopoda] AmTrH1.05G042400.1
[Amborella trichopoda] AmTrH2.05G041900.1
[Cinnamomum kanehirae] CKAN_00196800
[Liriodendron chinense] Lchi03432
[Liriodendron chinense] Lchi03433
[Brachypodium distachyon] Bradi1g37576.1
[Brachypodium distachyon] Bradi1g37560.1
[Brachypodium distachyon] Bradi1g37547.1
[Oryza sativa] LOC_Os06g37224.1_CYP701A9 (OsKOL5)
[Oryza sativa] LOC_Os06g37300.1_CYP701A8 (OsKOL4)
[Oryza sativa] LOC_Os06g37330.1_CYP701A19 (OsKOL3)
[Oryza sativa] LOC_Os06g37364.1_CYP701A6 (OsKO2)
[Populus trichocarpa] Potri.002G129700.1
[Arabidopsis thaliana] AT5G25900.1_CYP701A3 (AtKO1)
Clade
Moss
Liverwort
Hornwort
Lycophyte
Fern
Gymnosperm
Angiosperm
0.4

**Supplemental Figure 8** Phylogenetic tree of KO homologs in land plants. Nodes were labelled with percentage support values from 1000 standard non-parametric bootstraps by IQ-TREE 2. *M. polymorpha* proteins of interest were indicated with red arrows. Related to Figure 4.

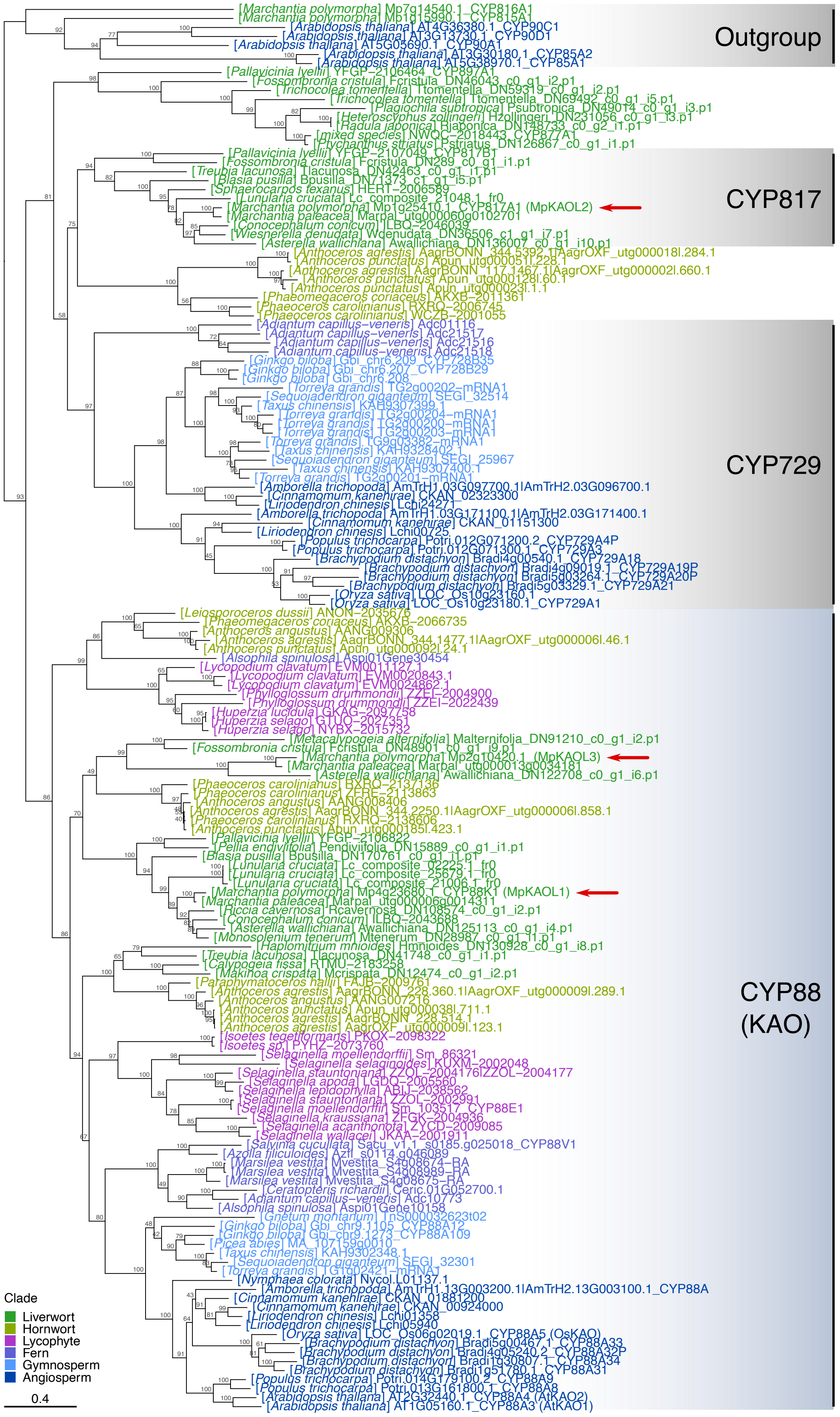
[Marchantia polymorpha] Mp7g14540.1_CYP816A1
[Marchantia polymorpha] Mp1g15990.1_CYP815A1
[Arabidopsis thaliana] AT4G36380.1_CYP90C1
[Arabidopsis thaliana] AT3G13730.1_CYP90D1
[Arabidopsis thaliana] AT5G05690.1_CYP90A1
[Arabidopsis thaliana] AT3G30180.1_CYP85A2
[Arabidopsis thaliana] AT5G38970.1_CYP85A1
Outgroup
[Pallavicinia lyellii] YFGP−2106464_CYP897A1
[Fossombronia cristula] Fcristula_DN46043_c0_g1_i2.p1
[Trichocolea tomentella] Ttomentella_DN59319_c0_g1_i2.p1
[Trichocolea tomentella] Ttomentella_DN69492_c0_g1_i5.p1
[Plagiochila subtropica] Psubtropica_DN49014_c0_g1_i3.p1
[Heteroscyphus zollingeri] Hzollingeri_DN231056_c0_g1_i3.p1
[Radula japonica] Rjaponica_DN148733_c0_g2_i1.p1
[mixed species] NWQC−2018443_CYP877A1
[Ptychanthus striatus] Pstriatus_DN126867_c0_g1_i1.p1
[Pallavicinia lyellii] YFGP−2107049_CYP817B1
[Fossombronia cristula] Fcristula_DN289_c0_g1_i1.p1
[Treubia lacunosa] Tlacunosa_DN42463_c0_g1_i1.p1
[Blasia pusilla] Bpusilla_DN71373_c1_g1_i5.p1
[Sphaerocarpos texanus] HERT−2006589
[Lunularia cruciata] Lc_composite_21048.1_fr0
[Marchantia polymorpha] Mp1g25410.1_CYP817A1 (MpKAOL2)
[Marchantia paleacea] Marpal_utg000060g0102701
[Conocephalum conicum] ILBQ−2046039
[Wiesnerella denudata] Wdenudata_DN36506_c1_g1_i7.p1
[Asterella wallichiana] Awallichiana_DN136007_c0_g1_i10.p1
CYP817
[Anthoceros agrestis] AagrBONN_344.5392.1|AagrOXF_utg000018l.284.1
[Anthoceros punctatus] Apun_utg000051l.228.1
[Anthoceros agrestis] AagrBONN_117.1467.1|AagrOXF_utg000002l.660.1
[Anthoceros punctatus] Apun_utg000128l.60.1
[Anthoceros punctatus] Apun_utg000023l.1.1
[Phaeomegaceros coriaceus] AKXB−2011361
[Phaeoceros carolinianus] RXRQ−2006745
[Phaeoceros carolinianus] WCZB−2001055
[Adiantum capillus−veneris] Adc01116
[Adiantum capillus−veneris] Adc21517
[Adiantum capillus−veneris] Adc21516
[Adiantum capillus−veneris] Adc21518
[Ginkgo biloba] Gbi_chr6.209_CYP728B35
[Ginkgo biloba] Gbi_chr6.207_CYP728B29
[Ginkgo biloba] Gbi_chr6.208
[Torreya grandis] TG2g00202−mRNA1
[Sequoiadendron giganteum] SEGI_32514
[Taxus chinensis] KAH9307399.1
[Torreya grandis] TG2g00204−mRNA1
[Torreya grandis] TG2g00200−mRNA1
[Torreya grandis] TG2g00203−mRNA1
[Torreya grandis] TG9g03382−mRNA1
[Taxus chinensis] KAH9328402.1
[Sequoiadendron giganteum] SEGI_25967
[Taxus chinensis] KAH9307400.1
[Torreya grandis] TG2g00201−mRNA1
[Amborella trichopoda] AmTrH1.03G097700.1|AmTrH2.03G096700.1
[Cinnamomum kanehirae] CKAN_02323300
[Liriodendron chinensis] Lchi24271
[Amborella trichopoda] AmTrH1.03G171100.1|AmTrH2.03G171400.1
[Cinnamomum kanehirae] CKAN_01151300
[Liriodendron chinensis] Lchi00725
[Populus trichocarpa] Potri.012G071200.2_CYP729A4P
[Populus trichocarpa] Potri.012G071300.1_CYP729A3
[Brachypodium distachyon] Bradi4g00540.1_CYP729A18
[Brachypodium distachyon] Bradi4g09019.1_CYP729A19P
[Brachypodium distachyon] Bradi5g03264.1_CYP729A20P
[Brachypodium distachyon] Bradi5g03329.1_CYP729A21
[Oryza sativa] LOC_Os10g23160.1
[Oryza sativa] LOC_Os10g23180.1_CYP729A1
CYP729
[Leiosporoceros dussii] ANON−2035676
[Phaeomegaceros coriaceus] AKXB−2066735
[Anthoceros angustus] AANG009306
[Anthoceros agrestis] AagrBONN_344.1477.1|AagrOXF_utg000006l.46.1
[Anthoceros punctatus] Apun_utg000092l.24.1
[Alsophila spinulosa] Aspi01Gene30454
[Lycopodium clavatum] EVM0011127.1
[Lycopodium clavatum] EVM0020843.1
[Lycopodium clavatum] EVM0024862.1
[Phylloglossum drummondii] ZZEI−2004900
[Phylloglossum drummondii] ZZEI−2022439
[Huperzia lucidula] GKAG−2097758
[Huperzia selago] GTJO−2027351
[Huperzia selago] NYBX−2015732
[Metacalypogeia alternifolia] Malternifolia_DN91210_c0_g1_i2.p1
[Fossombronia cristula] Fcristula_DN48901_c0_g1_i9.p1
[Marchantia polymorpha] Mp2g10420.1 (MpKAOL3)
[Marchantia paleacea] Marpal_utg000013g0034181
[Asterella wallichiana] Awallichiana_DN122708_c0_g1_i6.p1
[Phaeoceros carolinianus] HXRQ−2137136
[Phaeoceros carolinianus] ZFRE−2113863
[Anthoceros angustus] AANG008406
[Anthoceros agrestis] AagrBONN_344.2250.1|AagrOXF_utg000006l.858.1
[Phaeoceros carolinianus] RXRQ−2138606
[Anthoceros punctatus] Apun_utg000185l.423.1
[Pallavicinia lyellii] YFGP−2106822
[Pellia endiviifolia] Pendiviifolia_DN15889_c0_g1_i1.p1
[Blasia pusilla] Bpusilla_DN170761_c0_g1_i1.p1
[Lunularia cruciata] Lc_composite_02225.1_fr0
[Lunularia cruciata] Lc_composite_25679.1_fr0
[Lunularia cruciata] Lc_composite_21006.1_fr0
[Marchantia polymorpha] Mp4g23680.1_CYP88K1 (MpKAOL1)
[Marchantia paleacea] Marpal_utg000006g0014311
[Riccia cavernosa] Rcavernosa_DN108574_c0_g1_i2.p1
[Conocephalum conicum] ILBQ−2043688
[Asterella wallichiana] Awallichiana_DN125113_c0_g1_i4.p1
[Monosolenium tenerum] Mtenerum_DN28987_c0_g1_i1.p1
[Haplomitrium mnioides] Hmnioides_DN130928_c0_g1_i8.p1
[Treubia lacunosa] Tlacunosa_DN41748_c0_g1_i1.p1
[Calypogeia fissa] RTMU−2183258
[Makinoa crispata] Mcrispata_DN12474_c0_g1_i2.p1
[Paraphymatoceros hallii] FAJB−2009761
[Anthoceros agrestis] AagrBONN_228.360.1|AagrOXF_utg000009l.289.1
[Anthoceros angustus] AANG007216
[Anthoceros punctatus] Apun_utg000038l.711.1
[Anthoceros agrestis] AagrBONN_228.514.1
[Anthoceros agrestis] AagrOXF_utg000009l.123.1
[Isoetes tegetiformans] PKOX−2098322
[Isoetes sp.] PYHZ−2073760
[Selaginella moellendorffii] Sm_86321
[Selaginella selaginoides] KUXM−2002048
[Selaginella stauntoniana] ZZOL−2004176|ZZOL−2004177
[Selaginella apoda] LGDQ−2005560
[Selaginella lepidophylla] ABIJ−2038562
[Selaginella stauntoniana] ZZOL−2002991
[Selaginella moellendorffii] Sm_103517_CYP88E1
[Selaginella kraussiana] ZFGK−2004936
[Selaginella acanthonota] ZYCD−2009085
[Selaginella wallacei] JKAA−2001911
[Salvinia cucullata] Sacu_v1.1_s0185.g025018_CYP88V1
[Azolla filiculoides] Azfi_s0114.g046089
[Marsilea vestita] Mvestita_S4g08674−RA
[Marsilea vestita] Mvestita_S4g08989−RA
[Marsilea vestita] Mvestita_S4g08675−RA
[Ceratopteris richardii] Ceric.01G052700.1
[Adiantum capillus−veneris] Adc10773
[Alsophila spinulosa] Aspi01Gene10158
[Gnetum montanum] TnS000032623t02
[Ginkgo biloba] Gbi_chr9.1105_CYP88A12
[Ginkgo biloba] Gbi_chr9.1273_CYP88A109
[Picea abies] MA_107159g0010
[Taxus chinensis] KAH9302348.1
[Sequoiadendron giganteum] SEGI_32301
[Torreya grandis] TG1g02421−mRNA1
[Nymphaea colorata] Nycol.L01137.1
[Amborella trichopoda] AmTrH1.13G003200.1|AmTrH2.13G003100.1_CYP88A
[Cinnamomum kanehirae] CKAN_01881200
[Cinnamomum kanehirae] CKAN_00924000
[Liriodendron chinensis] Lchi01358
[Liriodendron chinensis] Lchi05940
[Oryza sativa] LOC_Os06g02019.1_CYP88A5 (OsKAO)
[Brachypodium distachyon] Bradi5g00467.1_CYP88A33
[Brachypodium distachyon] Bradi4g05240.2_CYP88A32P
[Brachypodium distachyon] Bradi1g30807.1_CYP88A34
[Brachypodium distachyon] Bradi1g51780.1_CYP88A31
[Populus trichocarpa] Potri.014G179100.2_CYP88A9
[Populus trichocarpa] Potri.013G161800.1_CYP88A8
[Arabidopsis thaliana] AT2G32440.1_CYP88A4 (AtKAO2)
[Arabidopsis thaliana] AT1G05160.1_CYP88A3 (AtKAO1)
CYP88
(KAO)
Clade
Liverwort
Hornwort
Lycophyte
Fern
Gymnosperm
Angiosperm
0.4

**Supplemental Figure 9** Phylogenetic tree of KAO and closely-related P450 enzymes in land plants. Nodes were labelled with percentage support values from 1000 standard non-parametric bootstraps by IQ-TREE 2. *M. polymorpha* genes of interest were indicated with red arrows. Related to Figure 4.

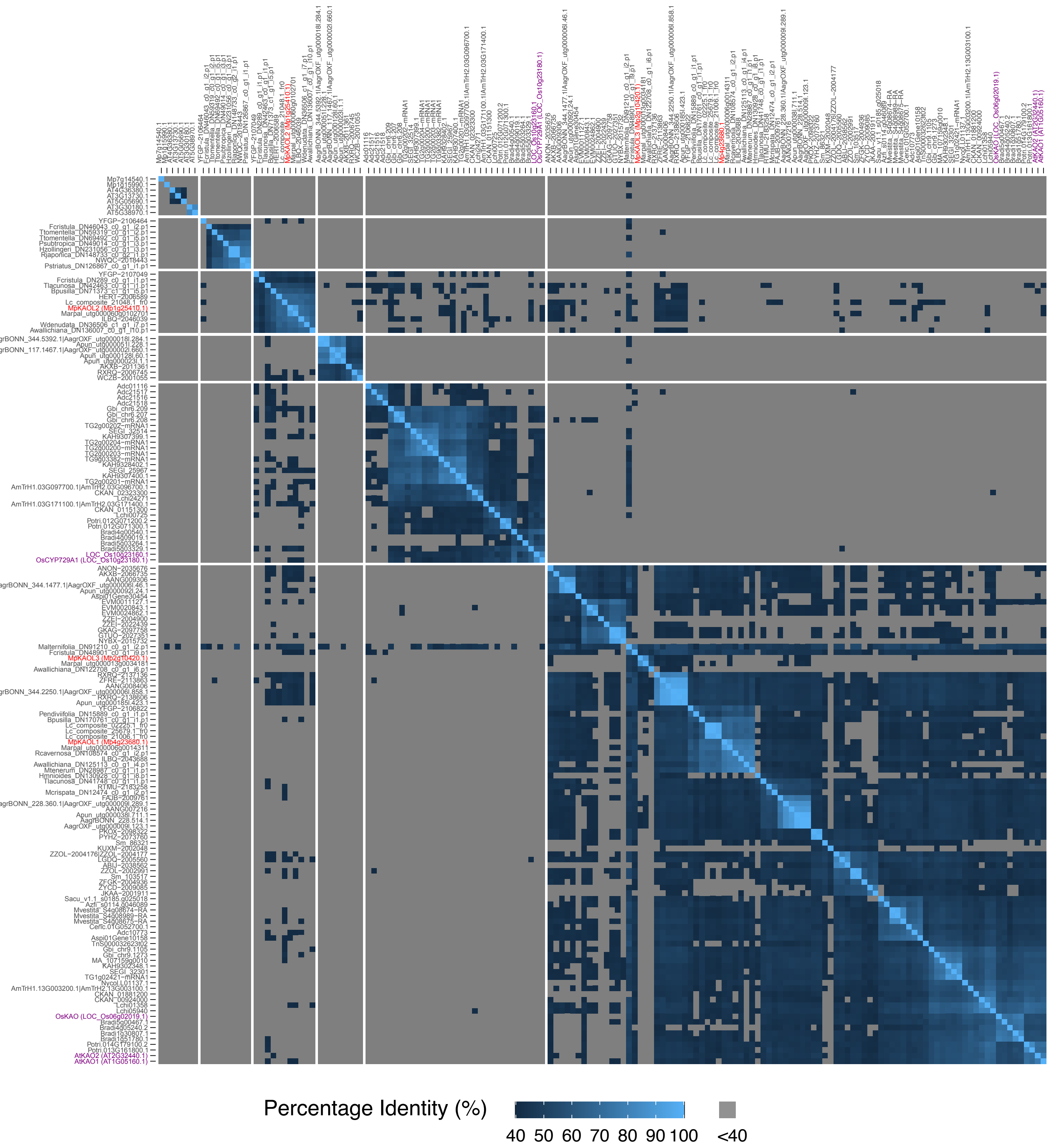


**Supplemental Figure 10** Heatmap of percentage identity for KAO and closely-related CYP enzymes in land plants. Sequences shown in Supplemental Figure 9 were included in the analysis, and percentage identities were calculated by local alignment using BLAST. An arbitrary threshold of 40% was set for the visualization of protein identities, as it is the threshold for defining membership in a CYP family (Nelson, 2006). MpKAOL2 and its liverwort homologs showed limited similarity to CYP729 or *bona fide* KAO (CYP88) family members. *M. polymorpha* genes of interest were colored in red, and their closely-related genes from *A. thaliana* or *O. sativa* were colored in purple. Related to Figure 4.

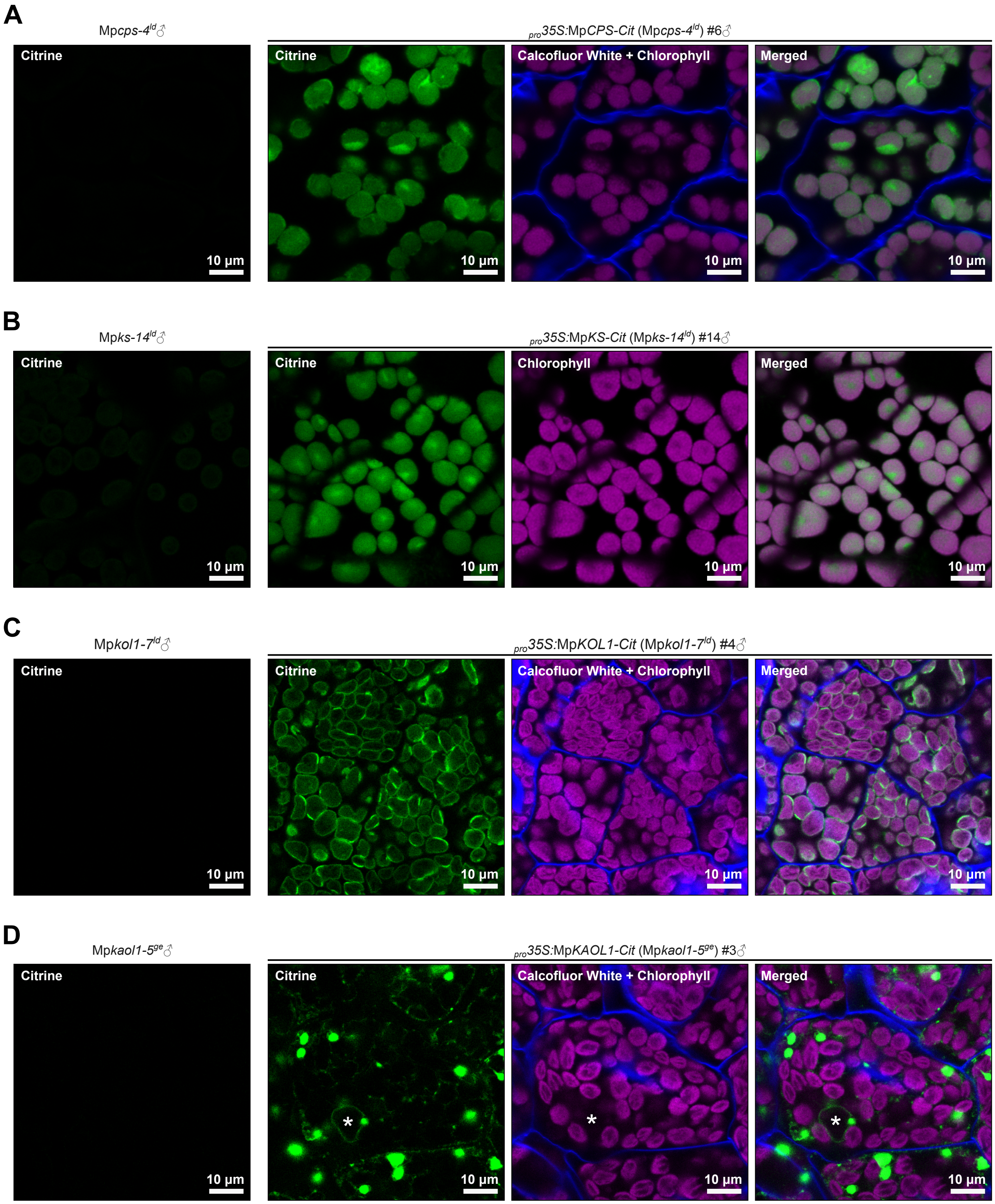


**Supplemental Figure 11** Subcellular localization of GA biosynthesis enzymes in *M. polymorpha*. A, C, and D, Images of 7-day-old plants cultured under cW+cFR, fixed and stained with calcofluor white to visualize the cell wall (shown in blue). B, Images of fresh gemmae. Citrine signals were shown in green, and the autofluorescence of chlorophyll was shown in magenta. Asterisks (*) in D indicate the position of a cell nucleus. Related to Figure 4.

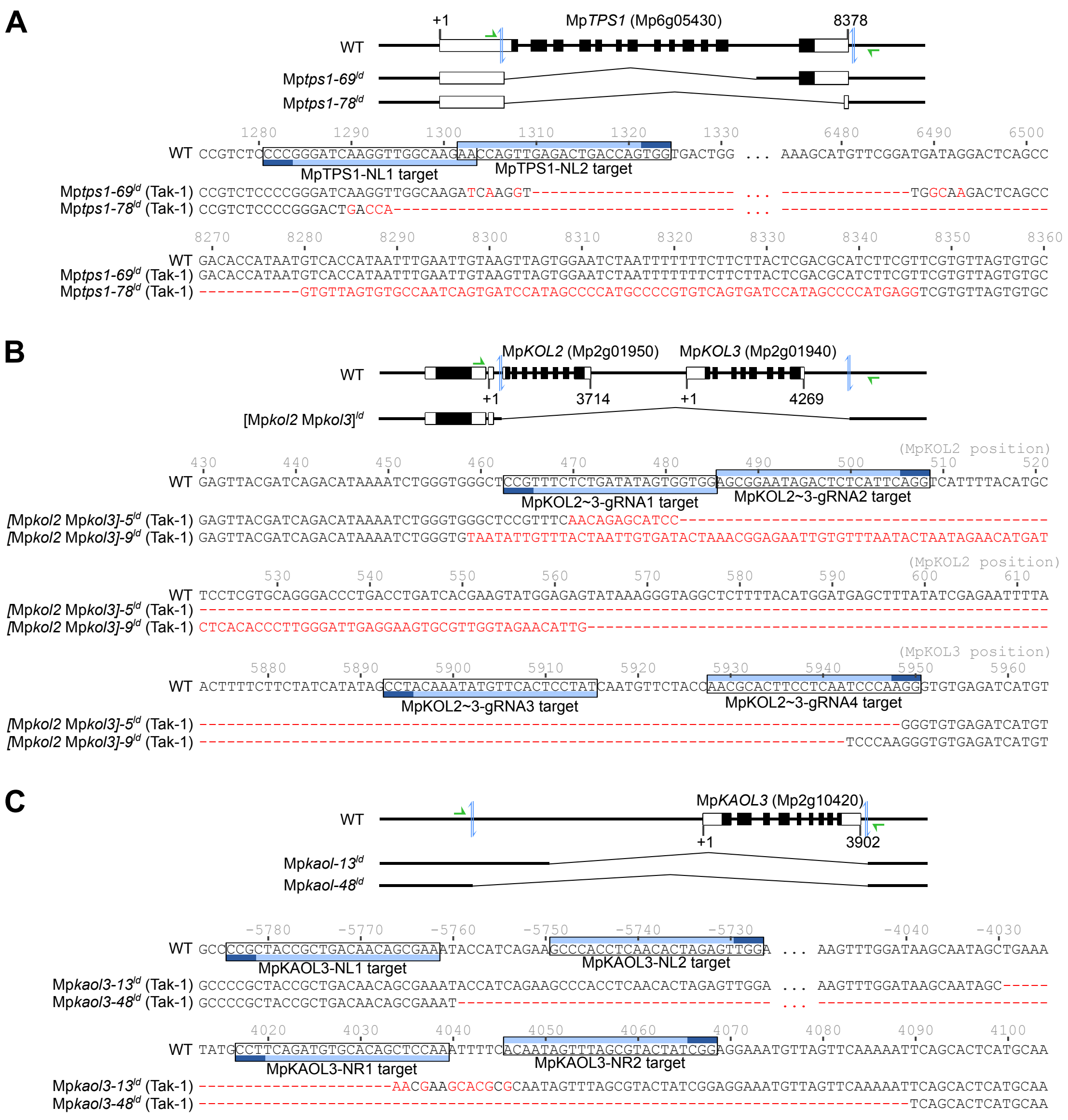


**Supplemental Figure 12** Genotype information for Mp*tps1*$^{ld}$, [Mp*kol2* Mp*kol3*]$^{ld}$ and Mp*kaol3*$^{ld}$ mutants. WT refers to reference sequences from MpTak_v6.1 genome assembly. In the schematic presentations of genomic structures, white and black rectangles represent untranslated and coding regions of exons, respectively. Targets of guide RNAs are indicated by blue arrows in the scheme and frames in the sequence (dark blue: the protospacer adjacent motif). Green arrows indicate the binding sites of genotyping primers. Indels and substitutions are shown in red letters. Numbers above the sequences indicate positions relative to the transcription start sites (+1). Related to Figure 4.

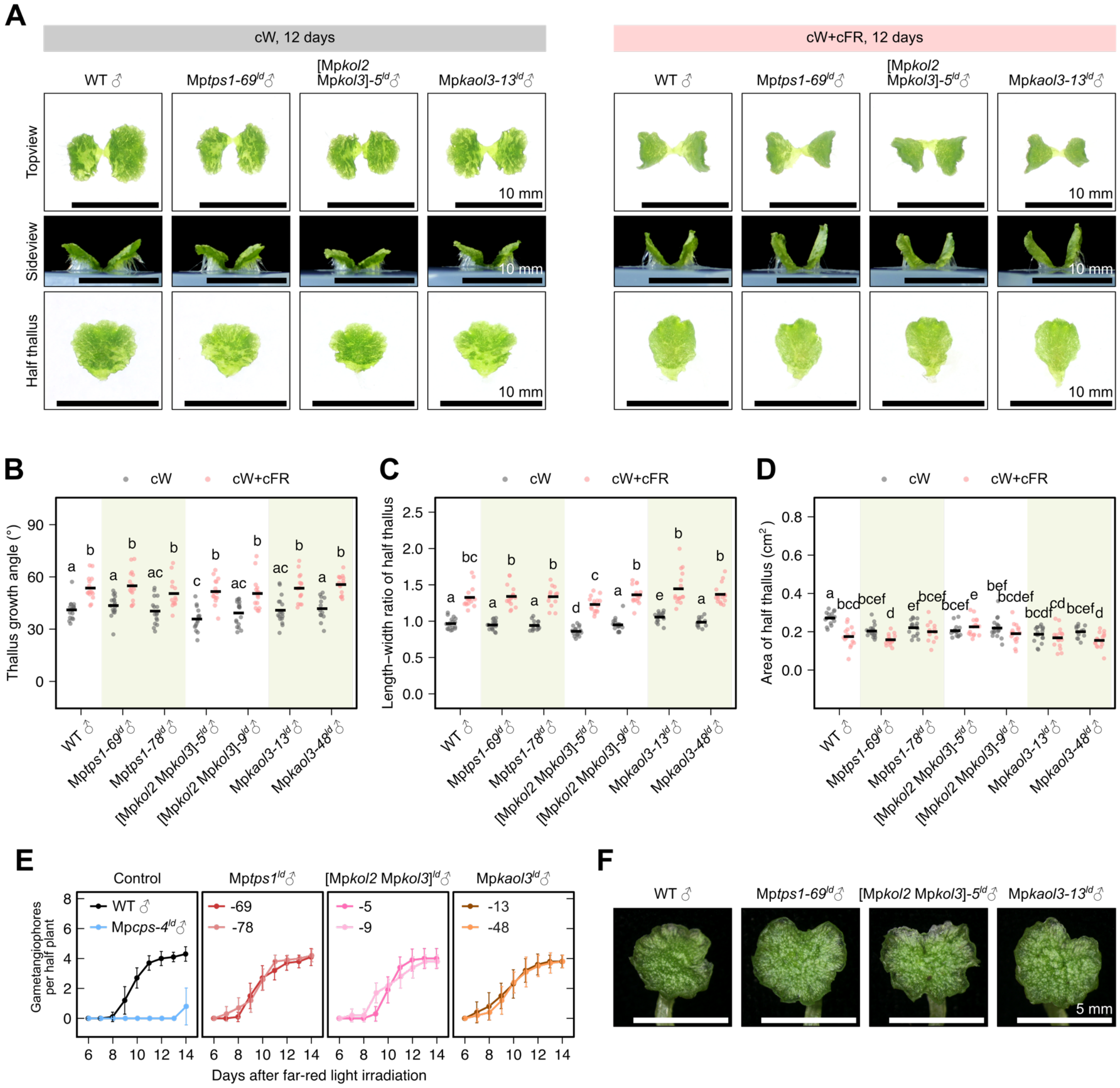

**Supplemental Figure 13** Phenotypes of Mp*tps1*$^{ld}$, [Mp*kol2* Mp*kol3*]$^{ld}$ and Mp*kaol3*$^{ld}$ mutants. A-D, Morphology (A) and measurements (B-D) of 12-day-old thalli grown from gemmae under cW or cW+cFR. Horizontal lines represent mean values, and letters represent multiple comparisons with non-pooled Welch's *t*-test and B-H adjustment (adjusted p<0.05 for non-overlapping letters, n=11-18). E-F, Gametangiophore formation progress (E) and morphology (F) of mutants, grown under cW for 7 days before transferred to cW+cFR in aseptic culture. Dots and error bars represent mean±SD in H (n=5). Bars = 5 mm in F. Related to Figure 4.

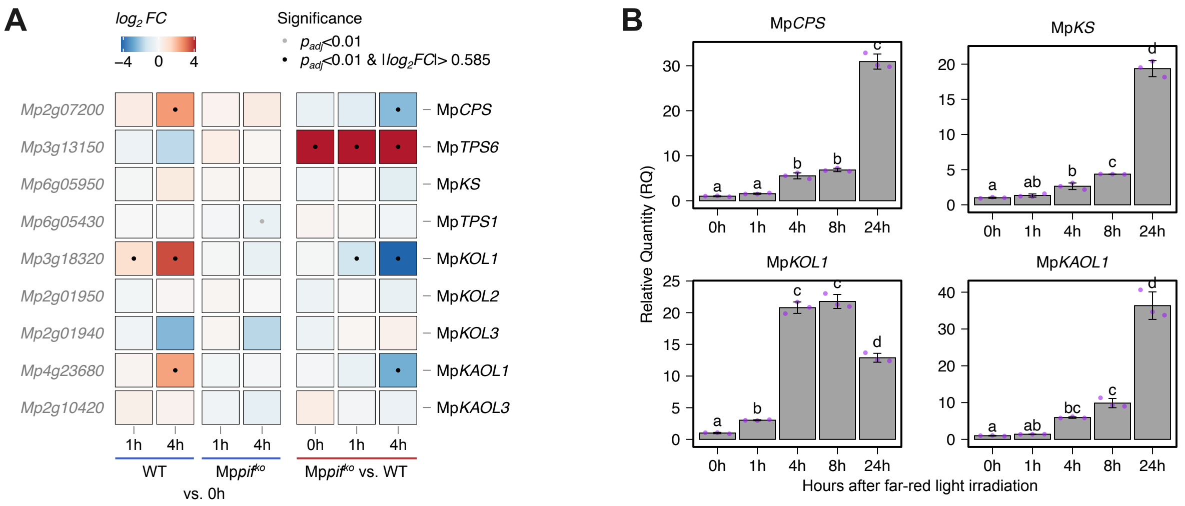


**Supplemental Figure 14** Up-regulation of GA biosynthesis genes by FR irradiation. A, Heatmap of gene expression changes after FR irradiation, using data from (Hernández-García et al., 2021). The transcriptomes were sequenced from plants cultured under continuous red light for 7 days, then irradiated with FR light for 0, 1 or 4 hours. FC, fold change. $p_{adj}$, adjusted $p$-value. B, Relative expression level of GA biosynthesis genes in Tak-1 wild-type plants, quantified by qPCR from plants cultured under cW for 7 days and then transferred to cW+cFR for indicated hours. The plots represent mean±SD from 3 biological replicates. Related to Figure 5.

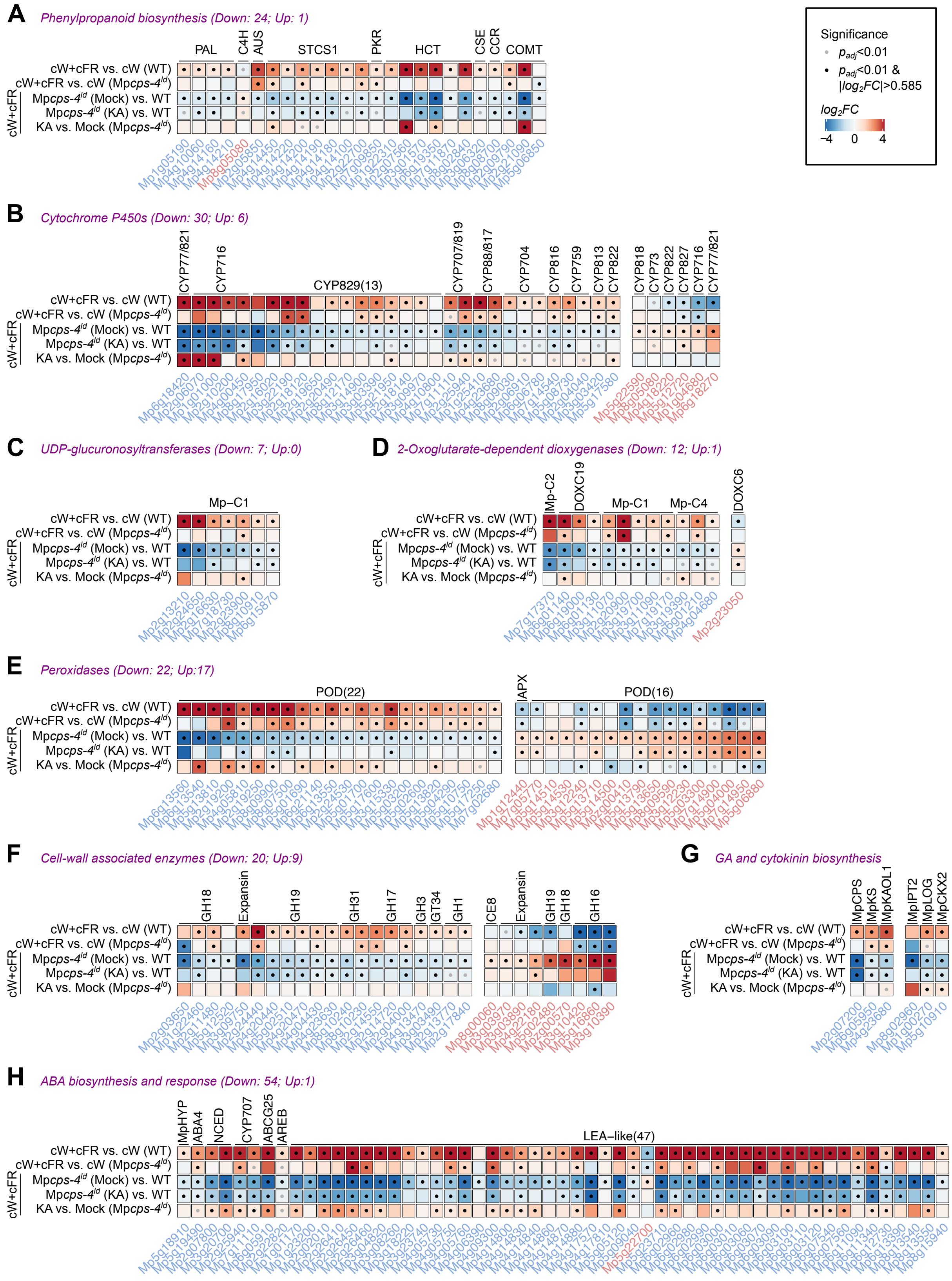


**Supplemental Figure 15** Heatmaps of genes from selected pathways or gene families, showing differential expression in Mp*cps-4*$^{ld}$ under cW+cFR. FC, fold change. $p_{adj}$, adjusted *p*-value. Related to Figure 5.

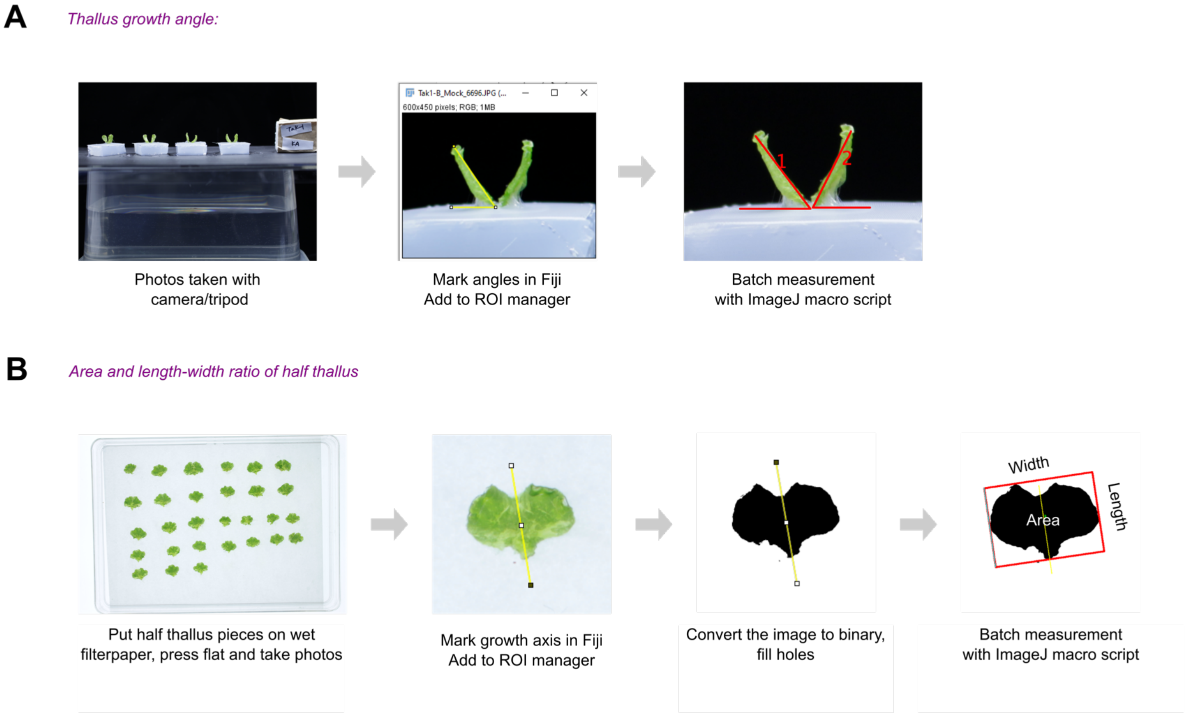


**Supplemental Figure 16** Quantification method for thallus morphology. A, For the measurement of growth angles, blocks of agar medium with plants were cut out and aligned at certain distance to a fixed camera. After photos were taken from the side view, angles between the thallus and the medium surface were measured for each half thallus with Fiji/ImageJ. B, For the measurement of thallus area and length-width ratio, photos were taken from the top for half thalli flattened on a filter paper. A growth axis was defined manually for each half-thallus, pointing from the basal end to the first bifurcation point. A minimum bounding rectangle was created around the thallus along the direction of the growth axis, and the edges of this rectangle defined the length and width of the thallus. Related to methods.

**Supplemental Table 1** List of plant materials

| Name | Genetic background | Sex | Vector for construction | Source |
|---|---|---|---|---|
| Tak-1 | - | Male | - | (Ishizaki et al., 2016) |
| Tak-2 | - | Female | - | (Ishizaki et al., 2016) |
| Mp*cps-4$^{ld}$* | Tak-1 | Male | pMpGE017-MpCPS-LD | This paper |
| Mp*cps-27$^{ld}$* | Tak-1 | Male | pMpGE017-MpCPS-LD | This paper |
| *$_{pro}$35S:*Mp*CPS-Cit* (Mp*cps-4$^{ld}$*) #6 | Mp*cps-4$^{ld}$* | Male | pMpGWB306-MpCPS-CDS | This paper |
| *$_{pro}$35S:*Mp*CPS-Cit* (Mp*cps-4$^{ld}$*) #7 | Mp*cps-4$^{ld}$* | Male | pMpGWB306-MpCPS-CDS | This paper |
| Mp*cps-120$^{ld}$* | Tak-2 | Female | pMpGE017-MpCPS-LD | This paper |
| Mp*cps-123$^{ld}$* | Tak-2 | Female | pMpGE017-MpCPS-LD | This paper |
| *$_{pro}$35S:*Mp*CPS-Cit* (Mp*cps-120$^{ld}$*) #2 | Mp*cps-120$^{ld}$* | Female | pMpGWB306-MpCPS-CDS | This paper |
| *$_{pro}$35S:*Mp*CPS-Cit* (Mp*cps-120$^{ld}$*) #6 | Mp*cps-120$^{ld}$* | Female | pMpGWB306-MpCPS-CDS | This paper |
| Mp*cps-5$^{ld}$* (Mp*BNB-Cit* ♂) | Mp*BNB-Cit* ♂ | Male | pMpGE018-MpCPS-LD | (Yamaoka et al., 2018) |
| Mp*cps-6$^{ld}$* (Mp*BNB-Cit* ♂) | Mp*BNB-Cit* ♂ | Male | pMpGE018-MpCPS-LD | (Yamaoka et al., 2018) |
| Mp*cps-2$^{ld}$* (Mp*BNB-Cit* ♀) | Mp*BNB-Cit* ♀ | Female | pMpGE018-MpCPS-LD | (Yamaoka et al., 2018) |
| Mp*cps-3$^{ld}$* (Mp*BNB-Cit* ♀) | Mp*BNB-Cit* ♀ | Female | pMpGE018-MpCPS-LD | (Yamaoka et al., 2018) |
| Mp*ks-14$^{ld}$* | Tak-1 | Male | pMpGE018-MpKS-LD | This paper |
| Mp*ks-19$^{ld}$* | Tak-1 | Male | pMpGE018-MpKS-LD | This paper |
| *$_{pro}$35S:*Mp*KS-Cit* (Mp*ks-14$^{ld}$*) #13 | Mp*ks-14$^{ld}$* | Male | pMpGWB106-MpKS-CDS | This paper |
| *$_{pro}$35S:*Mp*KS-Cit* (Mp*ks-14$^{ld}$*) #14 | Mp*ks-14$^{ld}$* | Male | pMpGWB106-MpKS-CDS | This paper |
| Mp*kol1-7$^{ld}$* | Tak-1 | Male | pMpGE018-MpKOL1-LD | This paper |
| Mp*kol1-13$^{ld}$* | Tak-1 | Male | pMpGE018-MpKOL1-LD | This paper |
| *$_{pro}$35S:*Mp*KOL1-Cit* (Mp*kol1-7$^{ld}$*) #2 | Mp*kol1-7$^{ld}$* | Male | pMpGWB106-MpKOL1-CDS | This paper |
| *$_{pro}$35S:*Mp*KOL1-Cit* (Mp*kol1-7$^{ld}$*) #4 | Mp*kol1-7$^{ld}$* | Male | pMpGWB106-MpKOL1-CDS | This paper |
| Mp*kaol1-5$^{ge}$* | Tak-1 | Male | pMpGE011-MpKAOL1-gRNA1 | This paper |
| Mp*kaol1-7$^{ge}$* | Tak-1 | Male | pMpGE011-MpKAOL1-gRNA1 | This paper |
| *$_{pro}$35S:*Mp*KAOL1$^{mut}$-Cit* (Mp*kaol1-5$^{ge}$*) #3 | Mp*kaol1-5$^{ge}$* | Male | pMpGWB106-MpKAOL1-CDSmut | This paper |
| *$_{pro}$35S:*Mp*KAOL1$^{mut}$-Cit* (Mp*kaol1-5$^{ge}$*) #4 | Mp*kaol1-5$^{ge}$* | Male | pMpGWB106-MpKAOL1-CDSmut | This paper |
| Mp*tps1-52$^{ld}$* | Tak-1 | Male | pMpGE018-MpTPS1-LD | This paper |
| Mp*tps1-69$^{ld}$* | Tak-1 | Male | pMpGE018-MpTPS1-LD | This paper |
| *[*Mp*kol2* Mp*kol3]-5$^{ld}$* | Tak-1 | Male | pMpGE018-MpKOL2/3-LD | This paper |
| *[*Mp*kol2* Mp*kol3]-9$^{ld}$* | Tak-1 | Male | pMpGE018-MpKOL2/3-LD | This paper |
| Mp*kaol3-13$^{ld}$* | Tak-1 | Male | pMpGE018-MpKAOL3-LD | This paper |
| Mp*kaol3-48$^{ld}$* | Tak-1 | Male | pMpGE018-MpKAOL3-LD | This paper |
| Mp*pif$^{ko}$* #1 | Tak-1×Tak-2 F1 | Female | - | (Inoue et al., 2016) |
| *g*Mp*PIF/*Mp*pif$^{ko}$* #1 | Mp*pif$^{ko}$* #1 | Female | - | (Inoue et al., 2016) |

**Supplemental Table 2** List of plasmids

| Name | Source | Identifier |
|---|---|---|
| pMpGE_En04 | (Hisanaga et al., 2019) | - |
| pBCGE12 | (Hisanaga et al., 2019) | - |
| pBCGE23 | (Hisanaga et al., 2019) | - |
| pBCGE34 | (Hisanaga et al., 2019) | - |
| pMpGE017 | (Hisanaga et al., 2019) | - |
| pMpGE018 | (Hisanaga et al., 2019) | - |
| pMpGE_En04-MpCPS-NL1 | This paper | - |
| pBCGE12-MpCPS-NL2 | This paper | - |
| pBCGE23-MpCPS-NR1 | This paper | - |
| pBCGE34-MpCPS-NR2 | This paper | - |
| pMpGE_En04-MpCPS-LD | This paper | - |
| pMpGE017-MpCPS-LD | This paper | - |
| pMpGE018-MpCPS-LD | This paper | - |
| pENTR/D-TOPO | Thermo Fisher | Cat#K240020 |
| pENTR-MpCPS-CDS | This paper | - |
| pMpGWB306 | (Ishizaki et al., 2015) | Addgene #68637 |
| pMpGWB306-MpCPS-CDS | This paper | - |
| pMpGE_En04-MpKS-NL1 | This paper | - |
| pBCGE12-MpKS-NL2 | This paper | - |
| pBCGE23-MpKS-NR1 | This paper | - |
| pBCGE34-MpKS-NR2 | This paper | - |
| pMpGE_En04-MpKS-LD | This paper | - |
| pMpGE018-MpKS-LD | This paper | - |
| pENTR-MpKS-CDS | This paper | - |
| pMpGWB106 | (Ishizaki et al., 2015) | Addgene #68560 |
| pMpGWB106-MpKS-CDS | This paper | - |
| pMpGE_En04-MpKOL1-gRNA1 | This paper | - |
| pBCGE12-MpKOL1-gRNA2 | This paper | - |
| pBCGE23-MpKOL1-gRNA3 | This paper | - |
| pBCGE34-MpKOL1-gRNA4 | This paper | - |
| pMpGE_En04-MpKOL1-gRNA1~4 | This paper | - |
| pMpGE018-MpKOL1-gRNA1~4 | This paper | - |
| pENTR-MpKOL1-CDS-NoStop | This paper | - |
| pMpGWB106-MpKOL1-CDS | This paper | - |
| pMpGE_En03 | (Sugano et al., 2018) | Addgene #71535 |
| pMpGE011 | (Sugano et al., 2018) | Addgene #71537 |
| pMpGE_En03-MpKAOL1-gRNA1 | This paper | - |
| pMpGE011-MpKAOL1-gRNA1 | This paper | - |
| pENTR-MpKAOL1-CDS-NoStop | This paper | - |
| pENTR-MpKAOL1-CDSmut-NoStop | This paper | - |
| pMpGWB106-MpKAOL1-CDSmut | This paper | - |
| pMpGE_En04-MpTPS1-NL1 | This paper | - |
| pBCGE12-MpTPS1-NL2 | This paper | - |
| pBCGE23-MpTPS1-NR1 | This paper | - |
| pBCGE34-MpTPS1-NR2 | This paper | - |
| pMpGE_En04-MpTPS1-LD | This paper | - |
| pMpGE018-MpTPS1-LD | This paper | - |
| pMpGE_En04-MpKOL2/3-gRNA1 | This paper | - |
| pBCGE12-MpKOL2/3-gRNA2 | This paper | - |
| pBCGE23-MpKOL2/3-gRNA3 | This paper | - |
| pBCGE34-MpKOL2/3-gRNA4 | This paper | - |
| pMpGE_En04-MpKOL2/3-LD | This paper | - |
| pMpGE018-MpKOL2/3-LD | This paper | - |
| pMpGE_En04-MpKAOL3-NL1 | This paper | - |
| pBCGE12-MpKAOL3-NL2 | This paper | - |
| pBCGE23-MpKAOL3-NR1 | This paper | - |
| pBCGE34-MpKAOL3-NR2 | This paper | - |
| pMpGE_En04-MpKAOL3-LD | This paper | - |

| | | |
|---|---|---|
| pMpGE017-MpKAOL3-LD | This paper | - |
| pPICZA | Thermo Fisher | Cat #V19020 |
| pPICZA-AtKO | This paper | - |
| pPICZA-MpKOL1 | This paper | - |
| pPICZA-MpKOL2 | This paper | - |
| pPICZA-MpKOL3 | This paper | - |
| pPICZA-AtKAO1 | This paper | - |
| pPICZA-MpKAOL1 | This paper | - |
| pPICZA-MpKAOL3 | This paper | - |

### **Supplemental Table 3** List of DNA oligos

| Name | Sequence (5′→3′) | Used for |
|---|---|---|
| MpCPS-NL1-OligoA | CTCGATCAACCTTACGAACCGGAC | pMpGE_En04-MpCPS-NL1 |
| MpCPS-NL1-OligoB | AAACGTCCGGTTCGTAAGGTTGAT | pMpGE_En04-MpCPS-NL1 |
| MpCPS-NL2-OligoA | CTCGGATAACTGCCACAGCGAAGC | pBCGE12-MpCPS-NL2 |
| MpCPS-NL2-OligoB | AAACGCTTCGCTGTGGCAGTTATC | pBCGE12-MpCPS-NL2 |
| MpCPS-NR1-OligoA | CTCGTTCGGGTACAAGGGTTTGGA | pBCGE23-MpCPS-NR1 |
| MpCPS-NR1-OligoB | AAACTCCAAACCCTTGTACCCGAA | pBCGE23-MpCPS-NR1 |
| MpCPS-NR2-OligoA | CTCGGAATGTCTAGTACGGAGCTT | pBCGE34-MpCPS-NR2 |
| MpCPS-NR2-OligoB | AAACAAGCTCCGTACTAGACATTC | pBCGE34-MpCPS-NR2 |
| MpCPS-gt-F | GGAACCTATCCGGGGATCCT | Genotyping of Mp*cps*$^{ld}$ |
| MpCPS-gt-R | ATGTGACGTTCGTTTGCTGC | Genotyping of Mp*cps*$^{ld}$ |
| CACC-MpCPS-CDS-F | CACCATGGCATTCTCGTTAGCAGGT | pENTR-MpCPS-CDS |
| MpCPS-CDS-R | GGCCACAGGCTCGAAGAGTA | pENTR-MpCPS-CDS |
| MpKS-NL1-OligoA | CTCGTGTGGAACATAGAGTCTTGC | pMpGE_En04-MpKS-NL1 |
| MpKS-NL1-OligoB | AAACGCAAGACTCTATGTTCCACA | pMpGE_En04-MpKS-NL1 |
| MpKS-NL2-OligoA | CTCGTCCACAGAGTCTTGTTCGTC | pBCGE12-MpKS-NL2 |
| MpKS-NL2-OligoB | AAACGACGAACAAGACTCTGTGGA | pBCGE12-MpKS-NL2 |
| MpKS-NR1-OligoA | CTCGTGCTTGCTGTCCTGATGTCC | pBCGE23-MpKS-NR1 |
| MpKS-NR1-OligoB | AAACGGACATCAGGACAGCAAGCA | pBCGE23-MpKS-NR1 |
| MpKS-NR2-OligoA | CTCGCAAGCATACGTCCGCCACTA | pBCGE34-MpKS-NR2 |
| MpKS-NR2-OligoB | AAACTAGTGGCGGACGTATGCTTG | pBCGE34-MpKS-NR2 |
| MpKS-gt-F | ACTGTGAGCTGAAACTGCAGA | Genotyping of Mp*ks*$^{ld}$ |
| MpKS-gt-R | GGACGGACATGGATCTAGCA | Genotyping of Mp*ks*$^{ld}$ |
| CACC-MpTPS4-CDS-F | CACCATGATGATCCATCCAGCTATTGTG | pENTR-MpKS-CDS |
| MpTPS4-CDS-R | GGCCTGTTCACTTTCGATGG | pENTR-MpKS-CDS |
| Mapoly0140s0010-gRNA1-F | CTCGGATCATGGCTTTTCTCCCGC | pMpGE_En04-MpKOL1-gRNA1 |
| Mapoly0140s0010-gRNA1-R | AAACGCGGGAGAAAAGCCATGATC | pMpGE_En04-MpKOL1-gRNA1 |
| Mapoly0140s0010-gRNA2-F | CTCGGATGCTCGCTCCATAAAAAC | pBCGE12-MpKOL1-gRNA2 |
| Mapoly0140s0010-gRNA2-R | AAACGTTTTTATGGAGCGAGCATC | pBCGE12-MpKOL1-gRNA2 |
| Mapoly0140s0010-gRNA3-F | CTCGATCCGACAAATAATGTTTGT | pBCGE23-MpKOL1-gRNA3 |
| Mapoly0140s0010-gRNA3-R | AAACACAAACATTATTTGTCGGAT | pBCGE23-MpKOL1-gRNA3 |
| Mapoly0140s0010-gRNA4-F | CTCGGTTCAGTTTAGAAACCCTCC | pBCGE34-MpKOL1-gRNA4 |
| Mapoly0140s0010-gRNA4-R | AAACGGAGGGTTTCTAAACTGAAC | pBCGE34-MpKOL1-gRNA4 |
| Dseq-KOL1-gRNA1~4F | GGATTGATGTACTTGACGAG | Genotyping of Mp*kol1*$^{ld}$ |
| Dseq-KOL1-gRNA1~4R | TTCGGCCTGAAGTCTAAGAG | Genotyping of Mp*kol1*$^{ld}$ |
| CACC-MpKOL1-CDS-F | CACCATGAAATGCTTCGGTTTG | pENTR-MpKOL1-CDS |
| MpKOL1-CDS-ns-R | AATCTTCGCTGGACAG | pENTR-MpKOL1-CDS |
| MpKAO-gRNA1F | CTCGCCAGGCTCCTCTCCCCCC | pMpGE_En03-MpKAOL1-gRNA1 |
| MpKAO-gRNA1R | AAACGGGGGGAGAGGAGCCTGG | pMpGE_En03-MpKAOL1-gRNA1 |
| MpKAO-Dseq1-F | GAGGCATTGAGATCGAGAGG | Genotyping of Mp*kaol1*$^{ge}$ |
| MpKAO-Dseq1-R | ATACTCTCGGCGGTCGTTGC | Genotyping of Mp*kaol1*$^{ge}$ |
| Mapoly0020s0131-F | CACCATGTTGGAGATTTCGTCCAC | pENTR-MpKAOL1-CDS-NoStop |
| MpKAOL1-CDS-ns-R | CAATCGTGAGAAGTTTATAAGACAG | pENTR-MpKAOL1-CDS-NoStop |
| MpKAOL1-mut-F | GCTGCCGCCGGGAGACATGGGCTGG | pENTR-MpKAOL1-CDSmut-NoStop |
| MpKAOL1-mut-R | GGTGCTTGCCCTTTCTGAAGACTGGG | pENTR-MpKAOL1-CDSmut-NoStop |
| MpTPS1-NL1-OligoA | CTCGGTTCTTGCCAACCTTGATCC | pMpGE_En04-MpTPS1-NL1 |
| MpTPS1-NL1-OligoB | AAACGGATCAAGGTTGGCAAGAAC | pMpGE_En04-MpTPS1-NL1 |
| MpTPS1-NL2-OligoA | CTCGAACCAGTTGAGACTGACCAG | pBCGE12-MpTPS1-NL2 |
| MpTPS1-NL2-OligoB | AAACCTGGTCAGTCTCAACTGGTT | pBCGE12-MpTPS1-NL2 |
| MpTPS1-NR1-OligoA | CTCGACGTGACTGTGTTGAGTCTA | pBCGE23-MpTPS1-NR1 |
| MpTPS1-NR1-OligoB | AAACTAGACTCAACACAGTCACGT | pBCGE23-MpTPS1-NR1 |
| MpTPS1-NR2-OligoA | CTCGCAGTCACGTCACTACGAGAC | pBCGE34-MpTPS1-NR2 |
| MpTPS1-NR2-OligoB | AAACGTCTCGTAGTGACGTGACTG | pBCGE34-MpTPS1-NR2 |
| MpTPS1-gt-F | GGCCTCTCGTAGCTTTGA | Genotyping of Mp*tps1*$^{ld}$ |
| MpTPS1-gt-R | CAGGAAGGTTTGCTTGCA | Genotyping of Mp*tps1*$^{ld}$ |
| Mapoly0130s0002-0003-gRNA1-F | CTCGCCACCACTATATCAGAGAAA | pMpGE_En04-MpKOL2/3-gRNA1 |
| Mapoly0130s0002-0003-gRNA1-R | AAACTTTCTCTGATATAGTGGTGG | pMpGE_En04-MpKOL2/3-gRNA1 |
| Mapoly0130s0002-0003-gRNA2-F | CTCGAGCGGAATAGACTCTCATTC | pBCGE12-MpKOL2/3-gRNA2 |
| Mapoly0130s0002-0003-gRNA2-R | AAACGAATGAGAGTCTATTCCGCT | pBCGE12-MpKOL2/3-gRNA2 |

| | | |
|---|---|---|
| Mapoly0130s0002-0003-gRNA3-F | CTCGATAGGAGTGAACATATTTGT | pBCGE23-MpKOL2/3-gRNA3 |
| Mapoly0130s0002-0003-gRNA3-R | AAACACAAATATGTTCACTCCTAT | pBCGE23-MpKOL2/3-gRNA3 |
| Mapoly0130s0002-0003-gRNA4-F | CTCGAACGCACTTCCTCAATCCCA | pBCGE34-MpKOL2/3-gRNA4 |
| Mapoly0130s0002-0003-gRNA4-R | AAACTGGGATTGAGGAAGTGCGTT | pBCGE34-MpKOL2/3-gRNA4 |
| Dseq-KOL2-3_gRNA1~4F | CTCCAAGTGTTGTGTAGCTG | Genotyping of *[Mpkol2 Mpkol3]*$^{ld}$ |
| Dseq-KOL2-3_gRNA1~4R | CTCTTAGCAGATGTGACCAC | Genotyping of *[Mpkol2 Mpkol3]*$^{ld}$ |
| MpKAOL3-NL1-OligoA | CTCGTTCGCTGTTGTCAGCGGTAG | pMpGE_En04-MpKAOL3-NL1 |
| MpKAOL3-NL1-OligoB | AAACCTACCGCTGACAACAGCGAA | pMpGE_En04-MpKAOL3-NL1 |
| MpKAOL3-NL2-OligoA | CTCGGCCCACCTCAACACTAGAGT | pBCGE12-MpKAOL3-NL2 |
| MpKAOL3-NL2-OligoB | AAACACTCTAGTGTTGAGGTGGGC | pBCGE12-MpKAOL3-NL2 |
| MpKAOL3-NR1-OligoA | CTCGTTGGAGCTGTGCACATCTGA | pBCGE23-MpKAOL3-NR1 |
| MpKAOL3-NR1-OligoB | AAACTCAGATGTGCACAGCTCCAA | pBCGE23-MpKAOL3-NR1 |
| MpKAOL3-NR2-OligoA | CTCGACAATAGTTTAGCGTACTAT | pBCGE34-MpKAOL3-NR2 |
| MpKAOL3-NR2-OligoB | AAACATAGTACGCTAAACTATTGT | pBCGE34-MpKAOL3-NR2 |
| MpKAOL3-gt-F | GGCACACACGAGACTCCC | Genotyping of Mp*kaol3*$^{ld}$ |
| MpKAOL3-gt-R | TCGCGAGGAGTAGGCTTT | Genotyping of Mp*kaol3*$^{ld}$ |
| pPICZA-AtKO-IF-F | ATTCGAAACGAGGAA ATGGCCTTCTTCTCCATGA | pPICZA-AtKO |
| pPICZA-AtKO-IF-R | CCCAAGCTGGCGGCC AGAACGCCTTGGATTGAT | pPICZA-AtKO |
| pPICZA-MpKOL1-IF-F | ATTCGAAACGAGGAA ATGAAATGCTTCGGTTTGTC | pPICZA-MpKOL1 |
| pPICZA-MpKOL1-IF-R | CCCAAGCTGGCGGCC AATCTTCGCTGGACAG | pPICZA-MpKOL1 |
| pPICZA-MpKOL2-IF-F | ATTCGAAACGAGGAA ATGACCAGACACTTGGGTGA | pPICZA-MpKOL2 |
| pPICZA-MpKOL2-IF-R | CCCAAGCTGGCGGCC AGCTGGCAAAATATTTTTCA | pPICZA-MpKOL2 |
| pPICZA-MpKOL3-IF-F | ATTCGAAACGAGGAA ATGGAGAGTACAGAGAAATC | pPICZA-MpKOL3 |
| pPICZA-MpKOL3-IF-R | CCCAAGCTGGCGGCC AGATGGCAGAACACCTTTGA | pPICZA-MpKOL3 |
| pPICZA-AtKAO1-IF-F | ATTCGAAACGAGGAA ATGGCGGAGACAACGAGTTG | pPICZA-AtKAO1 |
| pPICZA-AtKAO1-IF-R | CCCAAGCTGGCGGCC CTGATAACTAATTCTTGCCA | pPICZA-AtKAO1 |
| pPICZA-MpKAOL1-IF-F | ATTCGAAACGAGGAA ATGTTGGAGATTTCGTCCAC | pPICZA-MpKAOL1 |
| pPICZA-MpKAOL1-IF-R | CCCAAGCTGGCGGCC CAATCGTGAGAAGTTTATAA | pPICZA-MpKAOL1 |
| pPICZA-MpKAOL3-IF-F | ATTCGAAACGAGGAA ATGGCTGCGATTGTTCTCA | pPICZA-MpKAOL3 |
| pPICZA-MpKAOL3-IF-R | CCCAAGCTGGCGGCC GCTGCACACACGACGAGT | pPICZA-MpKAOL3 |
| MpEF1−qPCR_F | AAGCCGTCGAAAAGAAGGAG | qPCR for MpEF1 (reference) |
| MpEF1−qPCR_R | TTCAGGATCGTCCGTTATCC | qPCR for MpEF1 (reference) |
| MpCPSKS_qPCR-F | TCTTACACGGTTCTCGGGATG | qPCR for MpCPS |
| MpCPSKS_qPCR-R | GGATTGCGTTTTGAGGAAGATG | qPCR for MpCPS |
| MpKS-RT-F | CAAGCAAGGATAGCAATCCAG | qPCR for MpKS |
| MpKS-RT-R | TCGCATCATTCCCAACCAG | qPCR for MpKS |
| MpKO_qPCR-F1 | TGCAGCACTTCGAGTTGACC | qPCR for MpKOL1 |
| MpKO_qPCR-R1 | TGCAGTTTGTGGGAGGTGAC | qPCR for MpKOL1 |
| MpKAO_qPCR-F1 | GCCCTATGCGTTCAAACCTG | qPCR for MpKAOL1 |
| MpKAO_qPCR-R1 | GCTCGATGCCGATACAACTC | qPCR for MpKAOL1 |

Red letters indicate adapter sequences, and start codons are indicated with underline.

### Supplemental References


**Albone, K.S., Gaskin, P., MacMillan, J., Phinney, B.O., and Willis, C.L.** (1990). Biosynthetic Origin of Gibberellins $A_3$ and $A_7$ in Cell-Free Preparations from Seeds of *Marah macrocarpus* and *Malus domestica*. Plant Physiol. **94**: 132–142.

**Fujioka, S., Yamane, H., Spray, C.R., Phinney, B.O., Gaskin, P., MacMillan, J., and Takahashi, N.** (1990). Gibberellin $A_3$ Is Biosynthesized from Gibberellin $A_{20}$ via Gibberellin $A_5$ in Shoots of *Zea mays* L. Plant Physiol. **94**: 127–131.

**Hernández-García, J., Sun, R., Serrano-Mislata, A., Inoue, K., Vargas-Chávez, C., Esteve-Bruna, D., Arbona, V., Yamaoka, S., Nishihama, R., Kohchi, T., and Blázquez, M.A.** (2021). Coordination between growth and stress responses by DELLA in the liverwort *Marchantia polymorpha*. Current Biology.

**Hisanaga, T., Okahashi, K., Yamaoka, S., Kajiwara, T., Nishihama, R., Shimamura, M., Yamato, K.T., Bowman, J.L., Kohchi, T., and Nakajima, K.** (2019). A *cis*-acting bidirectional transcription switch controls sexual dimorphism in the liverwort. The EMBO Journal **38**: 1–12.

**Inoue, K., Nishihama, R., Kataoka, H., Hosaka, M., Manabe, R., Nomoto, M., Tada, Y., Ishizaki, K., and Kohchi, T.** (2016). Phytochrome signaling is mediated by PHYTOCHROME INTERACTING FACTOR in the liverwort *Marchantia polymorpha*. The Plant Cell **28**: 1406–1421.

**Ishizaki, K., Nishihama, R., Ueda, M., Inoue, K., Ishida, S., Nishimura, Y., Shikanai, T., and Kohchi, T.** (2015). Development of Gateway binary vector series with four different selection markers for the liverwort *Marchantia polymorpha*. PLoS ONE **10**: e0138876.

**Ishizaki, K., Nishihama, R., Yamato, K.T., and Kohchi, T.** (2016). Molecular genetic tools and techniques for *Marchantia polymorpha* research. Plant and Cell Physiology **57**: 262–270.

**Nelson, D.R.** (2006). Cytochrome P450 Nomenclature, 2004. In Cytochrome P450 Protocols, I.R. Phillips and E.A. Shephard, eds, Methods in Molecular Biology. (Humana Press: Totowa, NJ), pp. 1–10.

**Sugano, S.S., Nishihama, R., Shirakawa, M., Takagi, J., Matsuda, Y., Ishida, S., Shimada, T., Hara-Nishimura, I., Osakabe, K., and Kohchi, T.** (2018). Efficient CRISPR/Cas9-based genome editing and its application to conditional genetic analysis in Marchantia polymorpha. PLoS ONE **13**: e0205117.

**Yamaoka, S. et al.** (2018). Generative Cell Specification Requires Transcription Factors Evolutionarily Conserved in Land Plants. Current Biology **28**: 479-486.e5.
